# Differential histone tail citrullination by protein arginine deiminases observed via NMR spectroscopy

**DOI:** 10.64898/2026.05.01.722238

**Authors:** Alex J. Kowalczyk, Emma A. Morrison

## Abstract

Citrullination is a charge-modifying post-translational modification whereby proteinogenic arginine is converted to the non-coded amino acid citrulline by calcium-activated protein arginine deiminases (PADs; EC 3.5.3.15). The five known PAD enzymes in humans (PADs 1, 2, 3, 4, and 6) are differentially expressed and have distinct protein targets, including histones. While some PAD histone citrullination sites are known, a comprehensive investigation of all histone tail arginines targeted by catalytically active PADs 1-4 is lacking. Here, we sought to identify PAD citrullination sites in histone tails, within both histone peptides and reconstituted nucleosomes. Toward this objective, we utilized a real-time ^1^H-^15^N NMR spectroscopy-based assay. By monitoring both arginine and citrulline backbone amide peak intensities over time, we identified sites of citrullination in ^15^N-labeled histone tails within peptides and reconstituted nucleosome core particles. We found that PADs 1, 2, and 4 citrullinate all directly observable histone tail arginines to varying degrees. This is distinct from PAD3, which only moderately citrullinates H2A and H4 arginine residues and does not modify H3 tail arginines. Together, these data suggest a level of histone arginine specificity by each PAD. Furthermore, histone tail citrullination is altered within nucleosomes as compared to isolated peptides, which we interpret as reflecting differences in conformation and accessibility. We also speculate that citrullination increases nucleosomal histone tail dynamics, with implications for crosstalk between sites of histone citrullination and other important sites of regulation by PTMs (including lysines) within and between tails.

## Introduction

Precise and responsive gene expression requires dynamic regulation of chromatin. While regulation occurs across multiple length scales, at the foundational level of chromatin is the nucleosome core particle (NCP). The canonical NCP consists of ∼147 base pairs of DNA wrapped around a histone octamer containing two each of histones H2A, H2B, H3, and H4 (1, 2) (**Fig. 1A**). The core of the histone octamer is structured but also contains basic N-terminal (as well as C-terminal for H2A) intrinsically disordered regions (IDRs), referred to as histone tails, which mediate interactions with chromatin-associating proteins along with intra- and inter-nucleosomal interactions (3–7) (**Fig. 1B**). These histone tails contain a plethora of sites for post-translational modifications (PTMs) that ultimately lead to downstream transcriptional changes (8). PTM-dependent transcriptional changes are caused by indirect effects, through the recruitment of transcriptional regulatory, chromatin remodeling, and architectural complexes via their reader domains, and direct effects, by disrupting histone-DNA interactions to alter chromatin organization and increase accessibility to transcriptional machinery (9–11).

**Figure 1.**
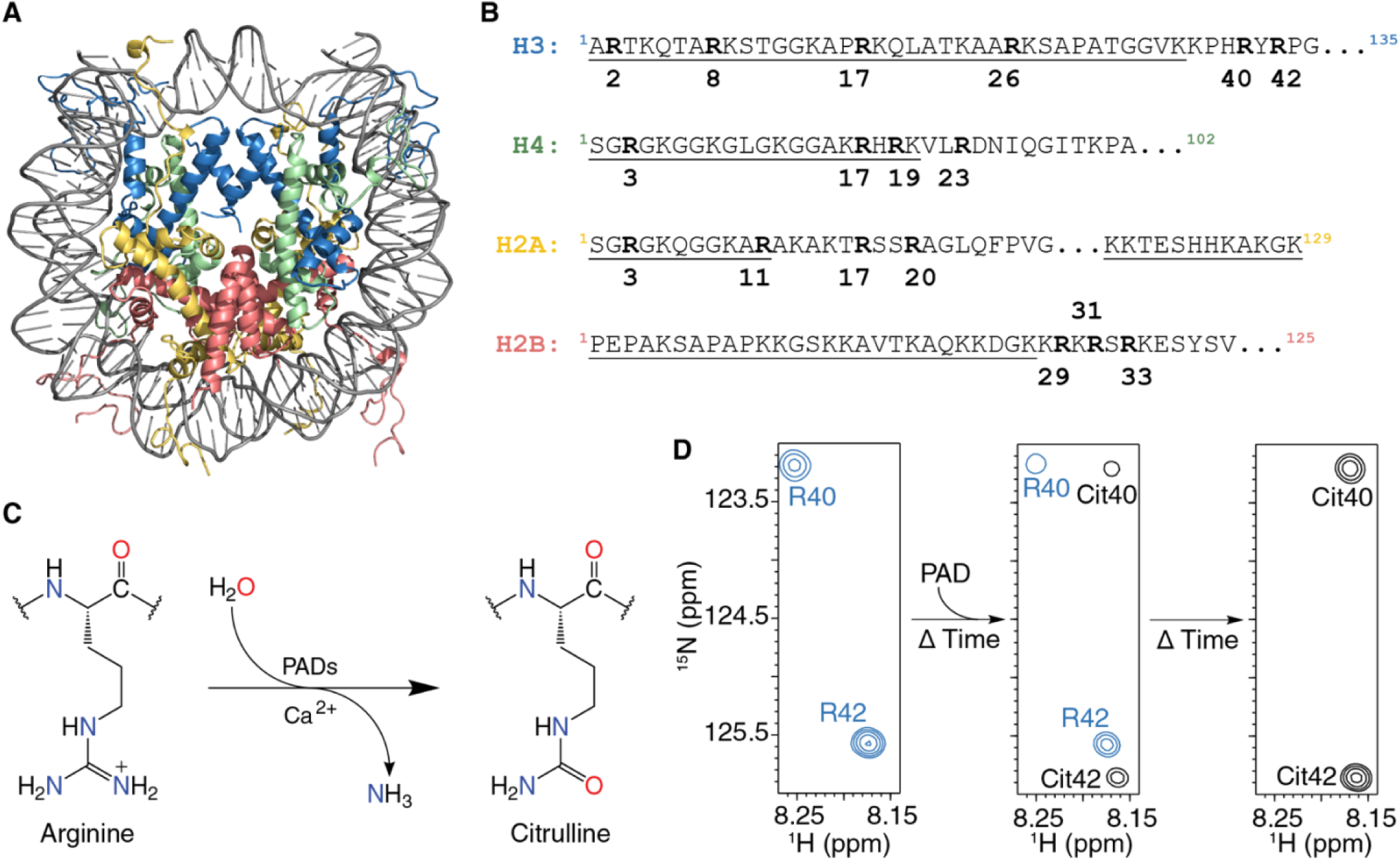
Utilizing real-time ^1^H-^15^N NMR spectroscopy to identify sites of citrullination by each PAD isozyme. (A) Structure of the nucleosome core particle (NCP) consisting of a histone octamer wrapped in 147 base-pairs of Widom 601 DNA (97). (B) Partial sequences of histones H2A, H2B, H3, and H4 with arginines in bold and the intrinsically disordered tail regions underlined. (C) Schematic depicting PAD-catalyzed citrullination of protein arginine residues. (D) Overview of the NMR-based citrullination assay. An initial spectrum is acquired of the ^15^N-labeled unmodified substrate, specifically H3(1-44) with R40 and R42 shown in this example. PAD isozyme is then added to the substrate, and spectra are acquired periodically as the reaction progresses. If an arginine is modified, the peak decays as a citrulline peak emerges.

Deimination of proteinogenic arginine, also known as citrullination, is a charge-altering PTM occurring on both the histone tails and core. Citrullination is implicated in chromatin organization yet remains understudied compared to other epigenetic modifications. Notably, PAD4-mediated citrullination occurs in neutrophils, leading to chromatin decondensation that can contribute to either highly controlled extracellular trap formation (NETosis) or dysregulated global chromatin decondensation and cell death known as Leukotoxic Hypercitrullination (12–17). Additionally, PAD2 citrullination has been associated with chromatin decondensation at genes downstream of estrogen receptor alpha (18–21). Despite citrullination being implicated in gene regulation, there has been only one proposed candidate effector (“reader”) protein for citrulline (22), standing in contrast to well-characterized epigenetic modifications that have a variety of associated reader proteins, such as lysine acetylation and methylation (5, 23). Furthermore, no known enzymes catalyze the reverse reaction to convert citrulline residues back to arginine (“erasers”), suggesting that the modification is irreversible (24). Together, these studies support a direct role of citrullination in chromatin organization and dynamics.

Charge-neutralizing modifications to arginine residues are predicted to alter protein-protein (25–27) and protein-DNA interactions (28), thereby affecting higher-order chromatin organization. Citrullination occurs by reacting the positively charged guanidium group of the arginine sidechain with water to form a neutral ureido functional group. The reaction produces the non-encoded amino acid citrulline and releases ammonia as a byproduct (**Fig. 1C**). Citrullination is catalyzed by a class of calcium-dependent enzymes known as peptidyl- or protein-arginine deiminases (PADs) (EC 3.5.3.15), which are largely believed to require a supraphysiological concentration of calcium for activation (29–31). There are five known isozymes within this enzyme class, PADs 1, 2, 3, 4, and 6, of which all but PAD6 have documented catalytic activity (32–40). The PAD enzymes are differentially expressed, both temporally and spatially, and have distinct protein targets, thus making the isozymes non-redundant (41–43). Although histones are a prominent category of PAD substrates, and histone citrullination leads to altered gene regulation, the epigenetic function of citrullination remains incompletely understood (20, 21, 44–49).

To identify histone citrullination sites, a variety of techniques have been employed, including biochemical assays, antibodies, and mass spectrometry. Prior studies have utilized both full-length and short peptide sequences of histones in colorimetric biochemical assays to assess kinetic activities of the PADs and identify sites of citrullination (33–37). The assay relies on reacting the ureido functional group of citrulline in a highly acidic environment to produce a measurable chromophore (50). While this assay detects whether citrulline is present within a substrate, it is unable to identify individual sites of modification. Most histone tails contain multiple arginines (**Fig. 1B**), so studying individual sites would require mutations that could alter PAD binding and enzymatic activity. Similarly, antibody-based detection suffers from a lack of site-specific resolution. Commercially available anti-citrullinated-H3 antibodies do not differentiate between individual histone positions unless specifically generated to do so, as in Guertin et al (21). Mass spectrometry detection of citrullination also has complicating factors. Citrullination leads to a small mass shift of +0.98 Da, which is difficult to distinguish from deamidation of nearby asparagine and/or glutamine residues (51) or incorporation of natural abundance ^13^C or ^15^N (52). While methods exist to enrich for citrulline prior to mass spectrometry, larger amounts of product are needed for detection (53–57). These approaches have improved our understanding of histone citrullination, yet a comprehensive assessment of histone tail citrullination at site-specific resolution by each catalytically competent PAD is lacking.

As the intrinsically disordered histone tails and their modifications play an important role in dynamic chromatin regulation (3, 4, 6, 58–60), we sought to identify individual histone tail arginine targets of each catalytically active PAD and understand how the native nucleosomal context affects citrullination activity. To this end, we utilized a real-time ^1^H-^15^N NMR-based kinetic assay to observe site-specific citrullination of the histone tails in the context of either peptides or reconstituted NCPs and simultaneously tracked both arginine decay and citrulline growth (**Fig. 1D**). We find that PADs 1, 2, and 4 largely citrullinate all histone tail arginines at varying observed completion times; in contrast, PAD3 distinctly does not citrullinate H3 tail arginines. Furthermore, widespread increases in peak intensity upon nucleosomal citrullination suggest an increase in histone tail mobility, likely due to decreased tail-DNA interactions. We speculate that citrullination increases subsequent enzymatic modification via histone PTM crosstalk.

## Methods

### Histone tail peptide purification

Histone peptides H3(1-44) (<u>ARTKQTARKS</u> <u>TGGKAPRKQL</u> <u>ATKAARKSAP</u> <u>ATGGVK</u>KPHR YRPG) and H2B(1-39) (<u>PEPAKSAPAP</u> <u>KKGSKKAVTK</u> <u>AQKKDGK</u>KRK RSRKESYSV) were expressed from pET3a vectors encoding for the N-terminal tail residues (underlined) and extending towards the core to include a native tyrosine for concentration determination. Histone tail peptides H2A(1-28) F25Y (<u>SGRGKQGGKA</u> <u>R</u>AKAKTRSSR AGLQYPVG) and H4(1-33) T30Y (<u>SGRGKGGKGL</u> <u>GKGGAKRHRK</u> VLRDNIQGIY KPA), both containing non-native tyrosine mutations for concentration determination, were expressed from plasmids encoding for N-terminal His-SUMO tags to minimize degradation and aid in purification. H3(1-44), H4(1-33) T30Y, and H2A(1-28) F25Y were expressed from BL21 (DE3) (New England BioLabs) and H2B(1-39) was expressed from Rosetta2 (DE3) pLysS chemically competent *E. coli* (Novagen). Expression of isotopically labeled histone tail peptides was induced with IPTG (0.3 mM final concentration) after growing transformed *E. coli* to an OD600 of ∼1.0 in M9 minimal media with ^15^N NH_4_Cl (1 g/L) and harvested three hours after induction. Histone tail peptides were extracted by sonicating cell pellets on ice in histone lysis buffer (50 mM Tris-HCl pH 7.5, 100 mM NaCl, and 2 mM EDTA containing 1 mM benzamidine HCl, 2-mercaptoethanol, 0.5% Triton X-100, DNase (Sigma), protease inhibitor cocktail tablets (Roche) as additives) and the lysate clarified via centrifugation for 30 minutes at 4°C and 17,600 rcf. Untagged histone tail peptides (H3 and H2B) were purified by running the soluble lysate over a benchtop gravity column containing a 5-mL bed of equilibrated Q-Sepharose (Cytiva) followed by a 10-mL wash with histone lysis buffer without additives. Tagged peptides (H2A and H4) were run over a nickel affinity gravity-flow column (Ni-Penta agarose resin, Marvelgent), eluted with imidazole, and dialyzed overnight to remove imidazole. The following day, His-SUMO tags were cleaved with 1:100 ULP1 SUMO protease for four hours and removed by running over a second nickel affinity gravity - flow column, resulting in scarless histone tails. The column eluates were combined, 0.45-μm syringe-filtered, and purified by Source 15S (Cytiva) cation exchange FPLC. Fractions containing histone tail peptides were combined, lyophilized, resuspended in water, and 0.22-μm syringe-filtered before further purification by PROTO 300 C18 10 µm (Higgins Analytical) HPLC (Shimadzu). Fractions containing purified histone tail peptides were successively lyophilized and resuspended in water three times to remove acetonitrile and trifluoroacetic acid prior to experimentation. Purified histone tail peptides were confirmed by ESI mass spectrometry using a Finnigan LTQ (Thermo Scientific) and were aliquoted into stocks after determining the concentration using a Nanodrop (Thermo Scientific) with a calculated extinction coefficient of 1490 M^-1^cm^-1^ (determined with ExPASy ProtParam) at 280 nm.

### NCP preparation

NCPs were prepared with the following human histone sequences: H2A (UniProt P0C0S8), H2B (UniProt P62807), H3 (UniProt Q71DI3, except C110A), H4 (UniProt P62805), and 147 base pairs of the Widom 601 DNA (5’-ATCGAGAATC CCGGTGCCGA GGCCGCTCAA TTGGTCGTAG ACAGCTCTAG CACCGCTTAA ACGCACGTAC GCGCTGTCCC CCGCGTTTTA ACCGCCAAGG GGATTACTCCC TAGTCTCCAG GCACGTGTC AGATATATAC ATCCGAT-3’). Histones were expressed and purified as previously described (61). Unlabeled histones were grown in LB, while ^15^N-labeled histones were grown in M9 minimal media with ^15^N NH_4_Cl (1g/L). NCPs were assembled as previously described (62, 63). Briefly, the H2A/H2B dimer and H3/H4 tetramer were separately refolded by dialyzing from denaturing conditions into 2 M KCl. Refolded components were purified by size-exclusion chromatography. ^15^N NCPs were reconstituted by mixing 1:2:1 (molar ratio) DNA:dimer:tetramer with only a single histone component ^15^N-labeled in a given sample. NCPs were formed by desalting via exponential gradient dialysis. Reconstituted NCPs were further purified by a 10-40% sucrose gradient to remove free DNA, subnucleosomal species, and aggregates. NCP quality was confirmed by both 5% native PAGE and 18% SDS-PAGE (**Fig. S1**).

### PAD preparation

PADs 1 (UniProt Q9ULC6), 2 (UniProt Q9Y2J8), and 3 (UniProt Q9ULW8) were recombinantly expressed with an N-terminal 10xHis-tag in BL21(DE3) *E. coli* (New England BioLabs). PAD4 (UniProt Q9UM07, except containing wild-type residues S55, A82, and A112 (64, 65)) was recombinantly expressed with an N-terminal GST tag in Rosetta2 (DE3) pLysS *E. coli* (Novagen). Transformed bacteria in LB media were induced at an OD600 of ∼0.6-0.7 with 50 µM (PADs 1-3) or 25 µM (PAD4) IPTG for overnight expression (∼16 hours). Cells were lysed by sonication, and the lysate was clarified by centrifugation. His-tagged PADs were purified over a nickel affinity gravity-flow column and eluted with imidazole. GST-PAD4 was purified over a glutathione gravity-flow column and cleaved off the beads with PreScission Protease. PADs were further purified by Source 15Q anion exchange and S200 size-exclusion chromatography into buffer containing 50 mM Tris-HCl pH 7.6, 10% glycerol (w/v), 400 mM NaCl, 0.5 mM TCEP, 0.1 mM EDTA. Fractions containing pure PAD were concentrated to 35-75 µM by benchtop centrifugation in 10k MWCO concentrators, and final PAD stocks were stored at -80°C. A single PAD prep for each isozyme was used across all assays conducted with peptide and NCP substrates shown in the main text figures. PAD stocks were uniformly diluted to 33 µM for the NMR-based assays.

PAD activity was assessed using a standard colorimetric assay with Nα-benzoyl-L-arginine ethyl ester (BAEE) for the test substrate (50) (see **Supplemental Methods, Fig. S2**). Activities of the PAD preps used in this paper to BAEE are consistent with prior studies (32, 33, 36).

### NMR histone tail assignments

Backbone peak assignments were obtained on ^13^C/^15^N-isotopically labeled histone tail peptides by acquiring HNCACB, HN(CO)CACB, and HNCA spectra with intervening ^1^H-^15^N HSQCs before and after each triple resonance experiment to confirm sample integrity. Triple resonance experiments were collected with 2048 (^1^H) x 128 (^13^C) x 50 (^15^N) total points, while the HSQC was collected with 1024 (^1^H) x 300 (^15^N) total points. All NMR assignment experiments were collected with an acquisition time of 102 (^1^H) and 22 (^15^N) ms, while both the HNCACB and HN(CO)CACB were collected with 5 ms (^13^C) and the HNCA was collected with 10 ms (^13^C). Each experiment was collected with a spectral width of 16.7 (^1^H) and 18.5 (^15^N) ppm. Additionally, the triple-resonance spectral widths included 41.4 ppm (^13^C) for the HNCA and 82.9 ppm (^13^C) for the HNCACB and HN(CO)CACB. The ^13^C /^15^N-H4(1-33) T30Y and H2A(1-28) F25Y data were collected with 4 scans each for the HNCACB and HN(CO)CACB and 2 scans each for the HNCA and HSQC. The ^13^C/^15^N-H2B(1-39) and H3(1-44) were collected with 8 scans each for the HNCACB and HN(CO)CACB experiments, and 4 scans each for the HNCA and HSQC experiments. Data were collected on ^13^C /^15^N-H2A(1-28) F25Y, H2B(1-39), H3(1-44), and H4(1-33) T30Y at concentrations of 2.0 mM, 1.0 mM, 0.5 mM, and 2.0 mM, respectively, in 20 mM MOPS pH 7.0, 100 mM KCl, 2 mM CaCl_2_, 0.5 mM TCEP, 0.1 mM EDTA, 5% D_2_O. To acquire backbone peak assignments for citrullinated residues, PAD1 was first added to each ^13^C/^15^N-isotopically labeled histone tail peptide and allowed to proceed for >72 hours at 4°C. Data on citrullinated histone tail peptides were collected as outlined above. All backbone peak assignment spectra were collected on a 600 MHz Bruker Avance spectrometer at 10°C. Assignments on the ^15^N-H3-NCP and ^15^N-H2A/H2B-NCPs were transferred from Morrison et al. and Ohtomo et al., respectively (7, 66). Assignment of the ^15^N-H4-NCP was conducted by transferring assignments from Zhou et al. (59) in addition to comparison with our H4(1-33) T30Y assignments due to differences in H4 sequences (i.e., Drosophila vs. human). Full tail peptide and NCP spectra, in addition to their PAD1 citrullinated end point, can be viewed in **Fig. S3**.

### NMR-based citrullination assay

Citrullination assays were conducted using ^1^H-^15^N SOFAST-HMQC experiments on a Bruker AVANCE NEO 800 MHz NMR Spectrometer with a cryoprobe. HMQCs were collected with 24 scans and 2048 (^1^H) x 176 (^15^N) total points for histone tail peptide assays. For NCP assays, HMQCs were collected with 40 scans and 1024 (^1^H) x 176 (^15^N) total points. HMQC acquisition times were 82 and 49 milliseconds (^1^H) for histone tail peptides and NCP assays respectively, and 49 ms (^15^N) for both assays. Spectral widths were 15.6 and 13.0 ppm (^1^H) for histone tail peptide and NCP assays, respectively, and 22.0 ppm (^15^N). Total time for HMQC data collection was 22 minutes 39 seconds and 22 minutes 00 seconds for histone tail peptide and NCP assays, respectively. For histone tail substrates, data were collected at 10°C with 100 μM ^15^N-labeled tails in 20 mM MOPS pH 7.0, 100 mM KCl, 2 mM CaCl_2_, 0.5 mM TCEP, 0.1 mM EDTA, 5% D_2_O. Data for nucleosome substrates were collected at 25°C with 50 μM ^15^N-labeled NCPs (meaning, 100 μM ^15^N-labeled histone) in the same buffer. After a baseline spectrum of the substrate was collected (spectrum 1, t=0), the sample was removed from the spectrometer, and the reaction was initiated by addition of 2 µL of 33 µM PAD isozyme to the substrate (final concentration of 200 nM enzyme, or 1:500 enzyme-to-substrate molar ratio). Timing of the experiment began upon PAD addition. Dead time between sample mixing and the start of the second HMQC was 15-21 and 14-17 minutes for tail peptide and NCP assays, respectively. After the sample was returned to the spectrometer, 19 spectra were queued to collect data over nearly 16 hours, with increasing time delays (18 spectra in the case of PAD1-H2A(1-28) F25Y). Spectra 2-5 were queued and run consecutively with no time delay (spectra 2-4 for PAD1-H2A(1-28) F25Y). Spectra 6-7, 8-9, and 10-20 were queued with 10-, 20-, and 40-minute time delays, respectively (spectra 5-6, 7-8, 9-19 for PAD1-H2A(1-28) F25Y). The time used in subsequent kinetic analyses accounts for the exact dead time of the given experiment. Data were collected as an n=1. For additional assays conducted at different conditions, see **Supplemental Methods, Supplemental Results, Figs. S4-S7.**

### NMR data analysis

All collected NMR data were transferred into NMRbox for processing and analysis (67). Processing was conducted using NMRPipe (68) and subsequently analyzed in CcpNmr Analysis v2.5.2 (69). Peak intensities were scaled down and plotted in Igor Pro 9.05.

Progress curve intensities were fit to the following functions to estimate the time for 50% completion (t_50%_): linear

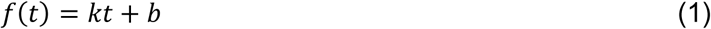

where *k* is the slope of the line, *t* is time, and *b* is the y-intercept; exponential

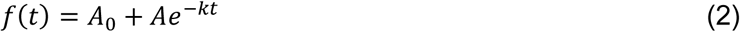

where *A*_0_ is the initial amplitude, *A* is the asymptotic amplitude, *k* is the rate, and *t* is time; or Gompertz sigmoidal

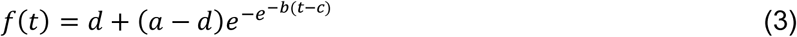

where *a* and *d* are horizontal asymptotes, *b* is the growth parameter, *c* is the inflection point of the curve, and *t* is time. The residual sum of squares (RSS) was utilized to determine best fits to the data in an unbiased manner (see **Table S1**, **Figs. S8-S14**). For completed reactions, functions were fit unconstrained. For incomplete reactions, data fits were performed while holding the amplitude at an estimated completion intensity based on the corresponding completed PAD1 reaction, adjusted to account for intensity differences due to slight variations in substrate concentration: The estimated completion intensity for a given residue was calculated from the PAD1 intensity (either minimum intensity for arginine, or maximum intensity for citrulline) multiplied by a correction factor, calculated as the average of the initial intensities of a given substrate used in PAD 2, 3, and 4 reactions divided by the corresponding PAD1 initial intensity. To estimate the t_50%_ for a given reaction, the intensity at 50% completion (*I*_50%_) was first calculated with the function

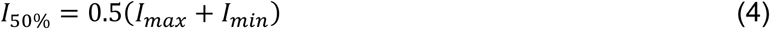

where *I_max_* is the maximum intensity and *I_min_* is the minimum intensity of a given residue. The acquired *I*_50%_ value was then fed back into the selected fit equation with the corresponding fit parameters (i.e., one of Eqs. 1-3) to acquire a time for 50% completion for a given residue. For a comparison of main text data compared to supplementary data fits, see **Fig. S15**. For utilizing adjacent (i+1) residues data to estimate a t_50%_, see **Fig. S16**.

## Results

### NMR-based assay to site-specifically monitor histone citrullination

The NMR-based assay design enables simultaneous site-specific detection of all visible arginines in the native sequence of a given histone substrate while tracking both arginine decay and citrulline accumulation (**Figs. 1D**, **2**). Prior studies have demonstrated the successful application of NMR to the understanding of enzymatic PTM sites and rates for both histone and non-histone substrates (6, 70–79). To conduct a comprehensive investigation of histone tail arginine targets of the four catalytically active PAD isozymes, we collected ^1^H-^15^N SOFAST-HMQC spectra of ^15^N-isotopically labeled substrates as citrullination reactions progressed (**Fig. 1B, 1D**). We acquired sequential ^1^H-^15^N SOFAST-HMQC spectra (∼22 minutes in length) following PAD addition, with increasing intervening delays introduced after ∼1.75 hours, for a total experimental time of ∼16 hours post PAD-addition (see **Methods** for details).

**Figure 2.**
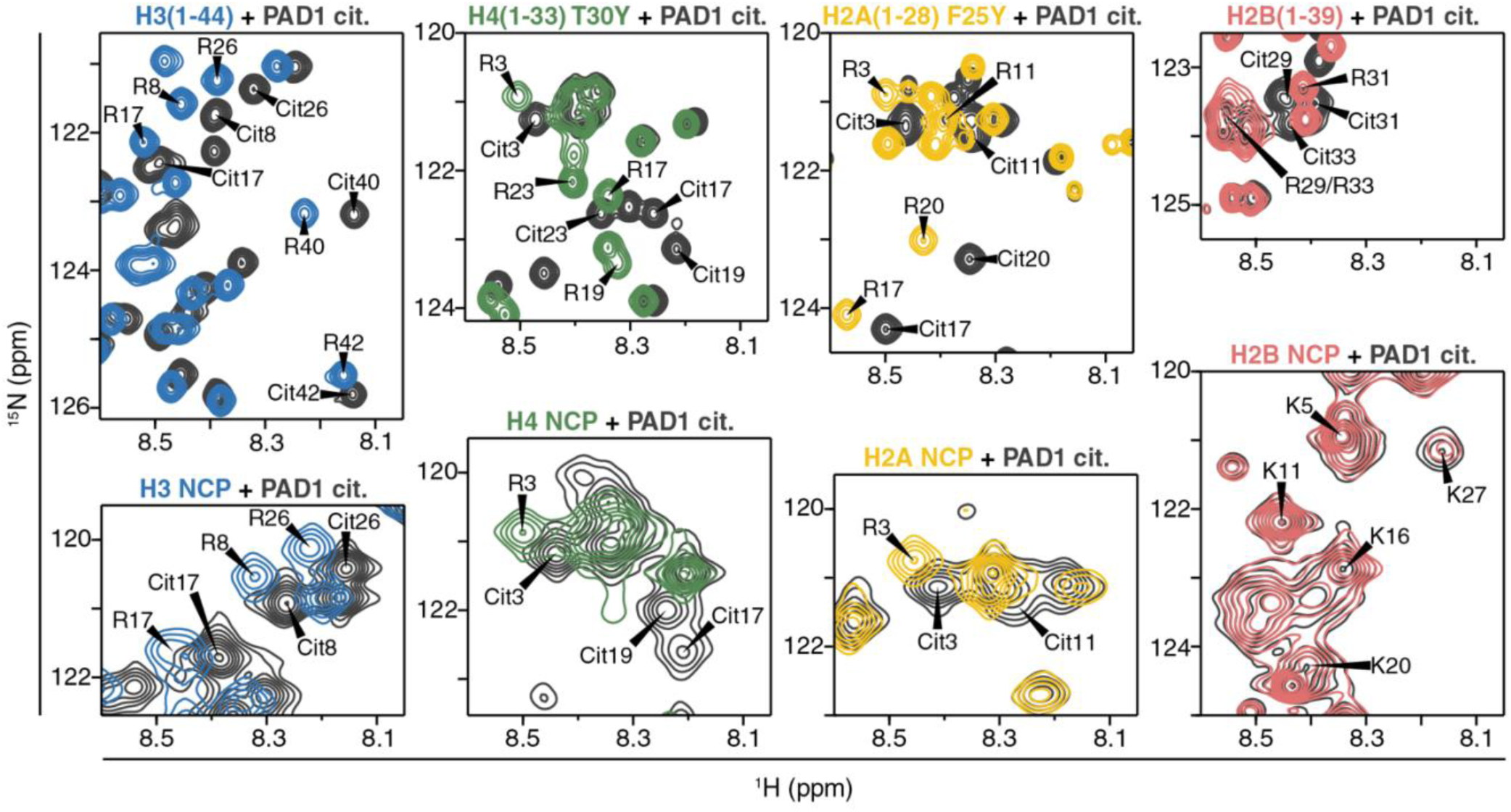
Spectra of arginines and citrullines in ^15^N-labeled histone tail peptides and reconstituted NCPs by PAD1. Top, left to right: ^1^H-^15^N SOFAST-HMQC spectra of ^15^N-labeled histone tail peptides of H3 (blue), H4 (green), H2A (yellow), and H2B (red) with their PAD1-citrullinated end-states in dark grey. Bottom, left to right: spectra of NCPs reconstituted with ^15^N-labeled histone H3 (blue), H4 (green), H2A (yellow), and H2B (red) with their PAD1-citrullinated end-states in dark grey. Spectra were acquired on either 100 µM tail peptide or 50 µM NCP (equivalent to 100 µM ^15^N-labeled histone) on an 800 MHz Bruker NMR spectrometer at 10°C (tail peptide) or 25°C (NCP) in 20 mM MOPS pH 7.0, 100 mM KCl, 2 mM CaCl_2_, 0.5 mM TCEP, 0.1 mM EDTA, and 5% D_2_O. Full spectra are shown in **Fig. S3**.

To study citrullination of histone tails within peptide and nucleosomal contexts, the substrates used in this study were ^15^N-labeled histone peptides and NCPs reconstituted with only one histone (H3, H4, H2A, or H2B) ^15^N-labeled within a given sample. Meaning, when collecting spectra on a given ^15^N-histone-labeled NCP, all histone arginines are present in the sample, but we are only monitoring the arginines and citrullines of the ^15^N-labeled histone. Furthermore, only residues of the dynamic histone tails are visible within ^1^H-^15^N spectra of a ^15^N-histone-labeled NCP due to the large size of the particle (∼200 kDa) (6, 7, 59). Differential spectral quality for peptide and nucleosomal substrates necessitated running the experiments at 10°C and 25°C, respectively. This complicates kinetic comparison of histone tails within peptides and NCP, so we limit our comparisons to qualitative observations of trends.

To enable a metric of comparison between histone arginine positions and PAD isoforms, progress curves of arginine decay and citrulline growth were fit to determine the time to reach 50% completion (t_50%_) (see **Methods** for details). For sequential reactions and simultaneous reactions that occur at different rates, the curve shape of positions modified later deviates from the exponential curve expected when only a single position is hit (or when a position is hit first or much faster than other positions) (71, 77). Due to differences observed in the shapes of progress curves across PAD-substrate combinations, multiple functions (linear, exponential, or sigmoidal) were used to fit the data and calculate t_50%_ values, similar to the approach used in He, et al (77). Although arginine and citrulline t_50%_ values at a given position should match, factors such as peak overlap, effects from modification of adjacent residues, and changes in dynamics may influence the progress curves, leading to differences in t_50%_ observed between some pairs of arginine and citrulline.

To demonstrate reproducibility, two datasets are presented for histone peptide-PAD combinations. The data displayed in main text figures (**Figs. 3, 4, 6, 8, 10**) were collected at 100 µM peptide, 1:500 enzyme-to-substrate molar ratio, and 2 mM Ca^2+^ while the data displayed in supplemental figures (**Figs. S4-S7, S15**) were collected at 50 µM peptide, 1:250 enzyme-to-substrate molar ratio, and 10 mM Ca^2+^ (see **Methods** and **Supplemental Methods** for additional details). Trends largely hold between the two data sets (see also **Supplemental Methods**, **Supplemental Results**).

**Figure 3.**
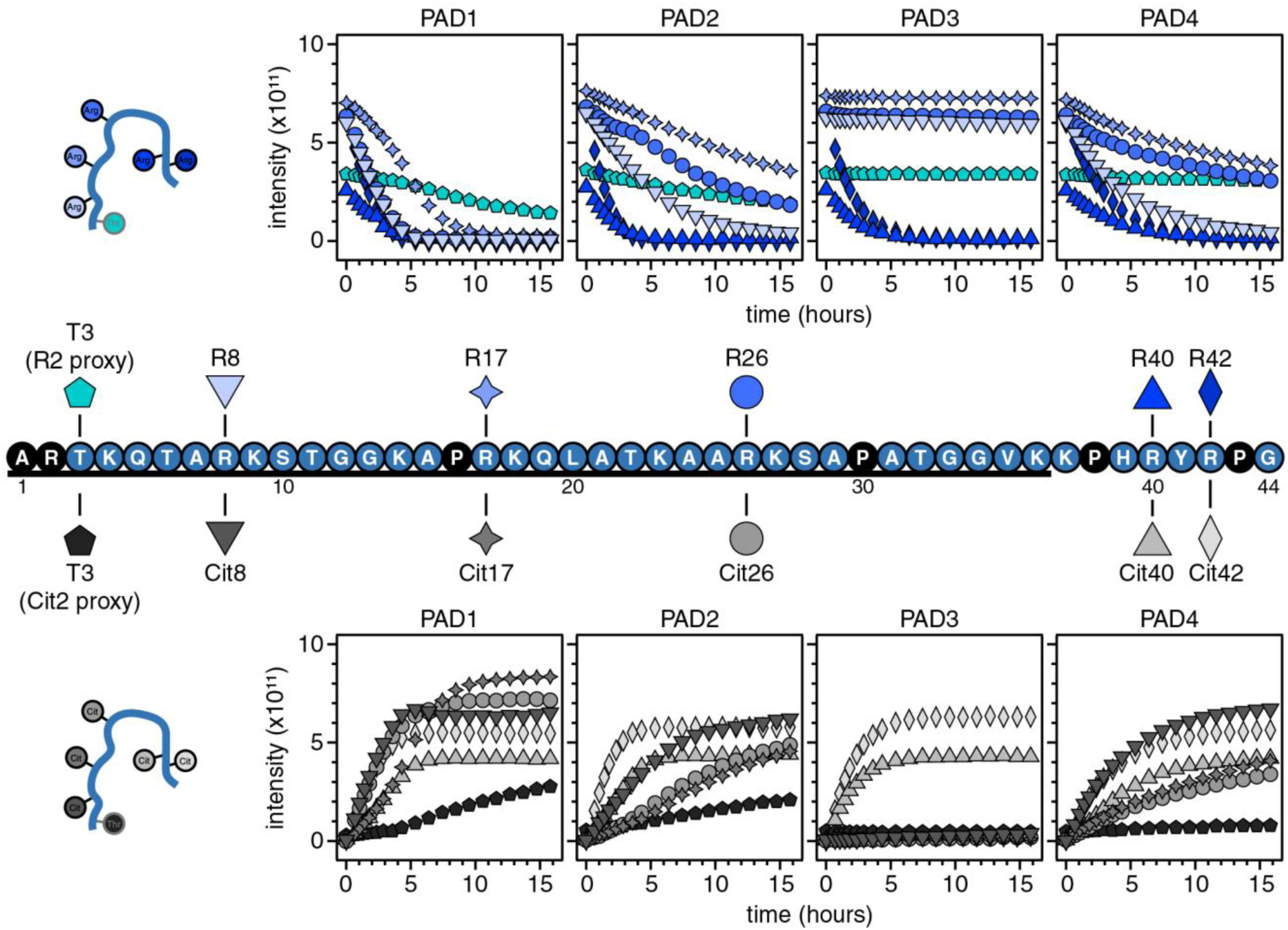
Progress curves of H3 peptide arginines and citrullines demonstrate site-specific differences in PADs 1, 2, and 4 versus PAD3. Top: Cartoon of unmodified H3 peptide (left) and ^15^N-H3 peptide arginine progress curves for PADs 1, 2, 3, 4 (right). Arginine progress curves are denoted by the shapes found above the H3(1-44) peptide sequence (middle). Black and blue residues represent unobserved and observed residues, respectively. Bottom: Cartoon of citrullinated H3 tail peptide (left) and graphs of ^15^N-H3 peptide citrulline progress curves for PADs 1, 2, 3, 4 (right). Citrulline progress curves are denoted by the shapes found immediately below the H3 peptide sequence. Data were collected with 100 µM ^15^N-H3(1-44) on an 800 MHz Bruker NMR Spectrometer at 10°C in 20 mM MOPS pH 7.0, 100 mM KCl, 2 mM CaCl_2_, 0.5 mM TCEP, 0.1 mM EDTA, and 5% D_2_O. PADs were added to a final concentration of 200 nM. Data were collected as n=1. For additional histone H3 tail peptide assays conducted at different conditions, see **Supplemental Methods, Supplemental Results, Fig. S4.**

**Figure 4.**
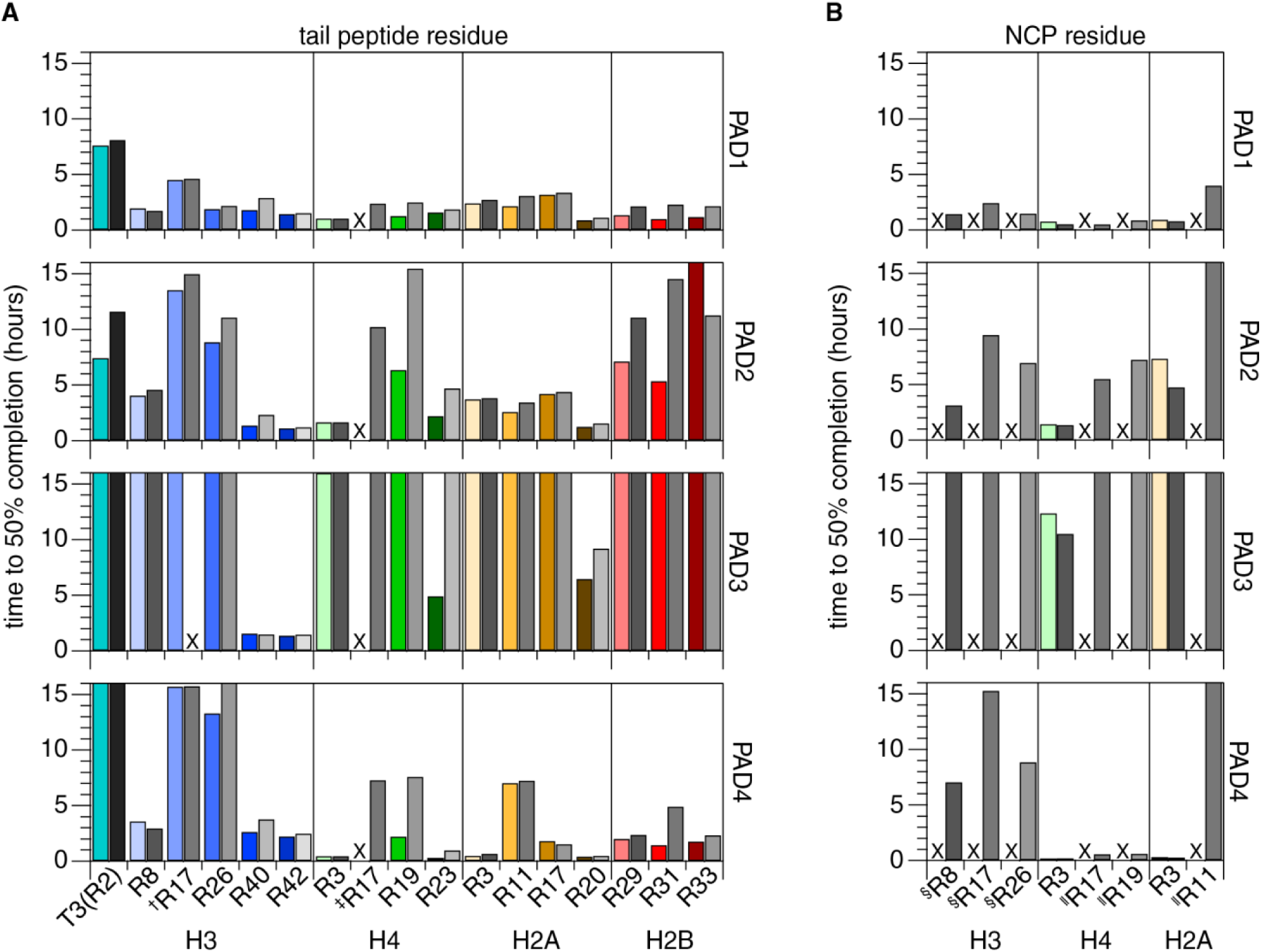
Assessing PAD site-specificity by determining time to 50% completion in the context of histone peptides and NCP. Time to 50% completion (t_50%_) graphs for PADs 1, 2, 3, and 4 (top to bottom) with each of the histone peptide (**A**) or NCP (**B**) residues. Arginines (left bar) and citrullines (right bar) are in colors and greyscale, respectively, as found in the progress curve graphs in **Figs. 3, 5-10**. “X” indicates that a t_50%_ was not determined for the following reasons: ^†^For H3 peptide with PAD3, the Cit17 t_50%_ was calculated to be a negative time value due to a minor decrease in intensity. ^‡^H4 tail peptide R17 could not be fit for any of the PADs due to severe overlap with Cit23. ^§^H3 R8/17/26 in NCP could not be fit reliably due to non-monotonic progress curves. ^‖^Residues R17/19 in ^15^N-H4-NCP and R11 in ^15^N-H2A-NCP were unobserved in the initial spectra and thus unable to be fit for calculations. All data were collected on an 800 MHz Bruker NMR spectrometer in 20 mM MOPS pH 7.0, 100 mM KCl, 2 mM CaCl_2_, 0.5 mM TCEP, 0.1 mM EDTA, and 5% D_2_O. Experiments with histone peptide substrate were run at 10°C with 100 µM ^15^N-labeled peptide. Experiments with NCP substrate were run at 25°C with 50 µM NCP (equivalent to 100 µM ^15^N-labeled histone). The y-axis is truncated at 16h, the length of the NMR-based assays. Any t_50%_ > 16h was deemed “very slow,” and any distinction beyond that was considered inaccurate to fit.

We acknowledge that the following comparisons are subject to the specific experimental conditions utilized in this paper and may change under different conditions, including the composition and concentrations of enzyme and substrate, pH, temperature, salt, and reducing agents.

### Evaluation of H3 tail peptide substrate: the H3 tail is citrullinated by PADs 1, 2, and 4

The histone H3 N-terminal tail has been a focus of PAD citrullination research and contains multiple known sites of modification (R2, R8, R17, and R26) (20, 80). We sought to acquire site-specific information in an H3 tail peptide extended out to G44 [referred to as H3(1-44)], which includes R40 and R42 in the “H3-latch” that passes between the two gyres of nucleosomal DNA (**Fig. 3, middle**) (81). Of the four arginines within the H3 tail, we observe H3 tail arginines R8, R17, R26, as well as R40 and R42 (**Fig. 2**). Although a peak for the R2 backbone amide proton is not observable under these conditions, we tracked changes in the adjacent residue (T3) as a proxy for R2 citrullination (**Figs. S3, S16; Supplemental Results**). This allowed monitoring citrullination of all arginines, whether directly or indirectly.

The arginine and citrulline progress curves vary across the PADs. PAD1 addition to ^15^N-labeled H3(1-44) leads to complete decay of arginine intensity (**Fig. 3, top**) and growth of citrulline intensity (**Fig. 3, bottom**) of the five directly visible arginine residues, indicating complete citrullination of R8, R17, R26, R40, and R42 by ∼9 hours (**Figs. 3, 4A**). The progress curves of T3 suggest R2 citrullination reaches completion near the end of the experiment (∼16 hours). PADs 2 and 4 also citrullinated all directly observable H3 tail peptide arginines, but only R8, R40, and R42 to completion within the experimental timeframe. T3 progress curves suggest partial R2 citrullination by PAD2 but a lack of activity by PAD4. Interestingly, PAD3 does not citrullinate any of the arginine residues within the true tail region of the H3 peptide yet fully citrullinates residues R40 and R42. Comparison across the PAD isozymes indicates that R40 and R42 are citrullinated the fastest (t_50%_ ∼1-4 hours, **Fig. 4A**, **Table S2**), while R17 is the slowest directly observable residue to be citrullinated by PADs 1, 2, and 4 (t_50%_ ∼4.5-16 hours), and R2 is only modified by PADs 1 and 2. Overall, these citrullination reactions are on a similar timescale as observed for other NMR-based kinetic studies of histone modification by methyl or acetyl transferases (6, 70, 72, 73, 78, 79).

### Evaluation of nucleosomal H3 tail substrate: augmented peak intensity suggests increased nucleosomal tail dynamics

We next asked how H3 tail citrullination occurs within the more native context of the NCP by conducting assays on ^15^N-H3-NCP. In the unmodified ^15^N-H3-NCP spectrum, residues T3 through K36 are resolved, allowing observation of R8, R17, and R26 (**Fig. S3**). Although T3 was utilized as a proxy for R2 in the H3 tail peptide assays, it was determined to be unsuitable for tracking R2 citrullination in the NCP due to partial peak overlap with the more intense, citrullinated form of S10 (**Fig. S3**). Upon PAD addition to the ^15^N-H3-NCP, PADs 1, 2, and 4 show citrullination activity at all directly observable arginine positions (R8, R17, and R26) ( **Fig. 5**), with PAD1 being the fastest overall. PAD3 fails to show a decrease in arginine peak intensity or an appearance of citrulline peaks, in agreement with a lack of modification of H3 tail arginines by PAD3 within the peptide (**Fig. 3**).

**Figure 5.**
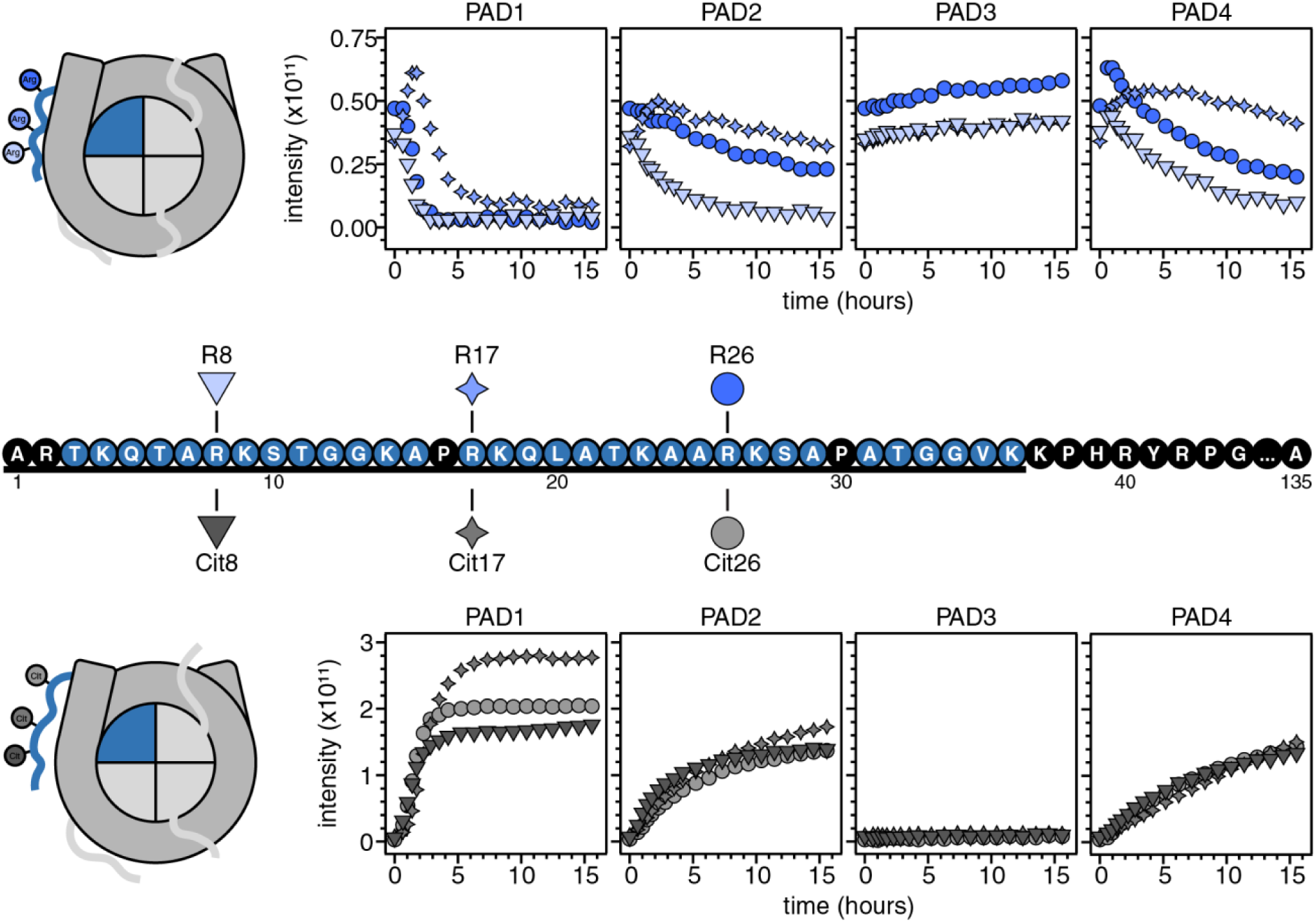
Increasing arginine intensities after PAD addition suggest citrullination affects nucleosomal H3 tail dynamics. Top: Cartoon of unmodified H3-NCP (left) and graphs of H3-NCP arginine progress curves for PADs 1, 2, 3, 4 (right). Arginine progress curves are denoted by the shapes found above the H3 sequence (middle). Black and blue residues represent unobserved and observed residues, respectively. Bottom: Cartoon of citrullinated ^15^N-H3-NCP (left) and ^15^N-H3-NCP citrulline progress curves for PADs 1, 2, 3, 4 (right). Citrulline progress curves are denoted by the shapes found immediately below the H3 sequence. Please note differences in arginine vs. citrulline y-axis scaling. Data were collected with 50 µM ^15^N-H3-NCP (equivalent to 100 µM ^15^N-labeled histone) on an 800 MHz Bruker NMR spectrometer at 25°C in 20 mM MOPS pH 7.0, 100 mM KCl, 2 mM CaCl_2_, 0.5 mM TCEP, 0.1 mM EDTA, and 5% D_2_O. PADs were added to a final concentration of 200 nM. Data were collected as n=1.

Notably, we find that a subset of H3 tail arginine-PAD isozyme combinations displays an initial increase in intensity (**Fig. 5, top**), suggesting additional process(es). To elaborate, with PAD1 and PAD2 treatment of ^15^N-H3-NCP, R17 first increases in intensity before decaying; a similar observation is made for all visible arginines (R8, R17, and R26) following PAD4 addition. Most non-arginine residues display a time-dependent increase in intensity after PAD addition for all isozymes (**Fig. S17**). These intensity increases are consistent with an increase in picosecond-to-nanosecond (ps-ns) dynamics within the tail, as previously observed with H3 tail arginine neutralization (28). We interpret the initial intensity increase in the subset of aforementioned arginines to indicate that citrullination increases H3 tail dynamics—the arginines that are modified slowly relative to other positions first report an increase in ps-ns timescale dynamics due to citrullination of other arginine(s) before decaying due to their own citrullination. The slow increase in peak intensity for R8, R17, and R26 (**Fig. 5, top**) for PAD3-treated ^15^N-H3-NCP is consistent with this model and suggests that citrullination of other residues within the NCP (either other tails or the histone core) increases H3 tail dynamics on the ps-ns timescale.

Despite the more complicated kinetic plots for the nucleosomal H3 tail arginines, the associated citrulline peaks largely display monotonic growth curves. The citrulline curves were fit to determine t_50%_ values (**Fig. 4B, Table S2**), which show similar trends across tail residues and isozymes as with peptide (**Fig. 4A,B**).

### Evaluation of H4 tail peptide substrate: the H4 tail is most effectively citrullinated by PADs 1 and 4

Next, we sought to determine which H4 tail peptide arginines are citrullinated by the PADs. Three arginine residues exist in the H4 tail: R3, R17, and R19. We extended the H4 tail peptide sequence out to residue A33 to include an additional arginine, R23, that exists towards the core of the nucleosome and added a non-native tyrosine for concentration determination [referred to as H4(1-33) T30Y] (**Fig. 6, middle**).

**Figure 6.**
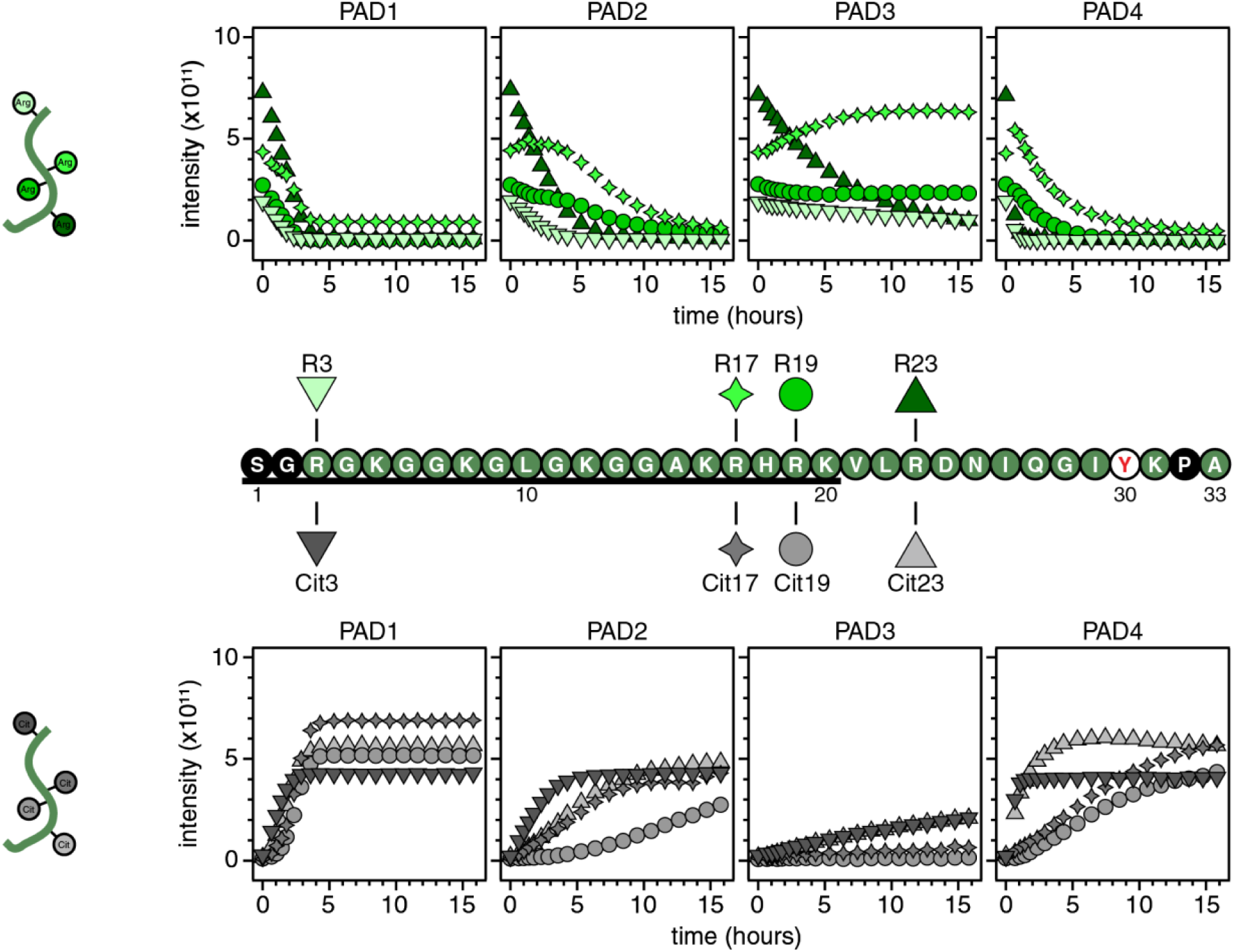
Progress curves of H4 tail peptide citrullination further highlight differences in histone modification sites by PADs 1, 2, and 4 versus PAD3. Top: Cartoon of unmodified H4 peptide (left) and graphs of ^15^N-H4 peptide arginine progress curves for PADs 1, 2, 3, 4 (right). Note that R17 intensity is influenced by the nearby growth of Cit23 (see Fig. 2, top, for H4 tail peptide arginine and citrulline chemical shifts). Arginine progress curves are denoted by the shapes found above the H4(1-33) T30Y peptide sequence (middle). Black and green residues represent unobserved and observed residues, respectively. Bottom: Cartoon of citrullinated H4 peptide (left) and graphs of H4 peptide citrulline progress curves for PADs 1, 2, 3, 4 (right). Citrulline progress curves are denoted by the shapes found immediately below the H4 peptide sequence. Data were collected with 100 µM ^15^N-H4(1-33) T30Y on an 800 MHz Bruker NMR spectrometer at 10°C in 20 mM MOPS pH 7.0, 100 mM KCl, 2 mM CaCl_2_, 0.5 mM TCEP, 0.1 mM EDTA, and 5% D_2_O. PADs were added to a final concentration of 200 nM. Data were collected as n=1. For additional histone H4 tail peptide assays conducted at different conditions, see Supplemental Methods, Supplemental Results, and Fig. S5.

Progress curves were compared. Treatment of the ^15^N-labeled H4(1-33) T30Y with PAD1 resulted in complete citrullination of all four arginines in the H4 peptide (R3, R17, R19, and R23) by 5 hours, as seen from both arginine and citrulline progress curves (**Fig. 6**). PADs 2 and 4 completely citrullinated residues R3 and R23, with slower citrullination of both R17 and R19, but PAD4 has shorter t_50%_ than PAD2 (**Fig. 4**). PAD3 citrullinates both R3 and R23, but minimal citrullination of R17 and R19 is observed. Faster citrullination of R23 as compared to R17 results in Cit23 emerging nearly on top of R17, skewing the arginine decay curves to varying degrees for each PAD and complicating interpretation. Additionally, the R19 peak may sense differential citrullination states of nearby arginines, R17 and R23, leading to peak shifting and altered intensity over time. The citrulline growth curves (**Fig. 6, bottom**) further support similar sites of H4 arginine modification by PADs 1, 2, and 4 (i.e., R3, R17, R19, and R23), in contrast to PAD3 (i.e., R3 and R23, but R17 and R19 only minimally).

### Evaluation of nucleosomal H4 tail substrate: citrullination leads to visibility of the H4 tail basic patch in the NCP

We then investigated H4 tail citrullination within the NCP. Prior to treatment with PADs, we only observe H4 tail residues R3-A15 in the unmodified ^15^N-H4-NCP (**Fig. 7, middle**). This is consistent with previous studies that only observed through A15 (59, 82). Following addition of the PADs to ^15^N-H4-NCP, we observe citrullination of R3 by all four PADs (**Fig. 7, top**), consistent with our observations of H4 R3 citrullination in the tail peptide (**Fig. 6**). As the ^15^N-H4-NCP becomes citrullinated, residues K16-K20, known as the basic patch, become observable in the spectra (**Fig. 7, middle/bottom, Figs. S3, S18**). Although R3 is the only arginine visible in the spectra, Cit3, Cit17, and Cit19 are all observable. In addition, we observe a near-global increase in intensity across the tail (**Fig. S18**). Previously, H4 K16 acetylation mimetics led to the appearance of basic patch residues, which was attributed to a disruption in the structure of the basic patch and its interactions with DNA (4, 59, 83–85). We speculate that citrullination of R3, R17, and/or R19 has similar effects, resulting in reduced H4 tail-DNA interaction, increased H4 tail dynamics, and thus resolved H4 basic patch residues in ^1^H-^15^N spectra.

**Figure 7.**
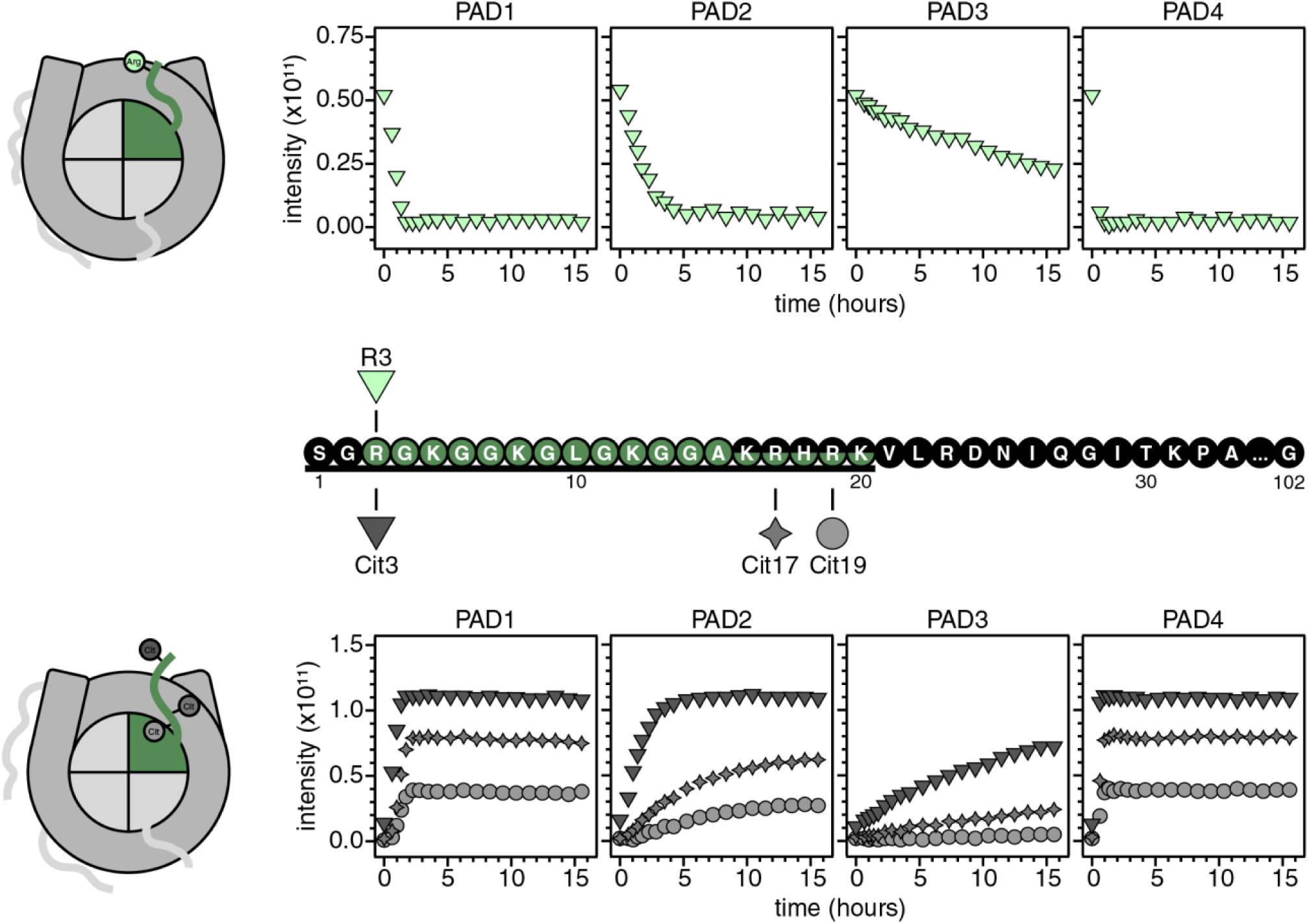
Citrullination resolves the basic patch region of the nucleosomal H4 tail. Top: Cartoon of unmodified H4-NCP (left) and progress curves of the sole observable arginine in ^15^N-H4-NCP for PADs 1, 2, 3, 4 (right). Arginine progress curves are denoted by the shapes found above the H4 sequence (middle). Black and green residues represent unobserved and observed residues, respectively. Citrullination of the NCP resolves the H4 tail from A15 to K20 (see **Fig. S3**). Bottom: Cartoon of citrullinated H4-NCP (left) and graphs of ^15^N-H4-NCP citrulline progress curves for PADs 1, 2, 3, 4 (right). Citrulline progress curves are denoted by the shapes found immediately below the H4 sequence. Please note differences in arginine vs. citrulline y-axis scaling. Data were collected with 50 µM ^15^N-H4-NCP (equivalent to 100 µM ^15^N-labeled histone) on an 800 MHz Bruker NMR spectrometer at 25°C in 20 mM MOPS pH 7.0, 100 mM KCl, 2 mM CaCl_2_, 0.5 mM TCEP, 0.1 mM EDTA, and 5% D2O. PADs were added to a final concentration of 200 nM. Data were collected as n=1.

Comparison of progress curves and t_50%_ values across nucleosomal H4 tail residues and isozymes provides insight into variation between PADs. Progress curves for PADs 1 and 4 appear similar, and R3, R17, and R19 are fully citrullinated within ∼2 hours. Values of t_50%_ suggest that PAD4 citrullinates nucleosomal H4 tail targets faster than PAD1, which is opposite the behavior observed for nucleosomal H3 tail targets. PAD2 citrullinates all three arginines, but only citrullination of R3 is complete well before the end of the experiment (t_50%_ ≈ 1.5 hours). PAD3 is slower, partially citrullinating R3 and only minimally citrullinating R17 and R19. Thus, while PADs 1 and 4 appear to simultaneously citrullinate the three tail arginines, PADs 2 and 3 citrullinate R3 substantially faster than R17 and R19. R3 is the first of the three tail arginines to be citrullinated within the peptide version, too, so the nucleosomal context does not appear to have a strong influence on the order of modification.

### Evaluation of H2A tail peptide substrate: the H2A tail is most effectively citrullinated by PAD1

Of the four canonical histones, the H2A N-terminal tail region is the smallest at 11 residues. The H2A tail contains two arginines, R3 and R11. We extended the sequence of the H2A tail peptide out to residue G28, including two additional arginines at positions 17 and 20 within the *a*1-extension helix and adding a non-native tyrosine for concentration determination [referred to as H2A(1-28) F25Y] (**Fig. 8, middle**).

**Figure 8.**
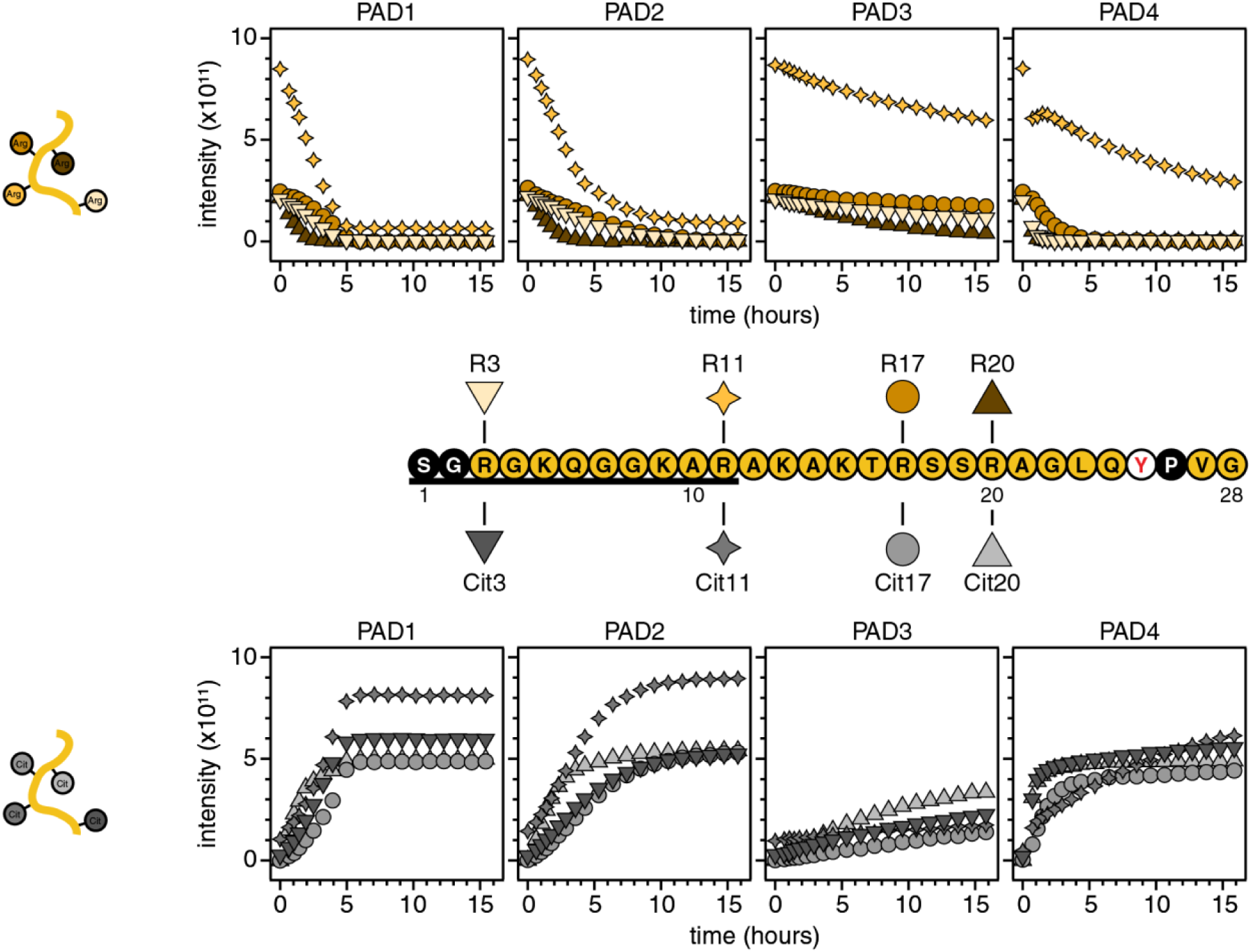
Broad citrullination of the H2A tail peptide is observed for all PADs. Top: Cartoon of unmodified H2A peptide (left) and progress curves of ^15^N-H2A peptide arginines with PADs 1, 2, 3, 4 (right). Arginine progress curves are denoted by the shapes found above the H2A(1-28) F25Y peptide sequence (middle). Black and yellow residues represent unobserved and observed residues, respectively. Bottom: Cartoon of citrullinated H2A peptide (left) and citrulline progress curve graphs of ^15^N-H2A peptide for PADs 1, 2, 3, 4 (right). Citrulline progress curves are denoted by the shapes found immediately below the H2A peptide sequence. Data were collected with 100 µM ^15^N-H2A(1-28) F25Y on an 800 MHz Bruker NMR spectrometer at 10°C in 20 mM MOPS pH 7.0, 100 mM KCl, 2 mM CaCl_2_, 0.5 mM TCEP, 0.1 mM EDTA, and 5% D_2_O. PADs were added to a final concentration of 200 nM. Data were collected as n=1. For additional histone H2A tail peptide assays conducted at different conditions, see **Supplemental Methods**, **Supplemental Results**, and **Figs. S6**.

H2A peptide progress curves were compared. When treating the ^15^N-labeled H2A(1-28) F25Y with PADs, we found that all arginine residues showed evidence of citrullination (**Figs. 4A, 8**). PADs 1 and 2 fully citrullinated residues R3, R11, R17, and R20 within the duration of the experiment, with reactions reaching completion within ∼5 or ∼10 hours, respectively. Although PAD4 fully citrullinated R3, R17, and R20 within ∼5 hours, R11 citrullination was not complete within the experimental timeframe. PAD3 showed evidence of citrullination activity at all four positions, but over a substantially longer timescale, and no positions reached completion by the end of data collection (**Fig. 8**). The non-tail residue R20 was the preferred target for all four PADs (**Fig. 4A**). While PADs 1, 2, and 3 each citrullinated the two tail arginines (R3 and R11) similarly, PAD4 had a strong preference for R3 (**Fig. 4A**).

### Evaluation of nucleosomal H2A tail substrate: citrullination leads to visibility of additional tail residues in the NCP

In the context of the nucleosome, histone H2A uniquely contains both N-terminal (residues 1-11) and C-terminal (residues 120-129) tails (**Fig. 1B**). However, only the N-terminal tail contains arginines. An initial spectrum of the unmodified ^15^N-H2A-NCP reveals that, prior to modification, residues R3 through K9 are visible for the N-terminal tail, along with T120 through K129 for the C-terminal tail (**Fig. S3**), consistent with the observations of Ohtomo, et al. (66). Thus, R3 is the sole visible arginine for the unmodified ^15^N-H2A-NCP (**Figs. 2, 9**).

Similar to observations with the nucleosomal H4 tail, additional residue(s) are observed in the spectra upon citrullination (**Fig. S3**). A10 becomes observable in the spectra with all PADs, and Cit11 additionally becomes observable with PAD1 citrullination (**Fig. 9, bottom**), suggesting increased tail dynamics on the ps-ns timescale. The near-global increase in intensity across the N-terminal tail, which is also observed for the N-terminal portion of the C-terminal tail, supports an increase in dynamics (**Fig. S19**). However, contributions from the numerous nucleosomal citrullination events (at different positions and histones) cannot be disentangled with the current data.

**Figure 9.**
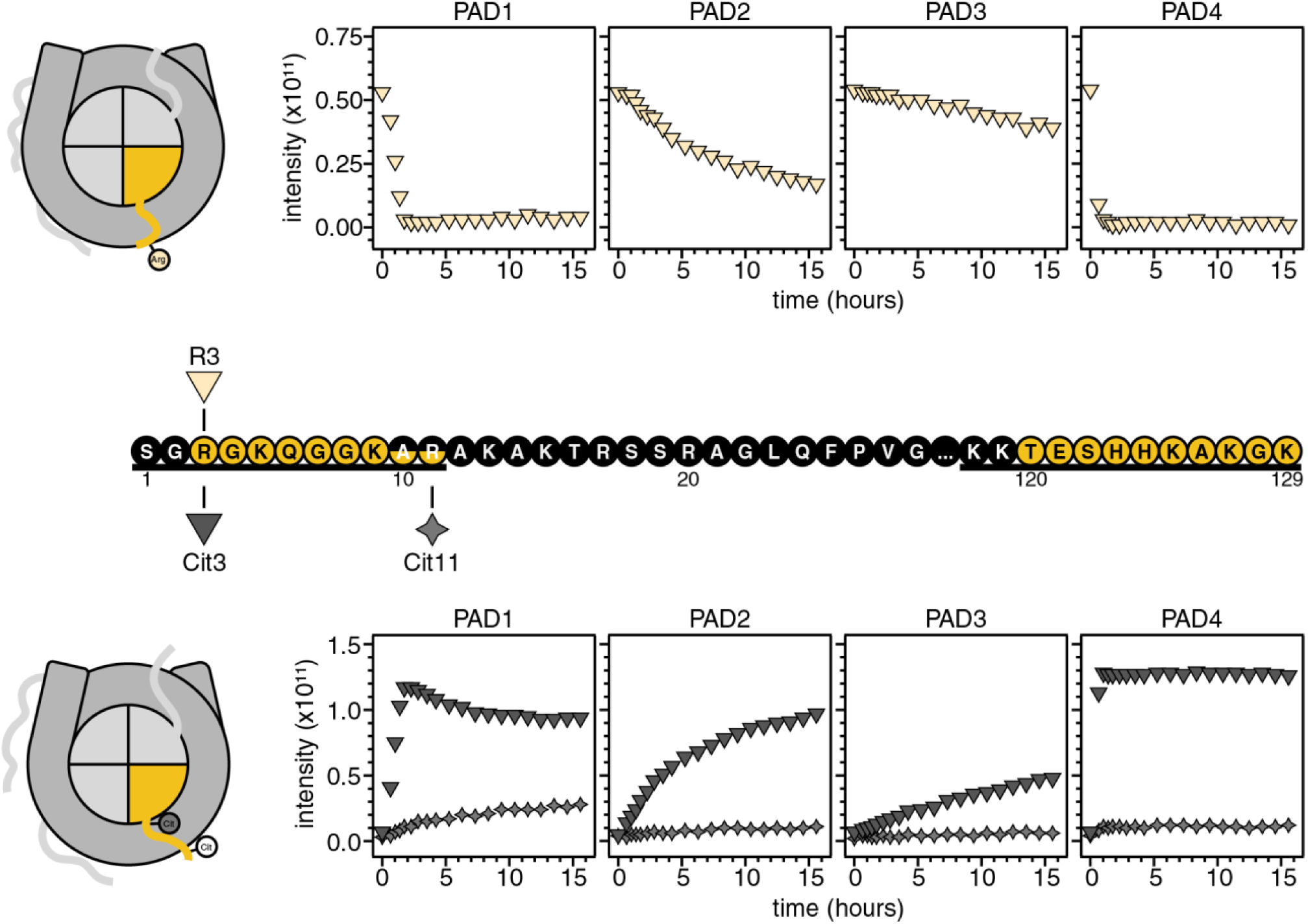
The sole visible arginine in H2A-NCP is citrullinated by PADs 1-4. Top: Cartoon of unmodified H2A-NCP (left) and progress curve graphs of the sole observable arginine in ^15^N-H2A-NCP for PADs 1, 2, 3, 4 (right). Arginine progress curves are denoted by the shapes found above the H2A sequence (middle). Black and yellow residues represent unobserved and observed residues, respectively. Citrullination of the NCP resolves the H2A tail through A10 (for PADs 2-4) or Cit11 (for PAD1) (see **Fig. S3**). Bottom: Cartoon of H2A-NCP (left) and citrulline progress curves of ^15^N-H2A-NCP for PADs 1, 2, 3, 4 (right). Citrulline curves are denoted by the shapes found immediately below the H2A sequence. Please note differences in arginine vs. citrulline y-axis scaling. Data were collected with 50 µM ^15^N-H2A-NCP (equivalent to 100 µM ^15^N-labeled histone) on an 800 MHz Bruker NMR spectrometer at 25°C in 20 mM MOPS pH 7.0, 100 mM KCl, 2 mM CaCl_2_, 0.5 mM TCEP, 0.1 mM EDTA, and 5% D_2_O. PADs were added to a final concentration of 200 nM. Data were collected as n=1.

Once treated with the PADs, we observe citrullination of R3 to varying degrees by all the PADs, consistent with the H2A tail peptide data described above (**Fig. 9, top**). Only PAD1 and PAD4 reach completion within the experimental timeframe. Citrullination of R11 is most substantial by PAD1, largely completing within the experimental timeframe. The nucleosomal context appears to exacerbate differences between R3 and R11 citrullination and also between the timescales of PAD1 and PAD2 reactions (**Fig. 4**)

### Evaluation of H2B tail peptide substrate: tail-adjacent residues are citrullinated in a peptide context

The human H2B sequence lacks arginine residues in the tail region. However, there are three arginine residues (R29, R31, and R33) in the region that passes between the two gyres of DNA within the NCP (**Fig. 10, middle**). Our H2B tail peptide sequence was extended out to residue V39 to include these three residues along with Y37 for concentration determination [referred to as H2B(1-39)].

**Figure 10.**
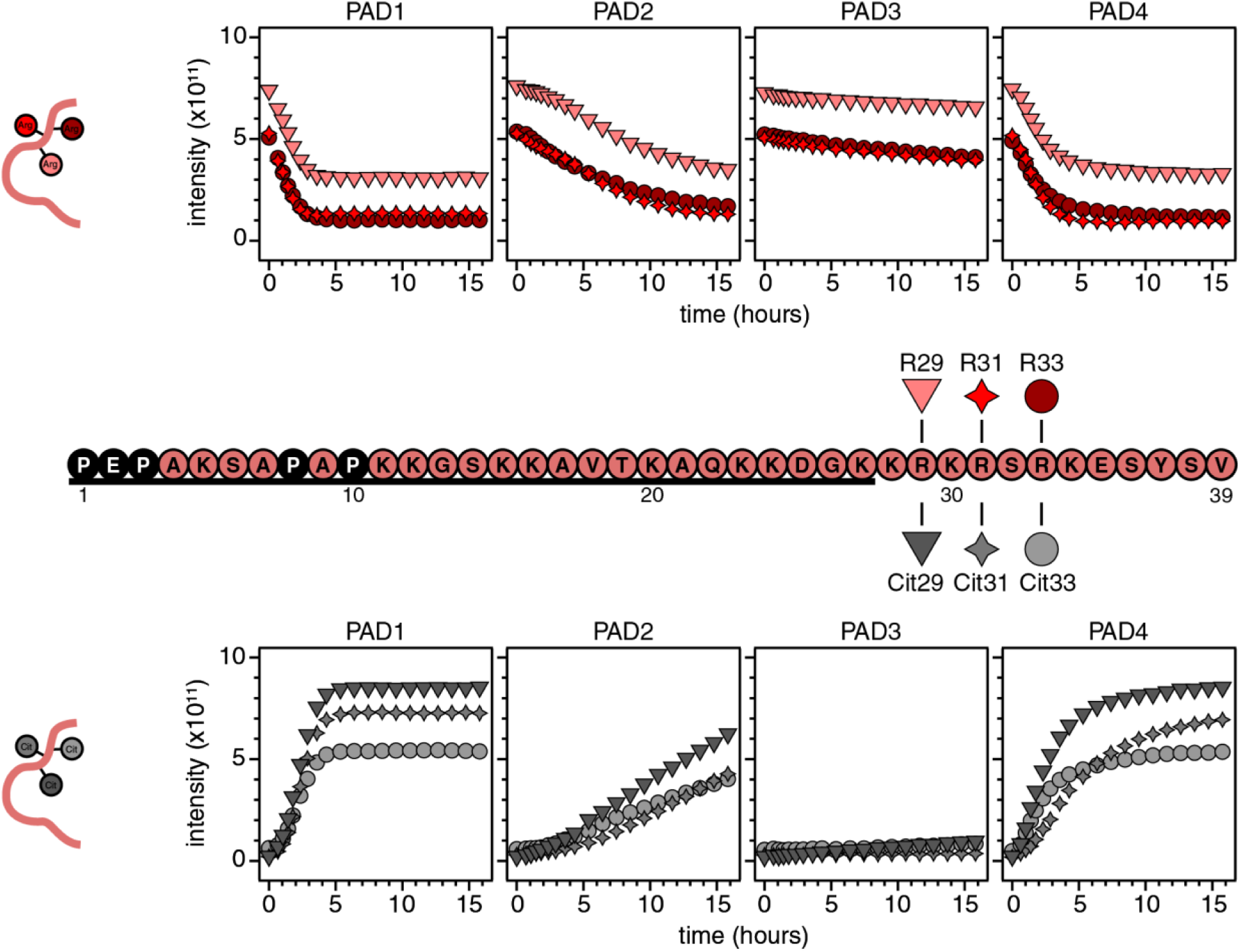
PADs 1, 2, and 4 citrullinate H2B peptide arginines, but minimal citrullination is observed by PAD3. Top: Cartoon of unmodified H2B peptide (left) and ^15^N-H2B peptide arginine progress curves for PADs 1, 2, 3, 4 (right). Progress curves are denoted by the shapes found above the H2B(1-39) peptide sequence (middle). Black and red residues represent unobserved and observed residues, respectively. Bottom: Cartoon of citrullinated H2B peptide (left) and citrulline progress curve graphs of ^15^N-H2B peptide for PADs 1, 2, 3, 4 (right). Citrulline progress curves are denoted by the shapes found immediately below the H2B peptide sequence. Data were collected with 100 µM ^15^N-H2B(1-39) on an 800 MHz Bruker NMR spectrometer at 10°C in 20 mM MOPS pH 7.0, 100 mM KCl, 2 mM CaCl_2_, 0.5 mM TCEP, 0.1 mM EDTA, and 5% D_2_O. PADs were added to a final concentration of 200 nM. Data were collected as n=1. For additional histone H2B tail peptide assays conducted at different conditions, see **Supplemental Methods**, **Supplemental Results**, and **Fig. S7**.

When treating ^15^N-labeled H2B(1-39) with each PAD, the three arginines were fully citrullinated by PAD1 and PAD4 within ∼4 hours and ∼5 hours, respectively. These arginines were only partially citrullinated by PAD2 and minimally modified by PAD3 within the experimental timeframe (**Fig. 10, 4A**). Each of the four PAD isozymes citrullinates the three arginines on a similar timescale.

### Evaluation of nucleosomal H2B tail substrate: nucleosomal tail intensities increase

We then asked what effects, if any, would be observed upon citrullination of ^15^N-H2B-NCP. Due to the lack of H2B tail arginines, we opted to track known lysine acetylation sites (K5, K11, K16, K20, and K27) spaced across the tail to test whether the H2B tail senses citrullination at other sites (**Fig. 11, top**). Slight increases in the raw intensities of these lysines are observed following PAD addition to ^15^N-H2B-NCP (**Fig. 11, middle**).

**Figure 11.**
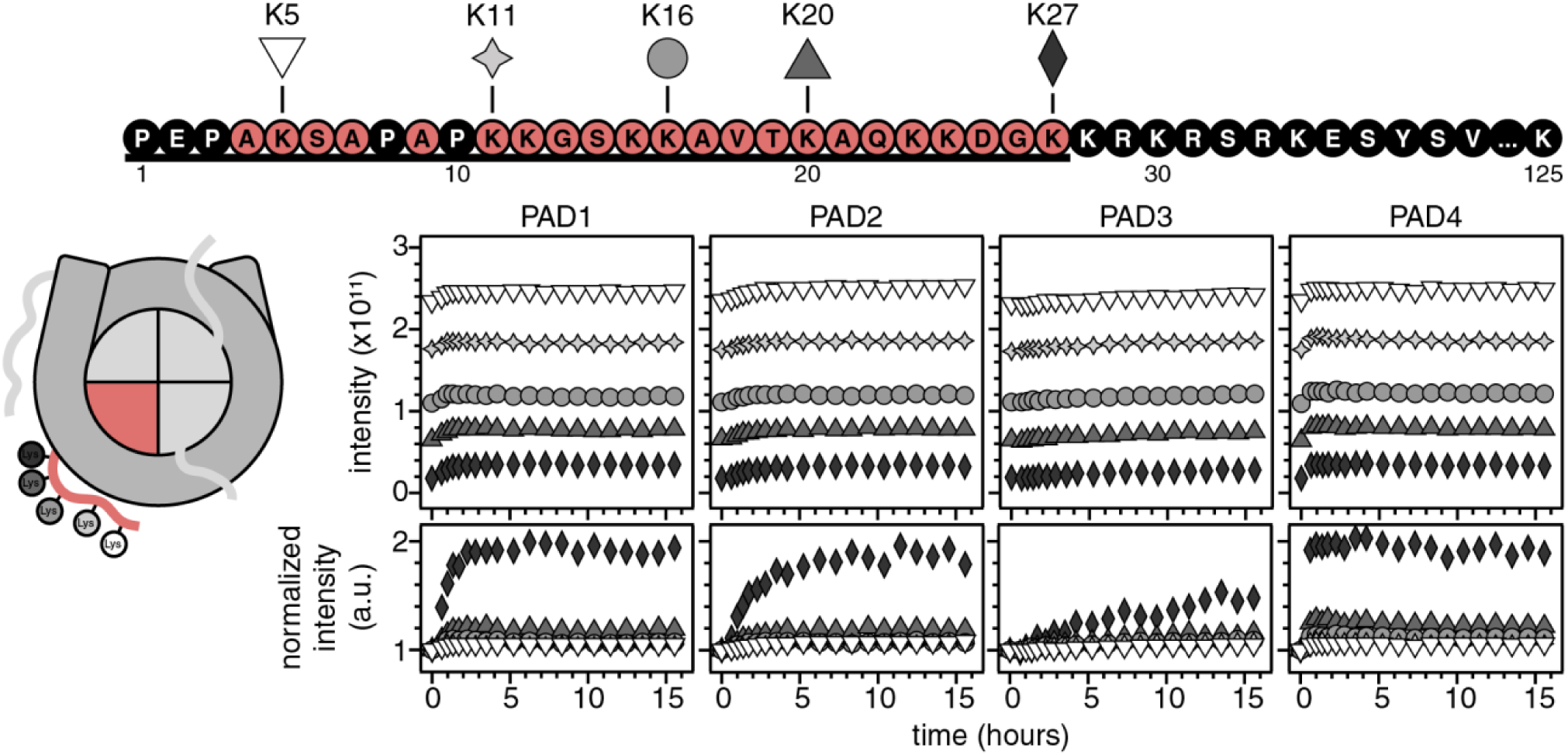
Citrullination of adjacent histones affects H2B lysine acetylation sites in the nucleosome. Top: H2B sequence. Black and red residues represent unobserved and observed residues, respectively. Left: cartoon of the H2B-NCP. Right, middle: Graphs of raw peak intensities of K5, K11, K16, K20, K27 as a function of time following PAD addition. Right, bottom: Graphs of scaled intensities (with respect to t=0) of K5, K11, K16, K20, K27 for PADs 1, 2, 3, and 4. The greatest relative intensity change is sensed closest to the core, at K27, suggesting the H2B tail becomes more dynamic in the NCP when adjacent histones are citrullinated. Data were collected with 50 µM ^15^N-H2B-NCP (equivalent to 100 µM ^15^N-labeled histone) on an 800 MHz Bruker NMR spectrometer at 25°C in 20 mM MOPS pH 7.0, 100 mM KCl, 2 mM CaCl_2_, 0.5 mM TCEP, 0.1 mM EDTA, and 5% D_2_O. PADs were added to a final concentration of 200 nM. Data were collected as n=1.

Scaling lysine intensities to time point zero reveals that relative intensities increase the greatest toward the core, at residue K27, and less so as the residues near the N-terminus (**Fig. 11, bottom**). The time-dependence of the intensity increases varies with PAD isozyme—intensities largely reach maxima by ∼1, 2, and 4 hours for PAD4, PAD1, and PAD2, respectively, while intensities do not plateau within the experimental timeframe for PAD3. These effects are not confined to the lysines (**Supplementary Fig. S20**).

There are a few plausible, nonexclusive explanations for these observations: i) R29, R31, and/or R33 are citrullinated and increase the dynamics of the H2B tail, and ii) citrullination of other histones within the complex increase H2B tail dynamics. R29, R31, and R33 do not appear in the NCP spectra, which could be because they are not citrullinated or because their effective tumbling is not fast enough to observe via backbone amides, even when citrullinated. It is interesting to look across the progress curves for the nucleosomal H3, H4, and H2A tails (**Figs. 5, 7, 9**) to find curves with plateaus matching H2B (**Fig. 11**) for each PAD: the H4 arginines (especially R3) and H2A R3 are similar for PADs 1 and 4, while H4 R3 and perhaps H3 R8 are similar for PAD2. This is speculative, as (with the current data) we cannot disentangle the contributions from individual citrullination events from the joint action of the citrullinations, which together could reach a critical citrullination threshold. It is, however, interesting to note that PAD4-treated H2B tail intensities increase the fastest (across PADs), but PAD1 most uniformly citrullinates (across histone tails). This suggests the possibility that the H4 arginines (especially R3) and/or H2A R3 have the largest impact on H2B tail mobility.

## Discussion

To better understand histone citrullination both in the context of isolated N-terminal sequences, as well as in the greater context of the nucleosome, we utilized an *in vitro* ^1^H-^15^N NMR-based enzymatic assay to observe how the PADs modify histone arginines when multiple substrates are simultaneously available for modification. In this assay, we can observe both arginine decay and citrulline growth simultaneously. We found that available arginine residues within the H2A, H2B, H3, and H4 histone tail peptides are modified by PADs 1, 2, and 4, to varying degrees. This is distinct from PAD3, which modifies H2A, partially modifies H4, and leaves H3 tail arginines unmodified (yet modifies the typically inaccessible R40/42 residues). When NCPs are treated with PADs, we find that, as expected, the same arginines that are modified in the N-terminal tails of the histone peptides are also modified within the context of the NCP (**Fig. 12A**). Interestingly, treatment of the ^15^N-NCPs with the PADs leads to increasing peak intensities and, in some instances, altered progress curves, suggesting that additional processes are involved as citrullination occurs, such as increasing tail dynamics.

**Figure 12.**
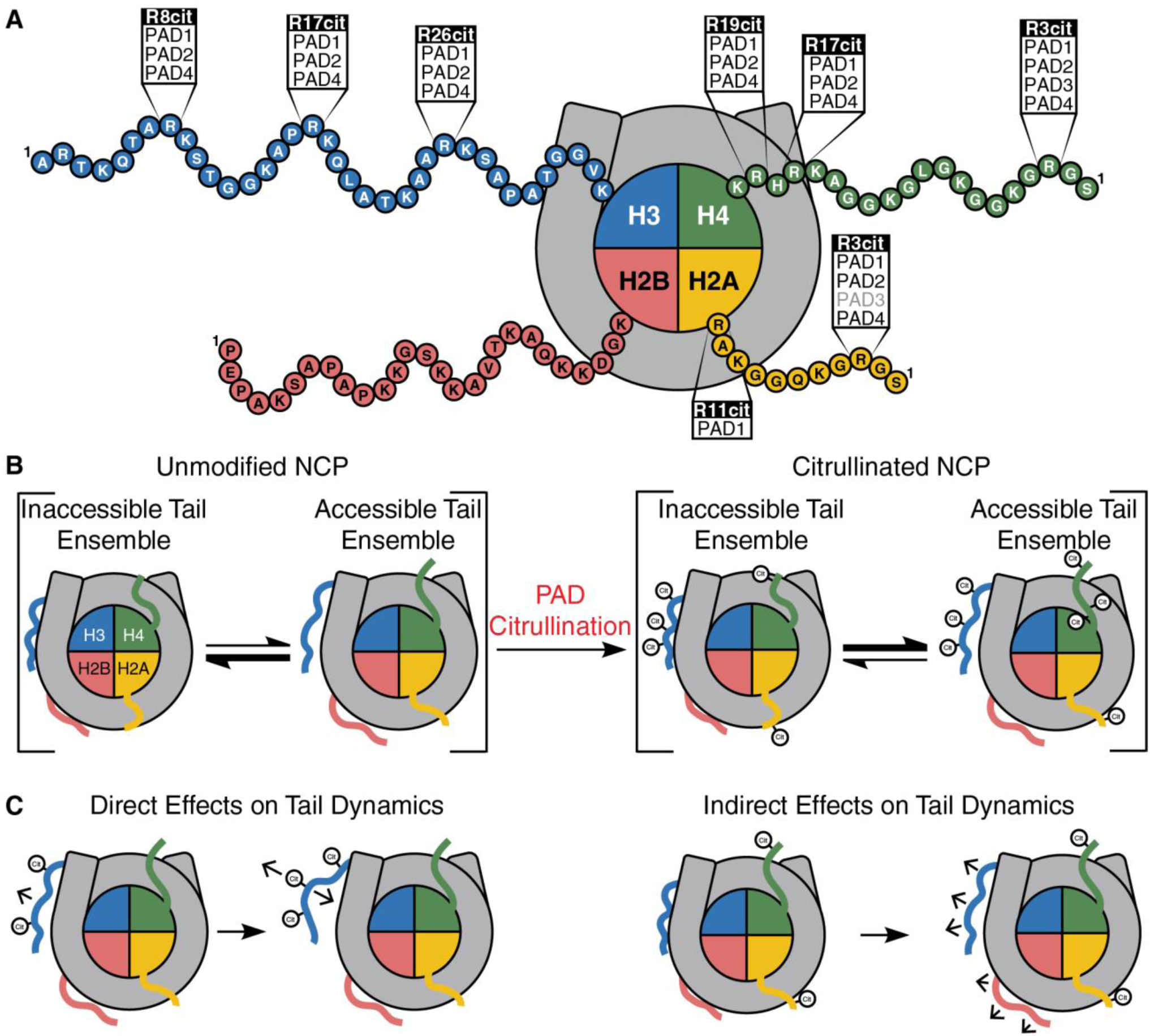
Summary of PAD histone citrullination sites in nucleosomes and proposed models for effects of nucleosome citrullination. **A.** PADs 1, 2, and 4 largely citrullinate all observable and accessible arginines within the N-terminal intrinsically disordered H3, H4, and H2A histone tails. In contrast, PAD3 only citrullinates H4 R3 and H2A R3. **B.** The histone tails dynamically interact with nucleosomal DNA, predominantly existing in inaccessible tail ensembles (left). PAD citrullination shifts the histone tail equilibria toward accessible ensembles (right). (Note that the cartoons are simplified depictions and are not intended to capture all conformations.) **C.** Citrullination of nucleosomal histone tails by the PADs may affect the dynamics of a given tail i. directly, by reducing the charge of the tail through successive citrullination events (left), or ii. indirectly, by citrullinating adjacent histones (right).

To gain additional insight into the differential histone modification sites and times of the PADs observed in our study, we considered our results in the context of structural studies of the isozymes. Crystal structures obtained with PAD4 and short, histone tail peptides H4(1-10), H3(4-13), and H3(14-23) highlight the importance of the residues flanking the arginine in the substrate (86). Consistent interactions between PAD4 and the various histone peptides include the sidechains of Q346 and W347 with the substrate ‘i-2’ (were ‘i’ is the arginine) backbone carbonyl, the R374 sidechain with the ‘i-1’ and ‘i’ backbone carbonyls, the R639 backbone carbonyl with the ‘i’ backbone amide, and two water-mediated interactions (involving the R372 sidechain and V469 backbone carbonyl along with ‘i+1’ and ‘i+2’ backbone carbonyls, respectively). Additional interactions may be found with the residues preceding the substrate arginine, depending on the substrate. Specifically, PAD4 D344 and Q346 engage the side chain hydroxyl group of the ‘i-2’ residue (S1) in H4(1-10), while Q346 additionally engages the backbone carbonyl of S1 (**Figs. S21**). With H3 tail peptides, only Q346 interacts with the ‘i-2’ residue; Q346 interacts with the side chain and backbone of T6 in H3(4-13) but only engages the backbone of A15 in H3(14-23). Our PAD4 results are consistent with these structural interactions: we observe fast citrullination of H4 R3 and H2A R3, which share the same N-terminal sequence, and moderate or slow modification of H3 R8 and R17, respectively. Consistent with these PAD4-histone interactions, a recent publication by Laposchan and coworkers demonstrated how mutations to PAD4 residues Q346 and R639, both mentioned above, along with G403 and H640 can tune PAD4 to have a broader substrate profile akin to PAD2 (87). Furthermore, they were able to subcategorize PAD1 and PAD2 as less specific isozymes, whereas PAD3 and PAD4 exhibit more specificity via their *in vitro* cell lysate PAD citrullination pipeline. This is supported by a combinatorial analysis conducted by Borbély et. al. of ‘i-1’ and ‘i+1’ residues nestled in a 9-mer peptide reacted with both PADs 2 and 4 PAD2 that again show PAD2 to have a much broader citrullination efficiency than PAD4 (88). When we align the sequences for PADs 1-3 to PAD4, we find that neither D344 or Q346 are conserved in their sequences (**Fig. S22**). In our sequences, D344 is replaced by asparagine in PADs 1 and 3, and arginine in PAD2, although this may have less impact on sequence specificity because D344 interacts with the substrate via its backbone carbonyl. Q346 is replaced by asparagine, aspartic acid, and arginine in PADs 1, 2, and 3, respectively, and may have a greater impact on specificity due to Q346 side-chain interactions with the substrate. Interestingly, active site volumes were found to be larger in PAD3 compared to PADs 1, 2, and 4, which may further confer differences in specificity and modification rates (86, 89–93). Continued studies on the sequence and structure of the PADs may further shed light on the specificity of the isozymes.

Many of our observations are consistent with prior studies (20, 21, 33, 36, 37, 46, 94–96): histone H3 is modified by PADs 1, 2, and 4, H4 is modified to varying degrees by the PADs, and H2A is modified by PAD4. In the H3 histone tail peptide, we observe citrullination of R8, R17, and R26 by PADs 1, 2, and 4, with R17 largely modified the slowest of the three directly observable arginines (**Fig 3**). Additionally, indirect readout through T3 indicates that PADs 1 and 2 modify R2 slowly, while virtually no modification is observed with PAD4 at these conditions (**Fig. 3**). Taken together, we speculate that the PADs uniformly disfavor substrates with a proline at the ‘i-1’ position (as with H3 R17) and that lack an ‘i-2 ‘residue (as with H3 R2). Interestingly, all four PADs citrullinate H3 R40 and R42 at comparable rates between the isozymes (**Fig. 4**), demonstrating shared specificity. Previously, systematic kinetic characterization of the PADs with histone H4 tail peptides uncovered that R3 and R19 are targets of PAD3 and PAD4, while R17 is a target of PAD1 (36). In our experiments, PAD1 citrullinates all four H4 tail arginines with similar timings overall, whereas PAD3 and PAD4 modify R23 at faster rates than either R17 or R19 (**Fig. 6**). In our extended histone H4 tail peptide, R17 and R19 appear to be weaker sites of modification by PAD3 and PAD4. Additionally, for PAD4, R3 is a major target of citrullination, while PAD3 only displays modest modification of the residue (**Fig. 6**). In the H4 histone tail peptide, R3 and R23 appear to be broad targets of the PADs. We have also shown that H2A tail peptide can be modified by all PADs, and PADs 1, 2, and 4 citrullinate H2B peptide arginines C-terminal to the tail region. Differences between the most abundant citrullination sites in cells (both general and PAD4-specific) (96) and the relative timings that we observe across the nucleosomal histone tails suggest that additional nuclear factors regulate histone citrullination in vivo.

Recently, NMR has been utilized to better understand dynamic interactions and modifiability of histone tails in the context of the nucleosome (6, 7, 28, 59, 66, 70, 72, 79, 97). Our lab utilized NMR spin relaxation experiments, which provide insights into molecular motions on the ps-ns time scale, with H3 tail arginine-to-glutamine mutations to better understand the role of histone tail arginines within the NCP (28). We proposed that arginines within the histone H3 tail act as anchor points via dynamic interactions with nucleosomal DNA. Individual charge-neutralizing mutations (R2Q, R8Q, R17Q, or R26Q) demonstrate regional increases in H3 tail mobility while simultaneous mutation of all four H3 tail arginines (R2/8/17/26Q) leads to global increases in H3 tail mobility. The near-global increases in non-arginine intensity and the transient increases in H3 tail arginine intensity following PAD addition support that enzymatic conversion of arginine to citrulline similarly increases tail mobility. We propose that citrullination shifts the nucleosome ensemble from favoring a largely inaccessible histone tail ensemble toward an accessible ensemble (**Fig. 12B**). Furthermore, NMR has been used to show that deposition of charge-modulating PTMs on one histone tail within the nucleosomal context can increase the dynamics and rate of subsequent modification on the same histone tail or even a different histone tail (6)(59, 70). We suspect that citrullination of H3, H4, and/or H2A histone tails within the nucleosome similarly leads to increased intensities across non-citrullinated tails (**Figs. 5, 11**, **S20**). Together, these studies support that modifications to histone tails affect the accessibility of the same or other histone tails within the complex in a form of crosstalk and suggest that initial citrullination of a histone tail may increase the rate at which successive modification events (whether citrullination or other) occur (**Fig. 12C**). Thus, increased histone tail dynamics brought on by citrullination may make the nucleosome more susceptible to additional epigenetic PTMs that lead to recruitment of transcription factors and alter the regulation of the underlying DNA (6, 7). Reducing the ability of histone tails to form strong interactions with the DNA of nearby nucleosomes through citrullination has implications for chromatin decondensation and regulation (20, 21, 45, 46, 98, 99). Future studies will be needed to deconvolute the relative contributions of different tail positions and modifications to histone tail crosstalk and chromatin organization.

Since our NMR kinetic assays monitor only the intrinsically disordered histone tails, it is conceivable that the increased intensities observed during NCP citrullination reactions reflect unobservable factors. These include citrullination of histone core residues and changes in DNA conformation and dynamics as the histones are modified. Changes in DNA conformation, for example, more open and flexible DNA at the entry/exit sites, could conceivably also lead to an overall increase in the intensity of histone tails interacting with these sites. It is interesting to consider that, if important histone-DNA contacts are lost due to citrullination, the histone octamer may become more prone to eviction by nucleosome remodelers or on its own, thus increasing nuclear DNA accessibility and gene expression. Additional NMR approaches may resolve core effects as histones are citrullinated by the PADs (100). Assays that report on distance, such as FRET, may reveal differences in the DNA conformation between citrullinated and unmodified NCPs (101, 102). Together, these additional molecular-level experiments will shed light on how citrullination affects chromatin at the nucleosome level.

As the effects of histone citrullination on nucleosome dynamics and chromatin organization remain to be fully elucidated, so do questions on the PADs themselves. *In vitro* studies demonstrate the need for relatively high calcium concentrations for PAD activation, on the order of millimolar (32–34, 36, 37), which are much higher than typical cellular calcium concentrations (103). Recent studies suggest that pore-forming complexes in NETosis-like processes may provide sufficient calcium concentrations for PAD activation (14–17). However, instances of specific PAD modification, such as PAD2 citrullination of H3 R26 in response to estradiol stimulation (20, 21), may require a different mode of calcium activation. Darrah et al. demonstrated that cross-reactive anti-PAD3 antibodies can increase PAD4 calcium sensitivity, lowering the amount required for PAD4 activation (104). Conceivably, other associations with the PADs may reduce the calcium needed for activation and perhaps impart an additional layer of specificity onto the enzymes. Recently, PADs 2-4 were shown to play an interrelated regulatory role with the cBAF complex-specific subunits BAF250A and DPF1 in skeletal muscle differentiation, uncovering a broader role for the PADs in chromatin regulation beyond histone citrullination (105). Finally, questions remain as to whether citrulline “readers” or “erasers” exist.

## Data availability

The data underlying this article are available in the manuscript, the supplementary materials, and upon reasonable request. Assignments are deposited in the Biological Magnetic Resonance Data Bank (BMRB) for PAD1 citrullinated histone H3(1-44) (53978), histone H4(1-33) T30Y (53979), PAD1 citrullinated histone H4(1-33) T30Y (53980), histone H2A(1-28) F25Y (53981), PAD1 citrullinated histone H2A(1-28) F25Y (53982), histone H2B(1-39) (53983), PAD1 citrullinated histone H2B(1-39) (53984).

## Supporting information

This article contains Supporting Information.

## Acknowledgments

Thanks to Drs. Catherine Musselman, Michael Poirier, and Karolin Luger for providing histone plasmids. Thank you to Sarah Meidl Zahorodny for purifying full-length histones and Widom 601 DNA. Thank you to Dr. Paul R. Thompson (UMass) for providing plasmids and purification protocols for PAD isozymes and for helpful discussion. Thank you to Drs. Carlos Castañeda and Scott Showalter for helpful discussions. Thanks to Dr. Dawn Wenzel for providing PreScission Protease. Thank you to Dr. Francis Peterson for providing the His-Sumo and Ulp1 plasmids, for use of the HPLC, and for assistance in setting up triple resonance experiments. Thank you to the MCW Department of Biochemistry for the use of the mass spectrometer.

## Author contributions

E.M. conceived of the initial project, acquired funding, and supervised the research. A.K. and E.M. designed the studies. A.K. prepared samples, performed the research, analyzed and visualized the data, and wrote the manuscript. E.M. edited the manuscript and contributed to the writing.

## Funding and additional information

This work was supported by the National Institutes of Health: R35 GM142594 to E.A.M. and S10 OD0250000 to the MCW NMR Facility. This study made use of NMRbox: National Center for Biomolecular NMR Data Processing and Analysis, a Biomedical Technology Research Resource (BTRR), which is supported by NIH grant P41GM111135 (NIGMS). This manuscript is subject to the NIH Public Access Policy. Through acceptance of the federal funding used to support the research reported in this manuscript, the NIH has been given a right to make this manuscript publicly available in PubMed Central upon the Official Date of Publication, as defined by the NIH. The content is solely the responsibility of the authors and does not necessarily represent the official views of the National Institutes of Health.

## Conflict of interest

The authors declare that they have no conflicts of interest with the contents of this article.

## Materials & correspondence

Correspondence and material requests should be addressed to Emma Morrison at.

## Abbreviations

PTM: post-translational modification
PAD: protein-arginine deiminase
NMR: nuclear magnetic resonance
HMQC: heteronuclear multiple quantum coherence
NCP: nucleosome core particle
IDR: intrinsically disordered region.

## Notes

### Competing Interest Statement

The authors have declared no competing interest.

### Summary of Updates

Text has been revised for clarity. Additional experimental data and analyses are included.

