## Supporting Information for "Differential histone tail citrullination by protein arginine deiminases observed via NMR spectroscopy"

### Supplemental Methods

***PAD activity assay.*** To ensure that purified PAD enzymes were functional, the enzymatic activity for each PAD was assessed in a discontinuous colorimetric assay as previously described (1–5). Briefly, this assay measures the production of ureido-containing products over time by comparison to a citrulline standard curve and utilizes Nα-benzoyl-L-arginine ethyl ester (BAEE) as the test substrate. In a 96-well plate, BAEE samples at a range of concentrations were pre-incubated at 37°C for 10 minutes in activity assay buffer (100 mM Tris-HCl, pH 7.6, 50 mM NaCl, 2 mM DTT, and 10 mM $CaCl_2$). The highest BAEE concentrations used were 10 times the reported $K_m$ of each PAD for BAEE (2–4), and these concentrations were diluted in half with activity assay buffer seven times. Next, 10 µL of PAD at 1.2 µM was added to initiate the reaction (200 nM final concentration, 60 µL final reaction volume). To quench the reaction, 200 µL of freshly prepared color development reagent (COLDER) solution (1 part solution A [80 mM diacetyl monoxime, 2.0 mM thiosemicarbazide], and 3 parts solution B [3 M $H_3PO_4$, 6 M $H_2SO_4$, 2 mM $NH_4Fe(SO_4)_2$]) was added to each reaction well. For color development, the reaction mixtures were heated at 95°C for 30 minutes, after which the plate was allowed to cool for 10 minutes, and then the absorbance at 530 nm was measured for each reaction using a FlexStation 3 plate reader (Molecular Devices). Absorbance values were then compared to an L-citrulline standard curve to determine the amount of product formation. For each substrate concentration, the initial rate of reaction was determined from the slope of the first 3 time points of product formation as a function of time. The initial rate of reaction was then plotted as a function of substrate concentration and fit to the Michaelis-Menten equation ($V_0=(V_{max}*[S])/K_m+[S]$) to determine $V_{max}$ and $K_m$. The turnover number, $k_{cat}$, was calculated from the equation $V_{max}=k_{cat}*[E_{tot}]$, and the catalytic efficiency $k_{cat}/K_m$ was determined for each PAD with BAEE. Activity assays were performed in duplicate. Catalytic efficiencies were compared to the literature values to assess enzyme prep quality (2–5). Although model arginine compounds are commonly used to assess PAD activity, they are generally poor substrates for PAD3 (3, 4).

***PAD preparation for supplemental NMR assays***. PADs 1 (UniProt Q9ULC6, except containing L184Q and S266P), 2 (UniProt Q9Y2J8), and 3 (UniProt Q9ULW8) were recombinantly expressed with an N-terminal 10xHis-tag in BL21(DE3) E. coli (New England BioLabs). PAD4 (UniProt Q9UM07, except containing wild-type residues S55, A82, and A112) was recombinantly expressed with an N-terminal GST tag in Rosetta2 (DE3) pLysS E. coli (Novagen). Transformed bacteria in LB media were induced at an OD600 of ~0.6-0.7 with 50 µM (PADs 1-3) or 25 µM (PAD4) IPTG for overnight expression (~16 hours). Cells were lysed by sonication, and the lysate was clarified by centrifugation. His-tagged PADs were purified over a nickel affinity gravity-flow column and eluted with imidazole. GST-PAD4 was purified over a glutathione gravity-flow column and cleaved off the beads with PreScission Protease. PADs were further purified by Source 15Q anion exchange chromatography. Note that these PAD preps were not purified via size-exclusion chromatography. Fractions containing PAD were concentrated by benchtop centrifugation in 10k MWCO concentrators, and final PAD stocks were stored at -80°C.

***NMR-based citrullination assay for supplemental NMR assays.*** Data were collected with 50 µM $^{15}$N-labeled histone tail peptides on an 800 MHz Bruker AVANCE NEO NMR Spectrometer at 10°C in 20 mM MOPS pH 7.0, 150 mM KCl, 10 mM $CaCl_2$, 2 mM DTT, 1 mM EDTA, and 5% $D_2O$. PADs were added to a final concentration of 200 nM (1:250 enzyme-to-substrate molar ratio). Sequential $^1$H-$^{15}$N SOFAST-HMQC were collected in the same manner as detailed in **Methods**. Assays conducted with PADs 2 and 4 and H4, H2A, and H2B tail peptide substrates were performed using a room-temperature probe; all others were collected using a cryoprobe. Assays with PAD 2 and 4 used two different enzyme preps between H3 and H4, H2A, and H2B tail peptide substrates

**Supplemental Results**

**Peak assignments of histone H2A, H2B, and H4 tail peptides.**

To assess PAD citrullination sites and rates for histone tail peptides, triple resonance experiments [HNCACB, HN(CO)CACB, and HNCA] were conducted on $^{13}C/^{15}N$ isotopically labeled H2A(1-28) F25Y, H2B(1-39), and H4(1-33) T30Y and their PAD1-citrullinated forms to acquire backbone peak assignments. The $^{1}H$ chemical shift dispersion in the $^{1}H$-$^{15}N$ HSQC/HMQC is characteristic of intrinsically disordered regions, with citrullination having marginal effects on $^{1}H$ chemical shift dispersion within the peptides. The expected 31 and 35 resonances were observed in the H4 and H2B tail peptide spectra, respectively. However, residues H4 G2 and H2B E2 were only observed under the assignment assay conditions (i.e., higher concentrations) and were not observed during the kinetic NMR assays. The H2A tail peptide $^{1}H$-$^{15}N$ HSQC spectrum revealed more resonances than expected (25-26 expected versus 32 observed). Two resonances are observed for residues A21-G28 of H2A: minor peaks for each of the seven residues around P26 may result from a *cis*-isomer of P26.

**Comparison of histone peptide citrullination assays collected at different conditions.**

Two datasets are presented for histone peptide-PAD combinations to demonstrate reproducibility: 1) the data displayed in the main text figures (**Figs. 3, 4, 6, 8, 10**) were collected with 100 μM peptide, 1:500 enzyme-to-substrate molar ratio, and 2 mM $Ca^{2+}$ and 2) the data displayed in the supplemental figures (**Figs. S4-S7, S15**) were collected with 50 μM peptide, 1:250 enzyme-to-substrate molar ratio, and 10 mM $Ca^{2+}$ (see **Methods** and **Supplemental Methods** for additional details). When comparing the two data sets (Figs. S4-S7), the trends in the progress curves are largely the same. Overall, reactions with PAD4 (and to a lesser extent PAD2) are slower with condition set 2. Before running the final assays reported in the main text, we added a size-exclusion chromatography polishing step to the purifications to improve their purity. Note that the peak intensities for assays conducted with PADs 2 and 4 and H4, H2A, and H2B tail peptide substrates are substantially lower because the data were collected using a room-temperature (rather than cryo) probe. On the other hand, reactions with PAD1 are substantially faster with condition set 2. Between collecting the two data sets, it was discovered that the plasmid contained L184Q and S266P mutations (as compared to UniProt Q9ULC6), which are located outside of the catalytic domain. It is unclear whether the faster rates are attributable to these mutations or the higher calcium concentration. Altogether, considering these differences between the enzyme preps, the similarities between the two datasets support the robustness of the conclusions.

**Comparison of H3 tail $t_{50\%}$ values between H3 tail assays at different conditions.**

To demonstrate the suitability and reproducibility of the $t_{50\%}$ calculations, the additional H3 tail peptide NMR citrullination assays conducted under condition set 2 were analyzed in the same way and compared with the main text data (**Fig. S15**). Under both sets of conditions, PAD1 modifies R8, R26, and R42 at comparable rates, followed by R40, R17, and finally T3(R2). However, the $t_{50\%}$ values for the condition set 2 reactions were overall faster than for condition set 1 (see above). For PAD2-H3 tail citrullination, within both sets of conditions, R40 and R42 were modified the fastest, followed by R8. Similar modification rates of T3(R2) and R26 were observed, with R17 citrullination occurring at the slowest rate for PAD2. The $t_{50\%}$ values were overall slower for condition set 2 (see above). PAD3 assays under both sets of conditions demonstrate similar modification of R40 and R42, and essentially no modification of T3(R2), R8, R17, and R26. PAD4 modification of the H3 tail peptide under both sets of conditions demonstrates similar timing for R8, R40, and R42, with slower modification of T3(R2), R17, and R26. As with PAD2, the $t_{50\%}$ values were overall slower for condition set 2 (see above).

**Assessment of using i+1 residues as a proxy for arginine.**

In our assays, H3 R2 is unobserved in both the histone tail peptide and NCP spectra (**Fig. S3**). To determine if T3 is a suitable proxy for R2 citrullination in the H3 histone tail peptide, i+1 residues (H2A G4, H2A A21, and H4 G4) were analyzed in the same manner as their respective arginine residues (H2A R3, H2A R20, and H4 R3) (**Fig. S16A**), and the resulting $t_{50\%}$ values were compared (**Fig. S16B**). For PADs 1, 2, and 4, all the i+1 residues are consistent with their respective arginine (**Fig. S16B**). The exception is with H2A G4 and H4 G4, where the unmodified form of the peak is minimally sensitive to citrullination. With PAD3, the $t_{50\%}$ values determined from citrullinated forms of H2A G4 and H4 G4 peaks agree with H2A R3 and H4 R3, but the $t_{50\%}$ values determined from the unmodified peak forms are artificially small. Overall, these data demonstrate that the i+1 residue of an arginine and citrulline is a reasonable proxy for unobserved sites of citrullination.

**Comparison of reaction times.**

To better understand whether some histone arginines are preferred substrates of the PADs, progress curves of arginine decay and citrulline growth were fit (see **Methods** for details) to determine the time for 50% completion ($t_{50\%}$) (**Figure 4**). Due to differences in the shapes of progress curves across PAD-substrate combinations, multiple functions were used to fit the data; thus, we compared $t_{50\%}$ values rather than the fit rates. While arginine and citrulline $t_{50\%}$ values at a given position should match, factors such as peak overlap, effects from modification of adjacent residues, and changes in dynamics may influence the progress curves, leading to differences observed in $t_{50\%}$.

Of the H3 tail peptide assays, PAD1 citrullination appears the most similar across all the arginines, with R8, R26, R40, and R42 having similar rates of modification, followed by R17, and modification of R2 (observed via T3) being the slowest. PAD2 demonstrates slightly greater variation in $t_{50\%}$ values, with R40/R42 modified the fastest, followed by R8, R2 (via T3), and R26 at comparable rates, and R17 the slowest to be modified by PAD2. PAD3 exhibits the greatest specificity among the PADs, modifying only R40/R42, with virtually no modification of R2 (via T3), R8, R17, and R26 observed. PAD4 modifies R8, R40, and R42 at similar rates, with R17/R26 much slower and virtually no modification of R2 (via T3). Broadly, R40 and R42, despite being inaccessible in the nucleosome, are targets of all the PADs, while R8, R17, and R26 are only modified by PADs 1-3, and only PADs 1 and 2 appear to modify R2 (via T3).

For the H4 tail peptide, PAD1 broadly citrullinates R3, R17, R19, and R23 at similar rates. PAD2 modifies R3 the fastest, followed by R23, while R17 and R19 are the slowest to be citrullinated. R23 is the fastest of the four arginines citrullinated by PAD3, with slow modification of R3, and virtually no citrullination of R17 and R19. Like PAD2, PAD4 modifies R3 and R23 quickly, with much slower modification of R17 and R19. Together, the PADs largely modify H4 tail peptide R3 and R23, with either slower (PADs 1, 2, and 4) or no modification (PAD3) of R17 and R19.

In the H2A tail peptide, PAD1 citrullinates R20 the fastest, with R3, R11, and R17 having similar citrullination rates. Similarly, PAD2 modifies R3, R11, and R17 at comparable rates, with R20 citrullinated the fastest. As with the other two PADs, PAD3 modifies R20 the fastest of the four arginines present. Although slow modification of R3, R11, and R17 is observed, all $t_{50\%}$ values fall outside the ~16-hour measurement window. Interestingly, PAD4 quickly modifies R3 and R20, followed by R17, and comparatively slower modification of R11. Across the PADs, H2A R20 in the tail peptide appears to be a major target of citrullination with R3, R11, and R17 modified at similar rates for PADs 1 and 2, while PAD3 modifies these residues at very slow rates. Distinctly, PAD4 demonstrates greater specificity, with relatively fast modification of R3 but slower modification of R11.

In the H2B tail peptide, R29, R31, and R33 are clustered in a very basic region along with numerous lysine residues. The three arginines are largely citrullinated by PAD1 at similar rates. The same is true with PAD2. As with the H2A tail peptide, PAD3 modifies R29, R31, and R33 at very slow rates, with their $t_{50\%}$ values falling outside the data collection window. As with PAD1 and PAD2, PAD4 modifies R29, R31, and R33 at comparable rates. H2B R29, R31, and R33 are targets of PADs 1, 2, and 4, with minimal citrullination of the three residues by PAD3.

We additionally sought to compare reaction progress within the NCP context. However, non-monotonic progress curves for several nucleosomal H3 tail arginines lead us to believe that these arginine intensities are convoluted by dynamic interactions with the nucleosomal DNA. The progress curves for citrullines were largely monotonic and were thus fit to acquire $t_{50\%}$ values (**Figure 4B**). Within the nucleosomal H3 tail, Cit8 was the fastest to reach 50% across all PADs, followed by Cit26, and lastly by Cit17. In assays monitoring the nucleosomal H4 tail, Cit3 reached 50% before Cit17 for PADs 2, 3, and 4, while Cit17 was narrowly fastest with PAD1; Cit19 was the last residue to reach 50% completion. Similarly, in the nucleosomal H2A tail, Cit3 reached 50% completion faster than Cit11. While some differences are observed within the context of the peptides, the relative order at which residues reach their $t_{50\%}$ appears to be similar across the PAD isozymes within the context of the NCPs (although rates vary).

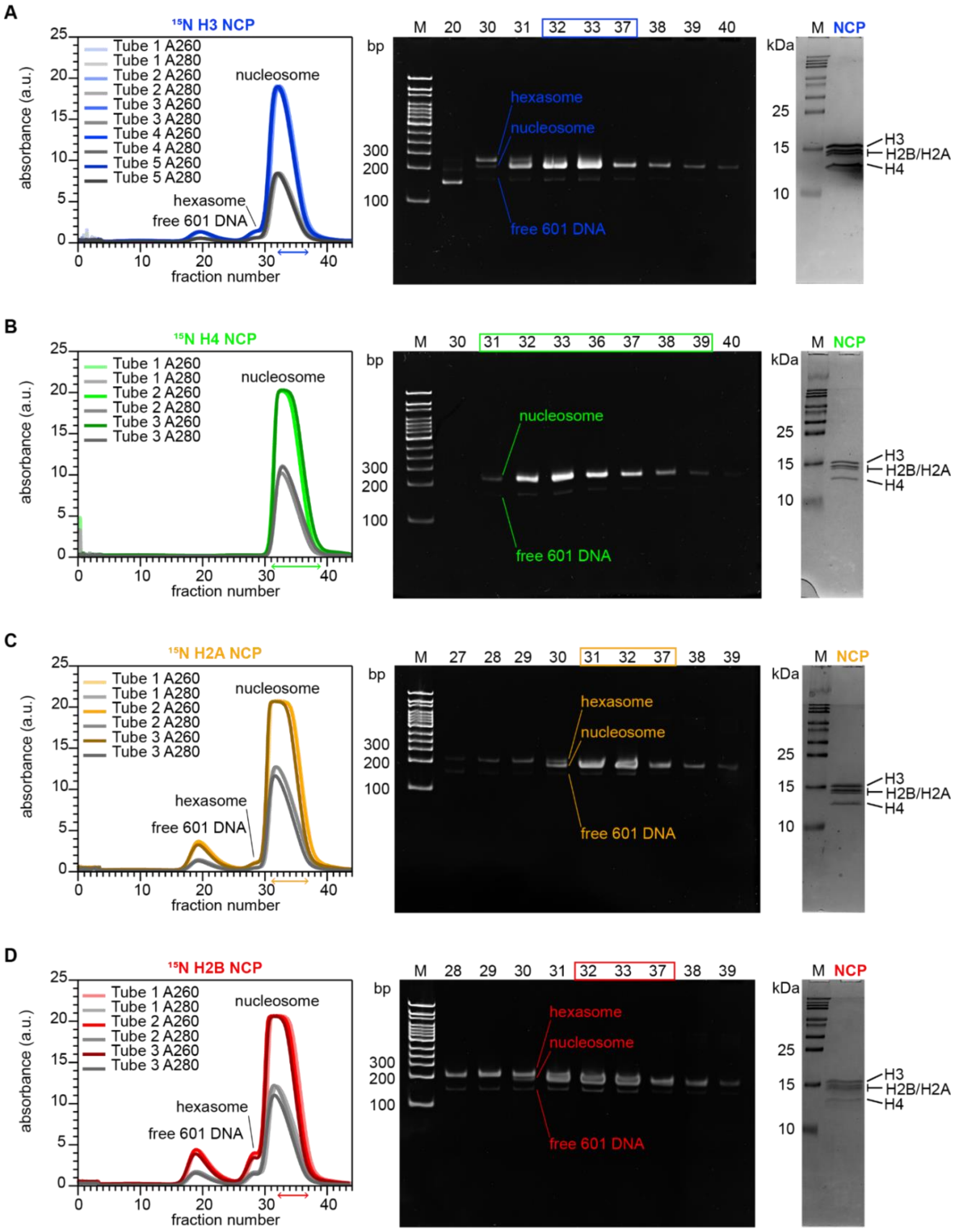


**Figure S1. Preparation quality of NCPs with isotopically labeled histones.** Left to right: Absorbance traces at 260 and 280 nm from sucrose gradient fractionation, 5% native-PAGE gels (stained with ethidium bromide) of sucrose gradient fractions, and 18% SDS-PAGE gels (stained with Coomassie) of final NCP samples with $^{15}$N-labeled H3 (**A**), H4 (**B**), H2A (**C**), and H2B (**D**). Double-sided arrows and boxed fraction numbers indicate the sucrose gradient fractions combined for the final $^{15}$N-labeled NCP sample.

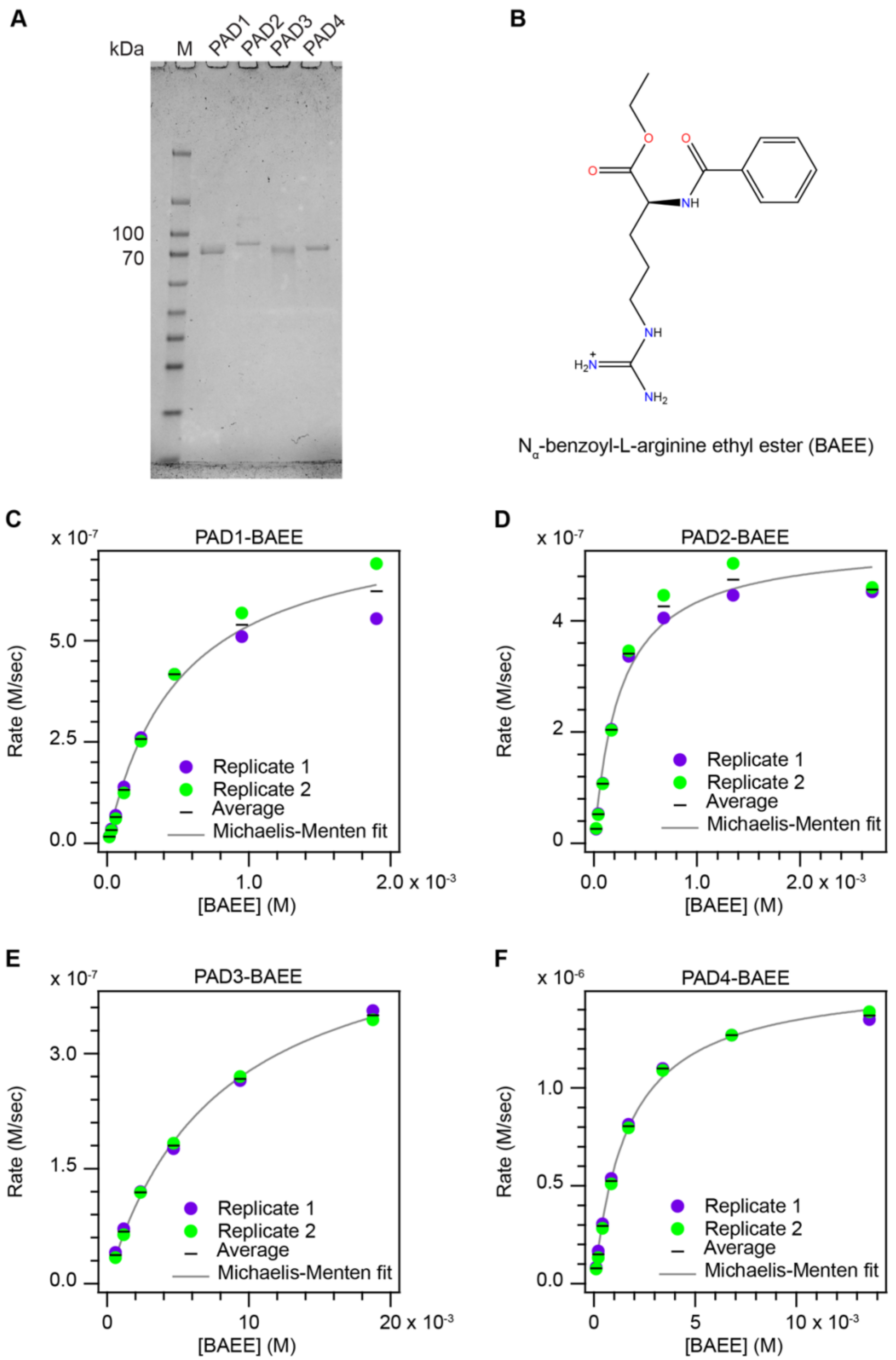


**G**

| Michaelis-Menten kinetic parameters for PAD BAEE reactions | | | | |
|---|---|---|---|---|
| | PAD1 | PAD2 | PAD3 | PAD4 |
| $k_{cat}$ ($s^{-1}$) | 4.1 ± 0.9 | 2.71 ± 0.15 | 2.46 ± 0.07 | 7.9 ± 0.2 |
| $K_m$ (M) | (5.2 ± 1.8) x $10^{-4}$ | (2.53 ± 0.10) x $10^{-4}$ | (77 ± 4) x $10^{-4}$ | (16.6 ± 1.6) x $10^{-4}$ |
| $k_{cat}/K_m$ ($M^{-1}s^{-1}$) | 8,100 ± 1200 | 10,700 ± 140 | 318 ± 8 | 4700 ± 300 |

**Figure S2. Activity assessment of purified, catalytically active PADs.** (**A**) SDS-PAGE gel (4-20% acrylamide gradient) of purified PADs 1-4 stained with Coomassie. (**B**) Chemical structure of model arginine substrate BAEE, commonly used in PAD activity assays. (**C**-**F**) Michaelis-Menten plots for PADs 1-4 and (**G**) table of BAEE Michaelis-Menten kinetic parameters determined for each PAD.

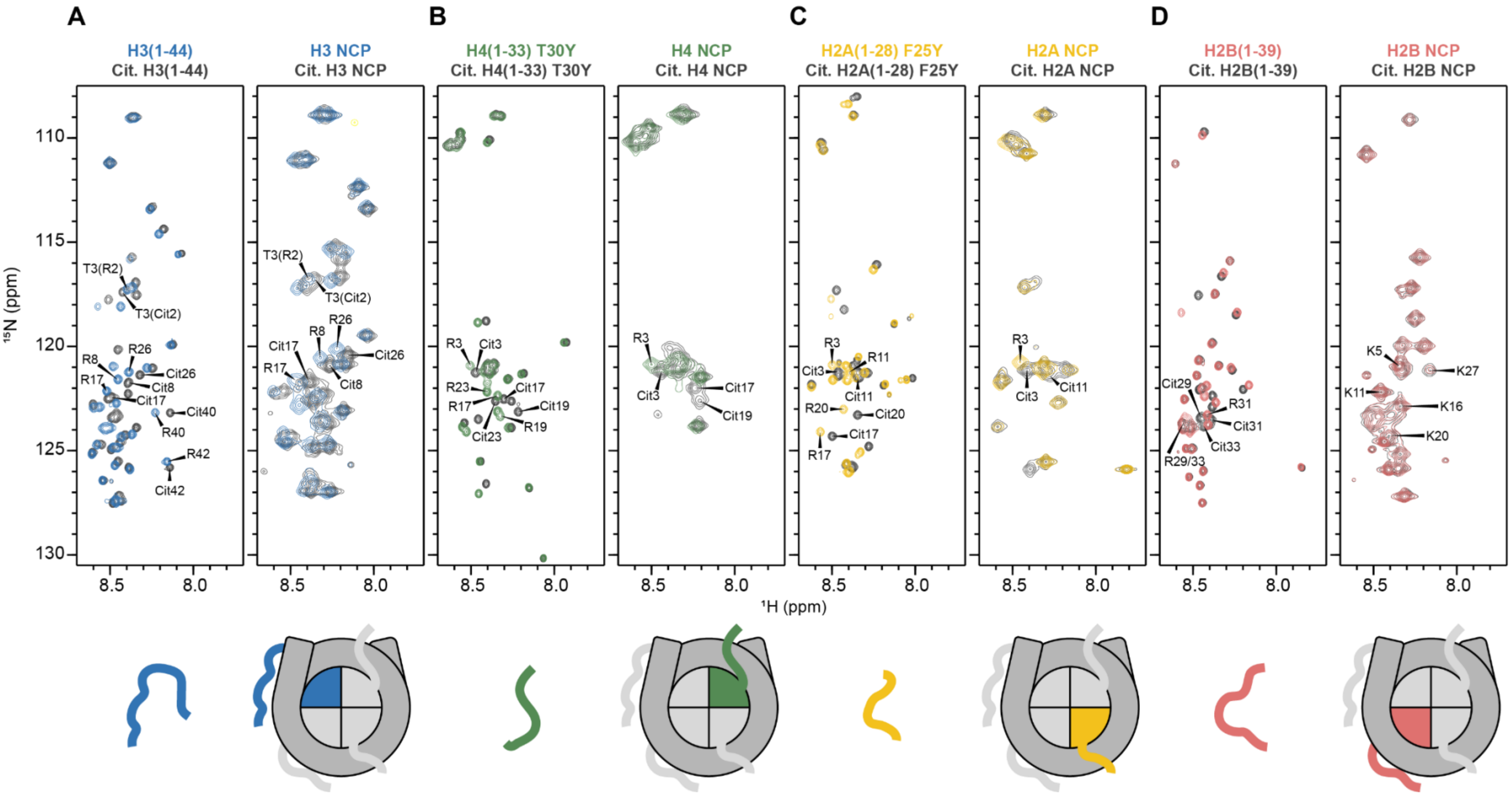


**Figure S3. Full $^{1}$H-$^{15}$N SOFAST-HMSQ spectra of $^{15}$N-labeled histone substrates with PAD1 citrullinated end states.** (**A**) Spectra of $^{15}$N-H3(1-44) (left, blue) and PAD1-citrullinated end state (left, grey). Spectra of $^{15}$N-H3 NCP (right, blue) and PAD1-citrullinated end state (right, grey). (**B**) Spectra of $^{15}$N-H4(1-33) T30Y (left, green) and PAD1-citrullinated end state (left, grey). Spectra of $^{15}$N-H4 NCP (right, green) and PAD1-citrullinated end state (right, grey). (**C**) Spectra of $^{15}$N-H2A(1-28) F25Y (left, yellow) and PAD1-citrullinated end state (left, grey). Spectra of $^{15}$N-H2A NCP (right, yellow) and PAD1-citrullinated end state (right, grey). (**D**) Spectra of $^{15}$N-H2B(1-39) (left, red) and PAD1-citrullinated end state (left, grey). Spectra of $^{15}$N-H2B NCP (right, red) and PAD1-citrullinated $^{15}$N-end state (right, grey). Arginines and citrullines are labeled.

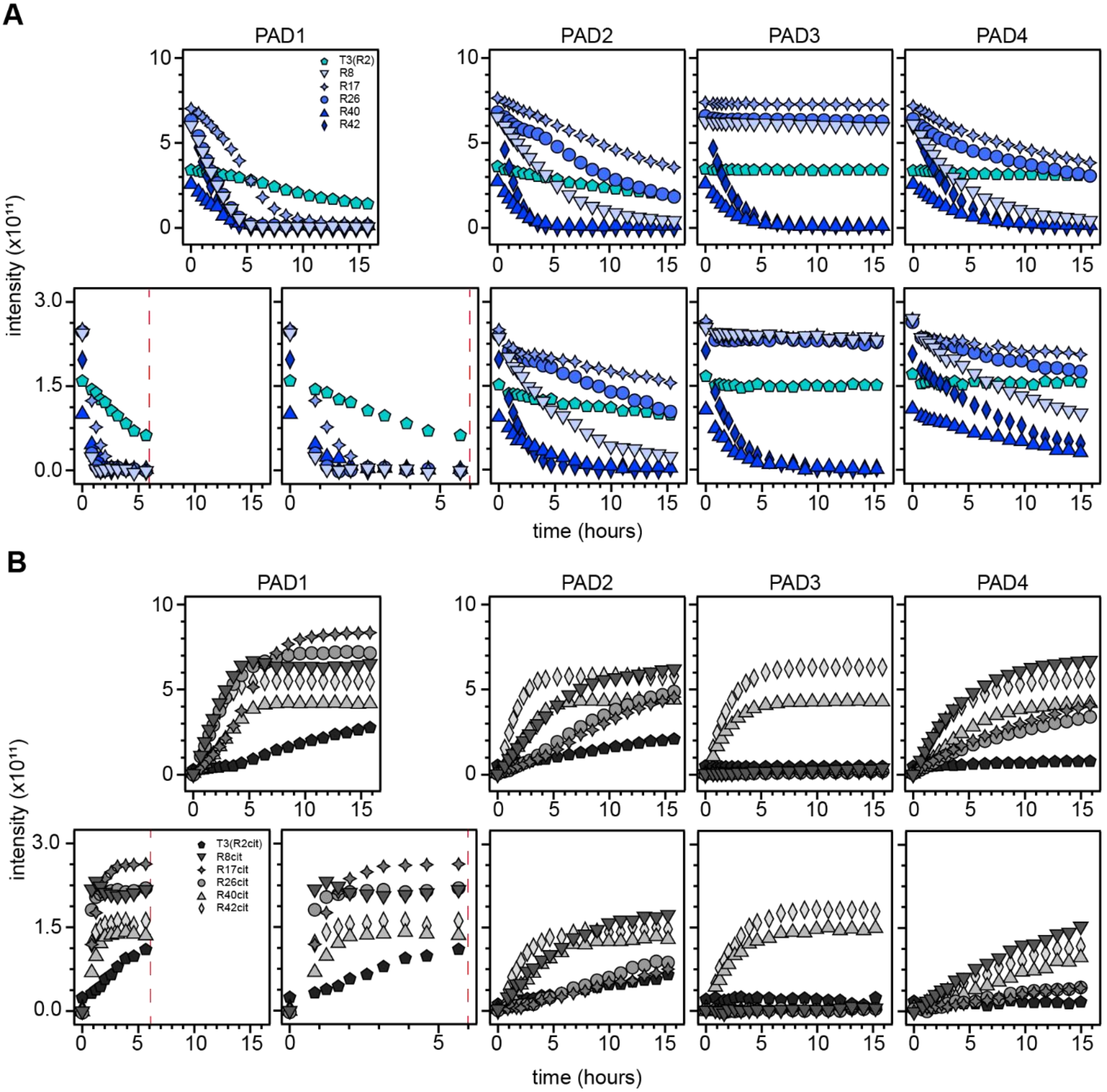


**Figure S4. Comparison of H3 tail peptide NMR assays at different conditions**. Progress curves for H3 tail peptide arginines (**A**) and citrullines (**B**) with PADs 1-4 (left to right) are compared between two sets of conditions: assays from **Figure 3** (**A, B** top) and assays conducted with 50 µM H3 tail peptide (**A, B** bottom). See **Supplementary Methods** for differences in PAD purification. (**A, B** top) Data were collected with 100 µM $^{15}$N-H3(1-44) on an 800 MHz Bruker NMR Spectrometer at 10°C in 20 mM MOPS pH 7.0, 100 mM KCl, 2 mM $CaCl_2$, 0.5 mM TCEP, 0.1 mM EDTA, and 5% $D_2O$. PADs were added to a final concentration of 200 nM. (**A, B** bottom) Data were collected with 50 µM $^{15}$N-H3(1-44) on an 800 MHz Bruker NMR Spectrometer at 10°C in 20 mM MOPS pH 7.0, 150 mM KCl, 10 mM $CaCl_2$, 2 mM DTT, 1 mM EDTA, and 5% $D_2O$. PADs were added to a final concentration of 200 nM. Assays conducted with PAD1 under the 50 µM histone tail peptide conditions are plotted with two x-axis scales to facilitate direct comparison between data sets.

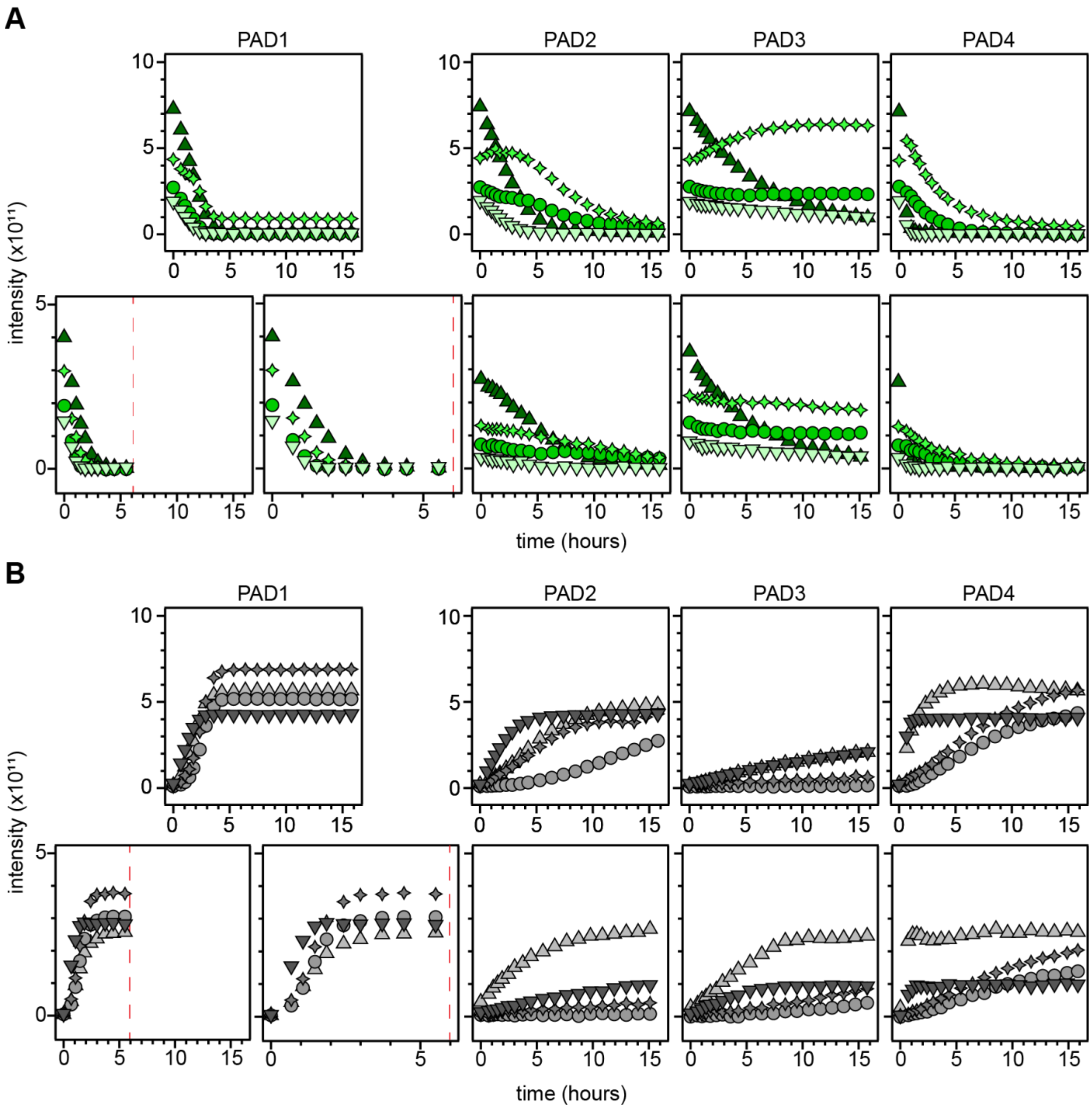


**Figure S5. Comparison of H4 tail peptide NMR assays at different conditions**. Progress curves for H4 tail peptide arginines (**A**) and citrullines (**B**) with PADs 1-4 (left to right) are compared between two sets of conditions: assays from **Figure 3** (**A, B** top) and assays conducted with 50 µM H3 tail peptide (**A, B** bottom). See **Supplementary Methods** for differences in PAD purification. (**A, B** top) Data were collected with 100 µM $^{15}N$-H4(1-33) T30Y on an 800 MHz Bruker NMR Spectrometer at 10°C in 20 mM MOPS pH 7.0, 100 mM KCl, 2 mM $CaCl_2$, 0.5 mM TCEP, 0.1 mM EDTA, and 5% $D_2O$. PADs were added to a final concentration of 200 nM. (**A, B** bottom) Data were collected with 50 µM $^{15}N$-H4(1-33) T30Y on an 800 MHz Bruker NMR Spectrometer at 10°C in 20 mM MOPS pH 7.0, 150 mM KCl, 10 mM $CaCl_2$, 2 mM DTT, 1 mM EDTA, and 5% $D_2O$. PADs were added to a final concentration of 200 nM. Assays conducted with PAD1 under the 50 µM histone tail peptide conditions are plotted with two x-axis scales to facilitate direct comparison between data sets. Assays conducted with PADs 2/4 under the 50 µM histone tail peptide conditions were performed using a room-temperature probe.

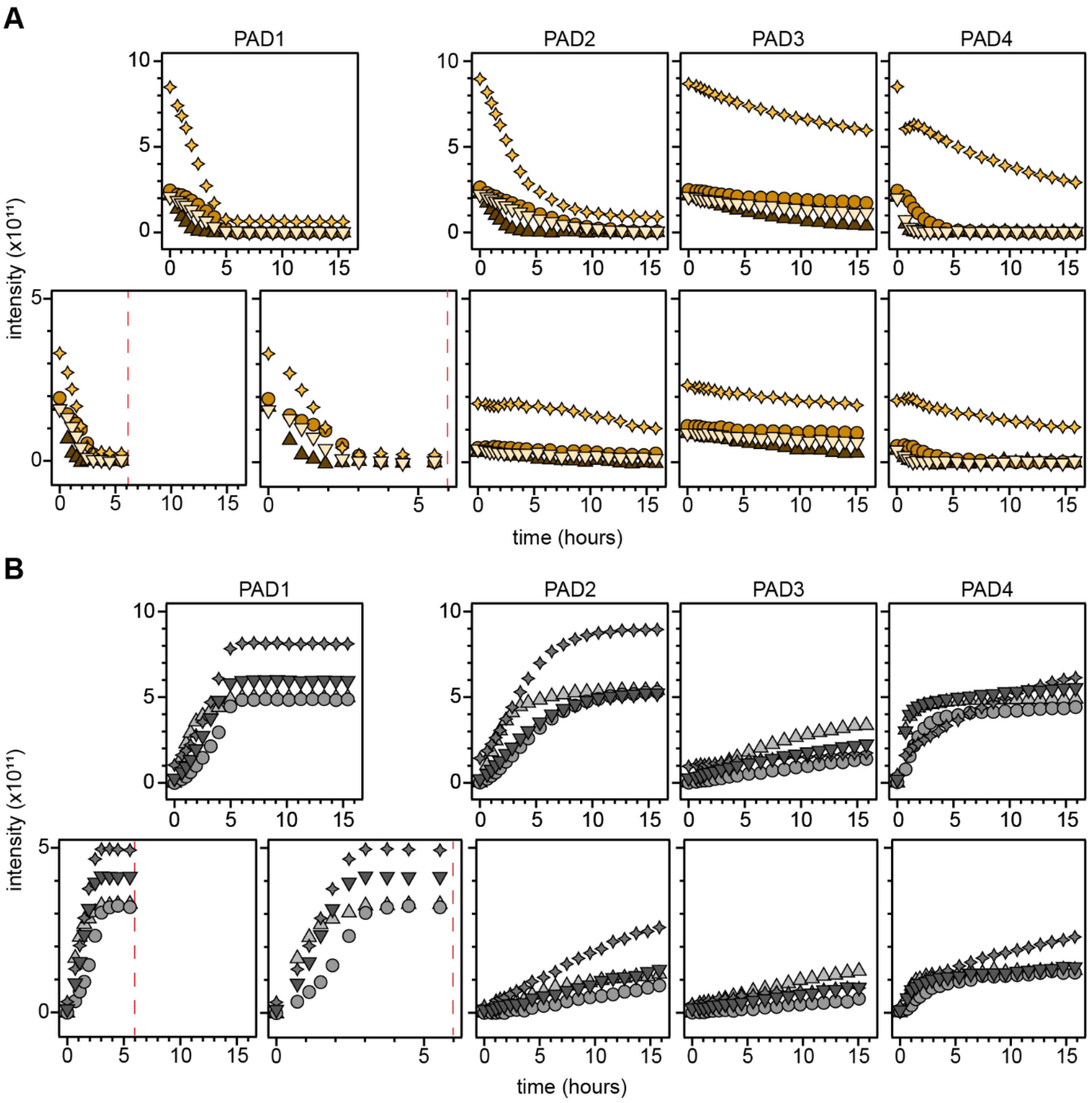


**Figure S6. Comparison of H2A tail peptide NMR assays at different conditions**. Progress curves for H2A tail peptide arginines (**A**) and citrullines (**B**) with PADs 1-4 (left to right) are compared between two sets of conditions: assays from **Figure 3** (**A, B** top) and assays conducted with 50 μM H3 tail peptide (**A, B** bottom). See **Supplementary Methods** for differences in PAD purification. (**A, B** top) Data were collected with 100 μM $^{15}$N-H2A(1-28) F25Y on an 800 MHz Bruker NMR Spectrometer at 10°C in 20 mM MOPS pH 7.0, 100 mM KCl, 2 mM $CaCl_2$, 0.5 mM TCEP, 0.1 mM EDTA, and 5% $D_2O$. PADs were added to a final concentration of 200 nM. (**A, B** bottom) Data were collected with 50 μM $^{15}$N-H2A(1-28) F25Y on an 800 MHz Bruker NMR Spectrometer at 10°C in 20 mM MOPS pH 7.0, 150 mM KCl, 10 mM $CaCl_2$, 2 mM DTT, 1 mM EDTA, and 5% $D_2O$. PADs were added to a final concentration of 200 nM. Assays conducted with PAD1 under the 50 μM histone tail peptide conditions are plotted with two x-axis scales to facilitate direct comparison between data sets. Assays conducted with PADs 2/4 under the 50 μM histone tail peptide conditions were performed using a room-temperature probe.

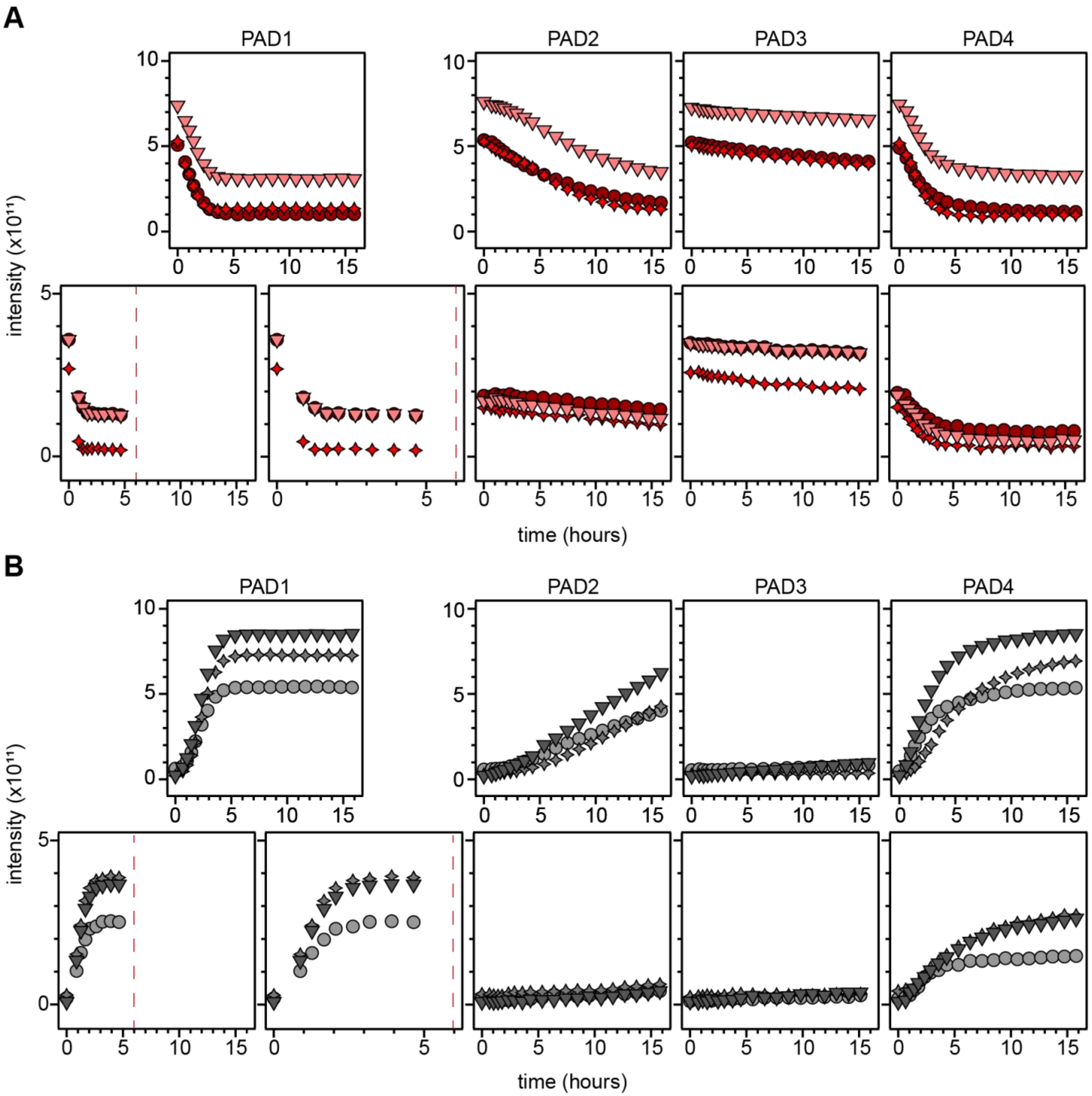


**Figure S7. Comparison of H2B tail peptide NMR assays at different conditions**. Progress curves for H2B tail peptide arginines (**A**) and citrullines (**B**) with PADs 1-4 (left to right) are compared between two sets of conditions: assays from **Figure 3** (**A, B** top) and assays conducted with 50 μM H3 tail peptide (**A, B** bottom). See **Supplementary Methods** for differences in PAD purification. (**A, B** top) Data were collected with 100 μM $^{15}$N-H2B(1-39) on an 800 MHz Bruker NMR Spectrometer at 10°C in 20 mM MOPS pH 7.0, 100 mM KCl, 2 mM $CaCl_2$, 0.5 mM TCEP, 0.1 mM EDTA, and 5% $D_2O$. PADs were added to a final concentration of 200 nM. (**A, B** bottom) Data were collected with 50 μM $^{15}$N-H2B(1-39) on an 800 MHz Bruker NMR Spectrometer at 10°C in 20 mM MOPS pH 7.0, 150 mM KCl, 10 mM $CaCl_2$, 2 mM DTT, 1 mM EDTA, and 5% $D_2O$. PADs were added to a final concentration of 200 nM. Assays conducted with PAD1 under the 50 μM histone tail peptide conditions are plotted with two x-axis scales to facilitate direct comparison between data sets. Assays conducted with PADs 2/4 under the 50 μM histone tail peptide conditions were performed using a room-temperature probe.

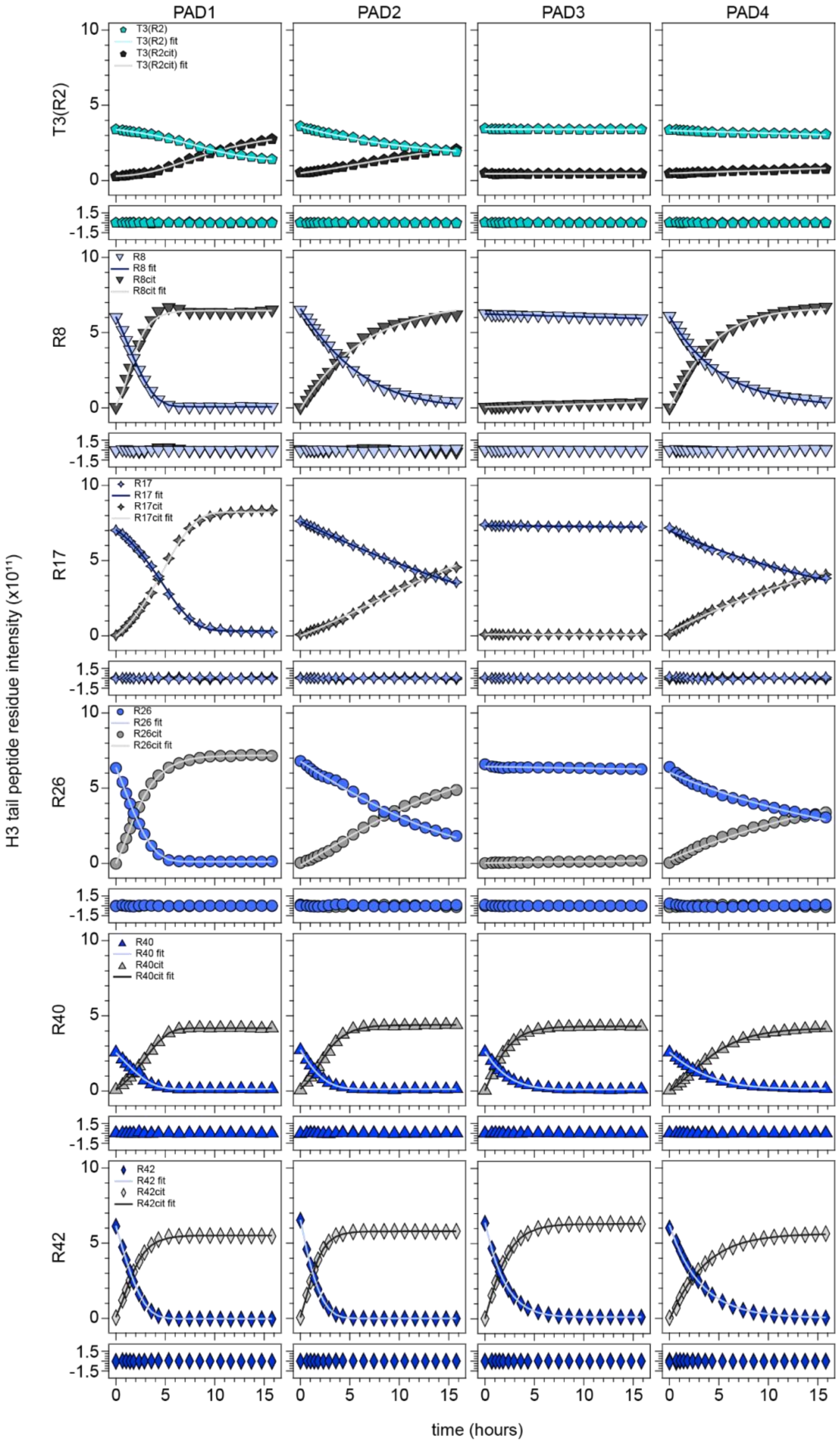


**Figure S8. H3 tail peptide progress curve fits for each PAD.** Top to bottom: progress curve graphs of H3 tail peptide residues T3(R2), R8, R17, R26, R40, and R42 for PADs 1-4 (left to right). The solid line is the fit used for a given residue (see Table S1). Residuals from each fit are shown immediately below the progress curve graph.

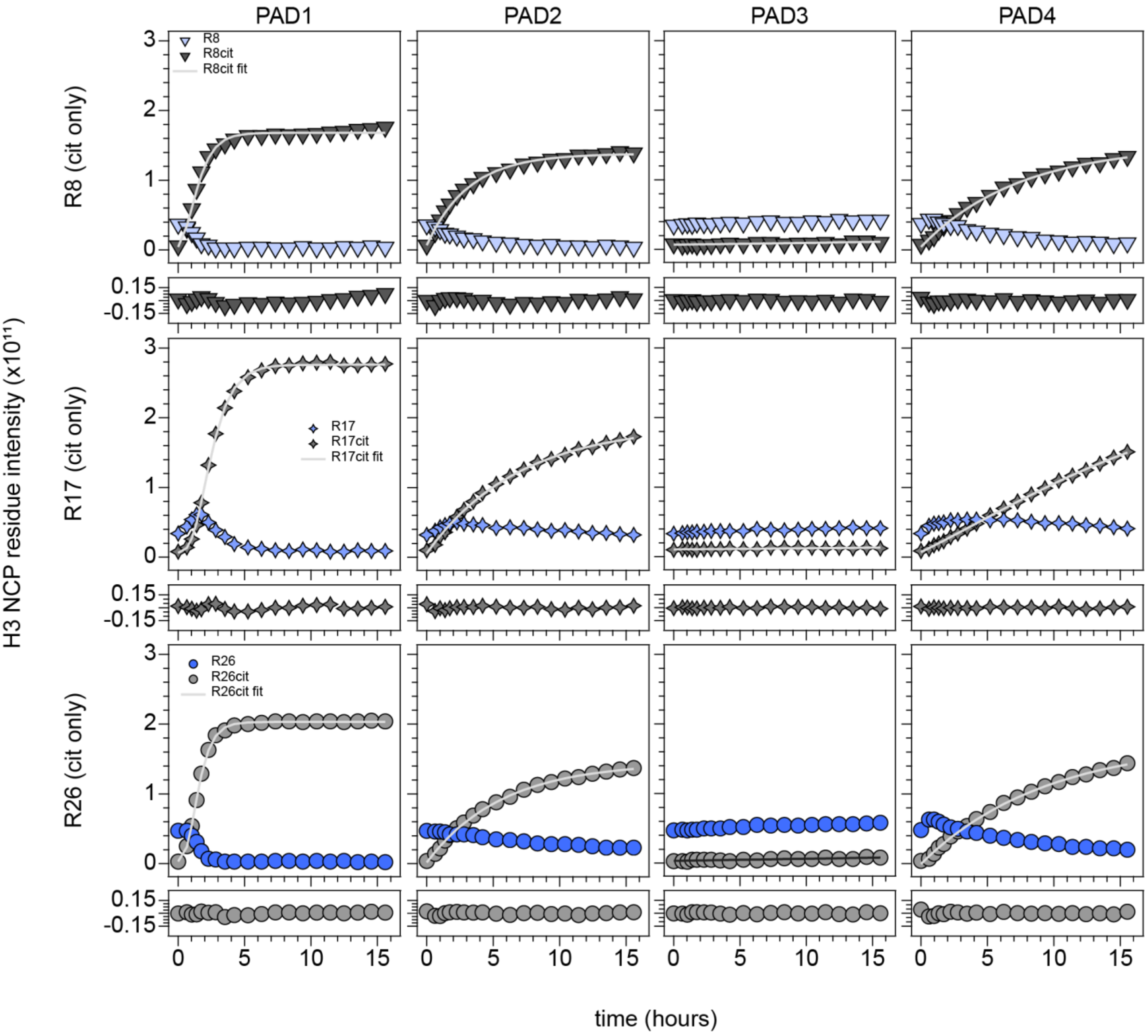


**Figure S9. H3 NCP progress curve fits for each PAD.** Top to bottom: progress curve graphs of H3 NCP residues R8, R17, and R26 for PADs 1-4 (left to right). The solid line is the fit used for a given residue (see Table S1). Residuals from each fit are shown immediately below the progress curve graph.

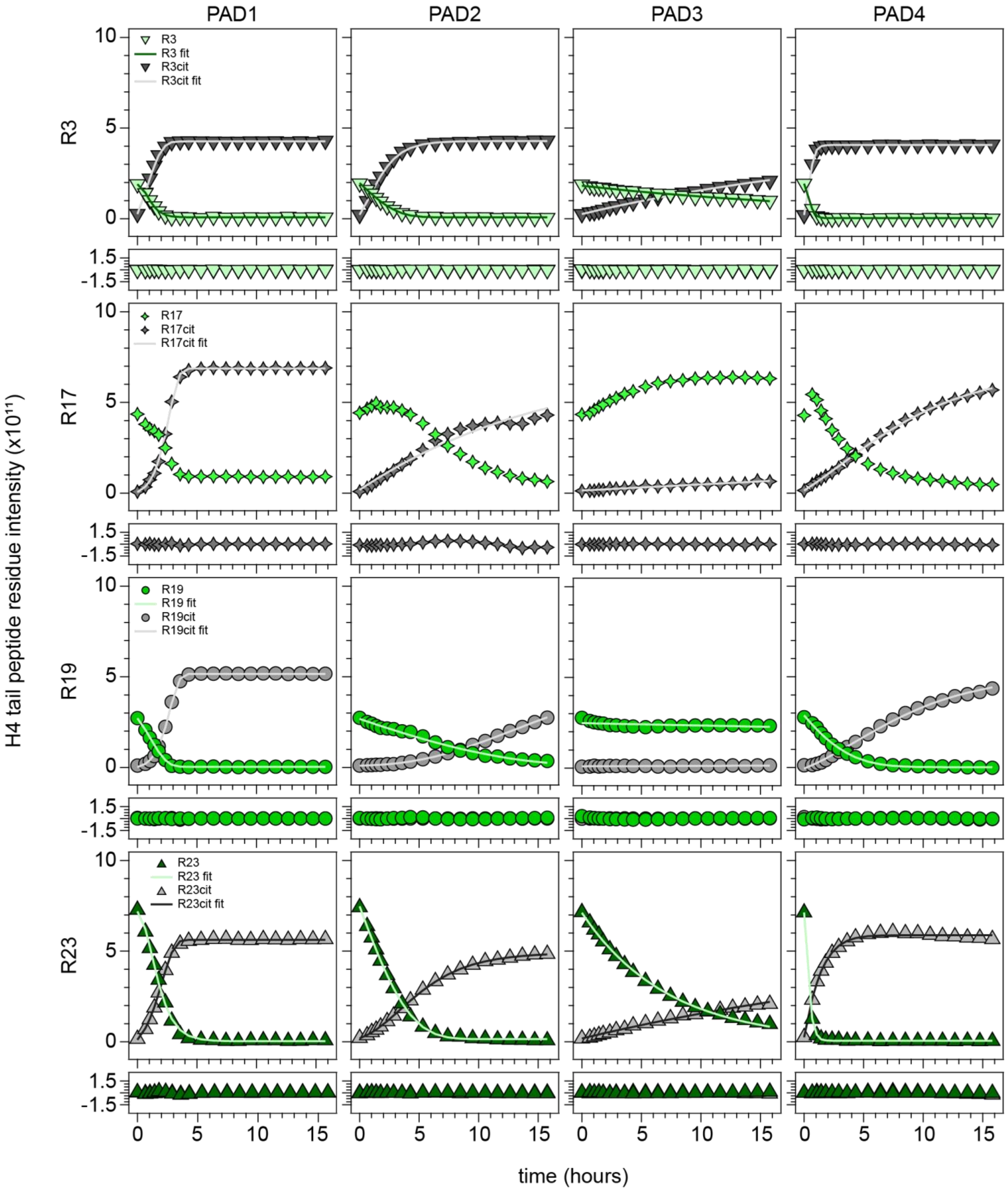


**Figure S10. H4 tail peptide progress curve fits for each PAD.** Top to bottom: progress curve graphs of H4 tail peptide residues R3, R17, R19, and R23 for PADs 1-4 (left to right). The solid line is the fit used for a given residue (see Table S1). Residuals from each fit are shown immediately below the progress curve graph.

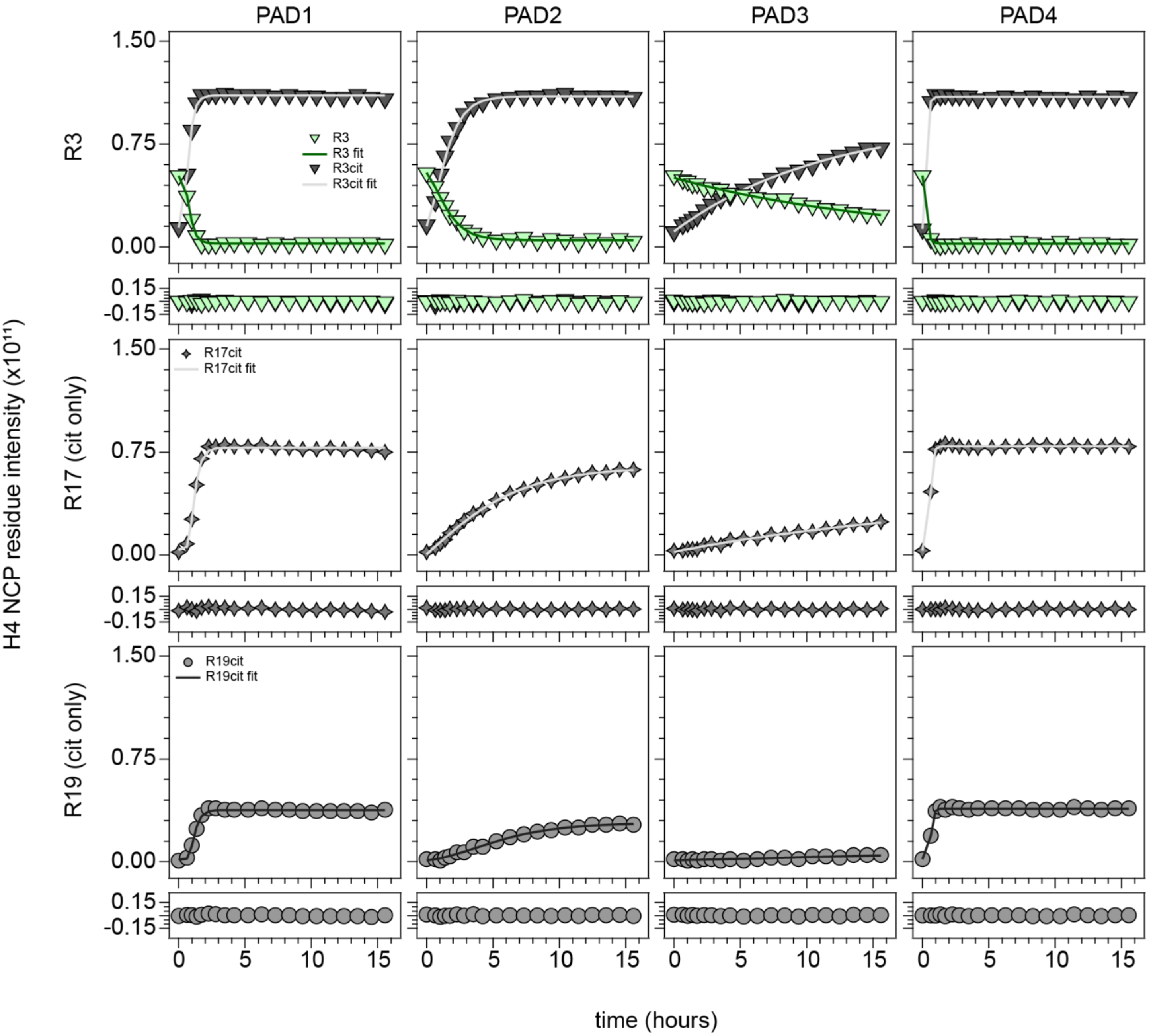


**Figure S11. H4 NCP progress curve fits for each PAD.** Top to bottom: progress curve graphs of H4 NCP residues R3, R17, and R19 for PADs 1-4 (left to right). The solid line is the fit used for a given residue (see Table S1). Residuals from each fit are shown immediately below the progress curve graph.

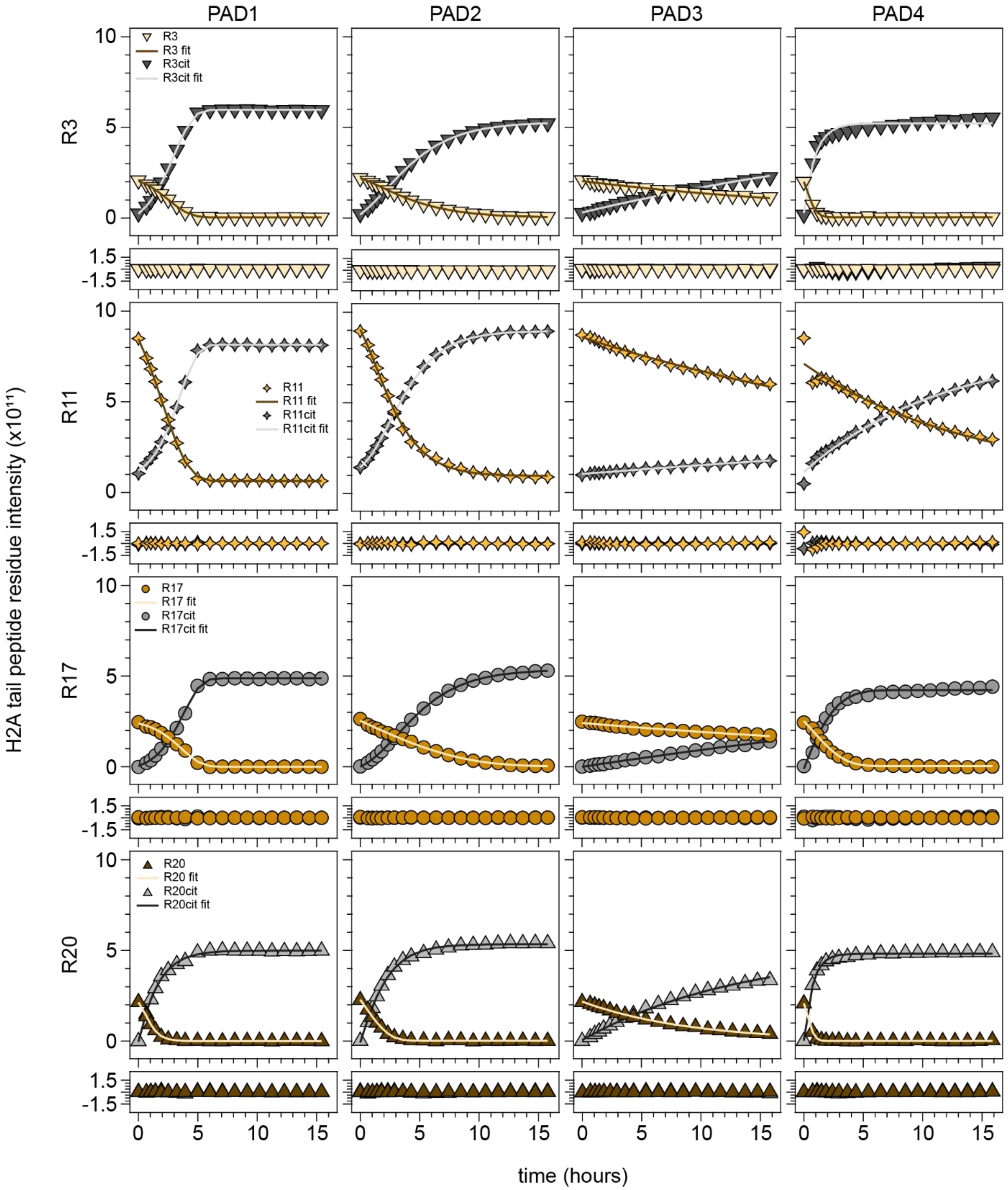


**Figure S12. H2A tail peptide progress curve fits for each PAD.** Top to bottom: progress curve graphs of H2A tail peptide residues R3, R11, R17, and R20 for PADs 1-4 (left to right). The solid line is the fit used for a given residue (see Table S1). Residuals from each fit are shown immediately below the progress curve graph.

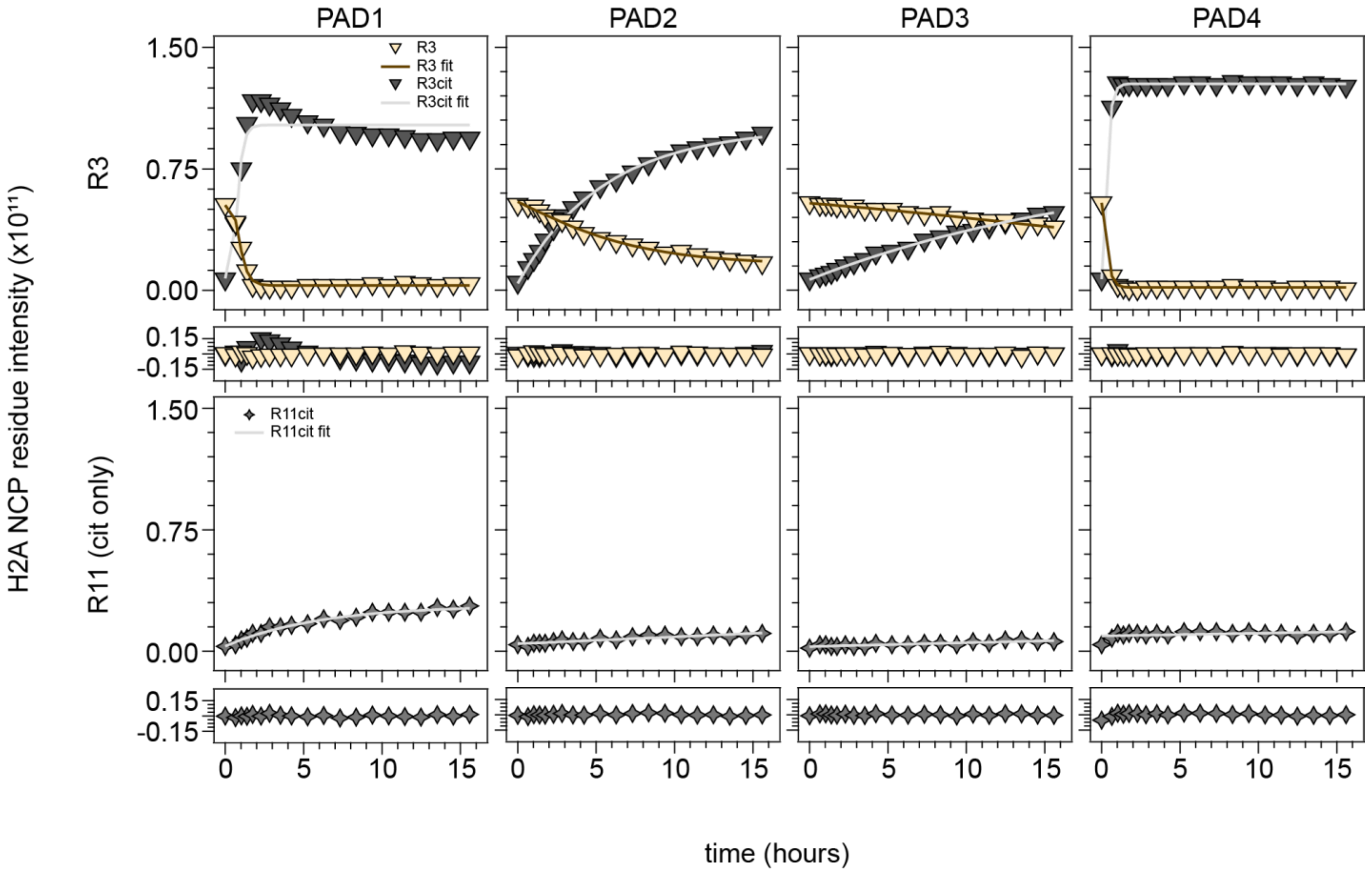


**Figure S13. H2A NCP progress curve fits for each PAD.** Top to bottom: progress curve graphs of H2A NCP residues R3 and R11 for PADs 1-4 (left to right). The solid line is the fit used for a given residue (see Table S1). Residuals from each fit are shown immediately below the progress curve graph.

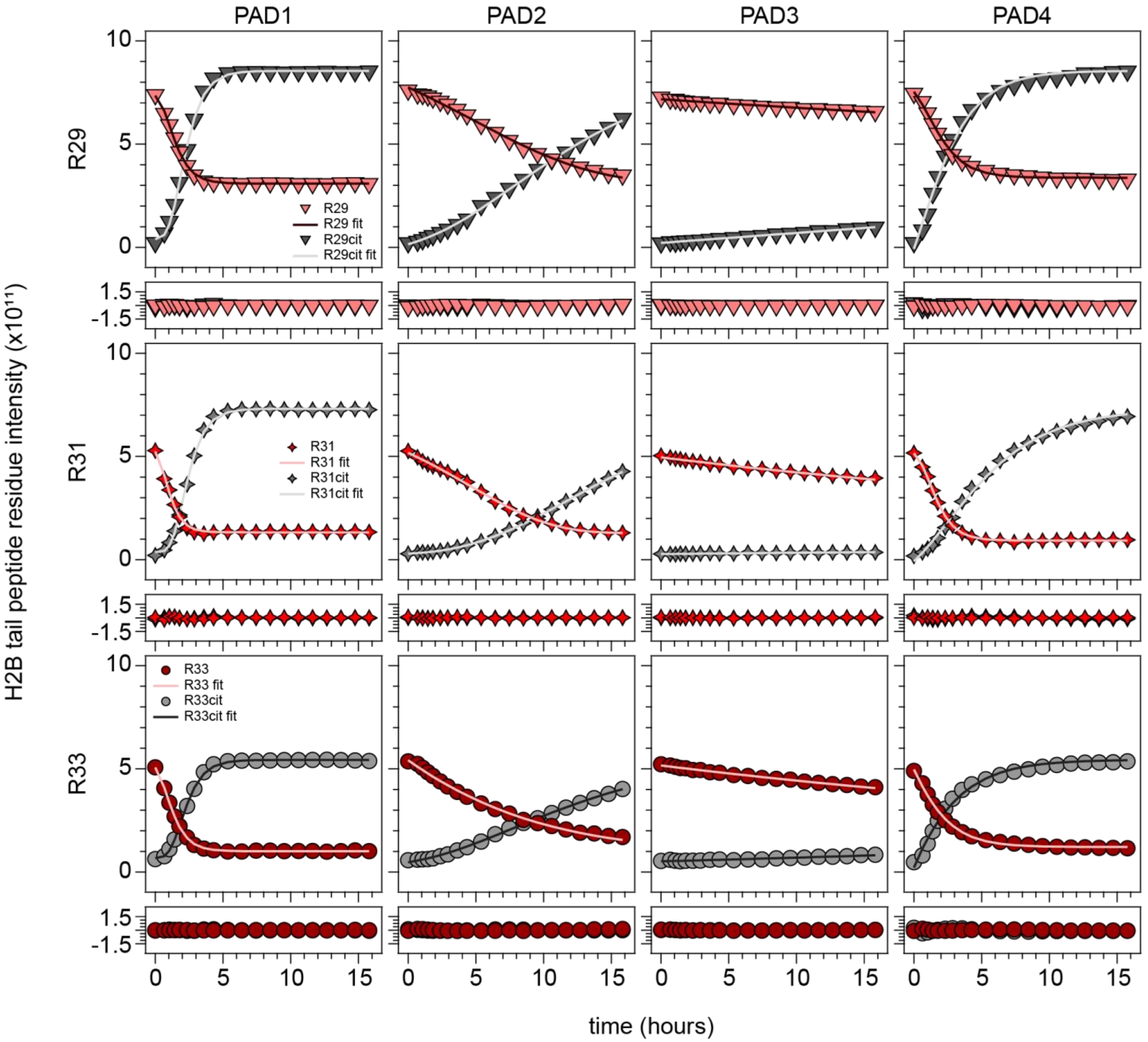


**Figure S14. H2B tail peptide progress curve fits for each PAD.** Top to bottom: progress curve graphs of H2B tail peptide residues R29, R31, and R33 for PADs 1-4 (left to right). The solid line is the fit used for a given residue (see Table S1). Residuals from each fit are shown immediately below the progress curve graph.

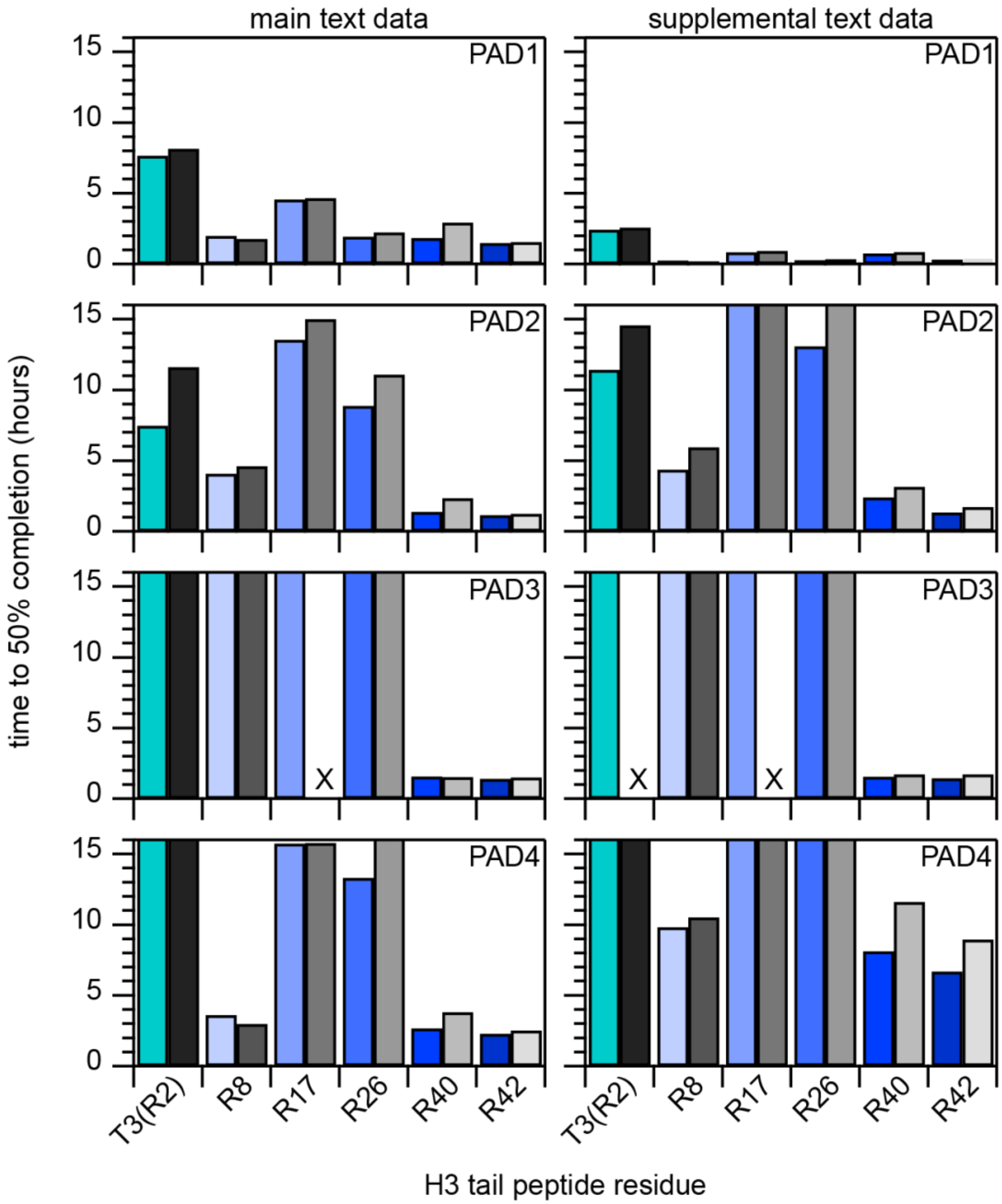


**Figure S15. Comparing H3 tail t50% values between H3 tail NMR assays at different conditions**. Time to 50% completion ($t_{50\%}$) graphs for PADs 1, 2, 3, and 4 (top to bottom) with data from **Figure 11** (left) and **Figure S4-S7.** Both main text and supplemental text data were collected on an 800 MHz Bruker NMR spectrometer with cryoprobe at 10°C. Main text data were collected with 100 µM $^{15}$N-H3(1-44) in 20 mM MOPS pH 7.0, 100 mM KCl, 2 mM $CaCl_2$, 0.5 mM TCEP, 0.1 mM EDTA, and 5% $D_2O$. Supplemental text data were collected with 50 µM $^{15}$N-H3(1-44) in 20 mM MOPS pH 7.0, 150 mM KCl, 10 mM $CaCl_2$, 2 mM DTT, 1 mM EDTA, and 5% $D_2O$. “X” indicates that a $t_{50\%}$ was not determined due to a fitting error.

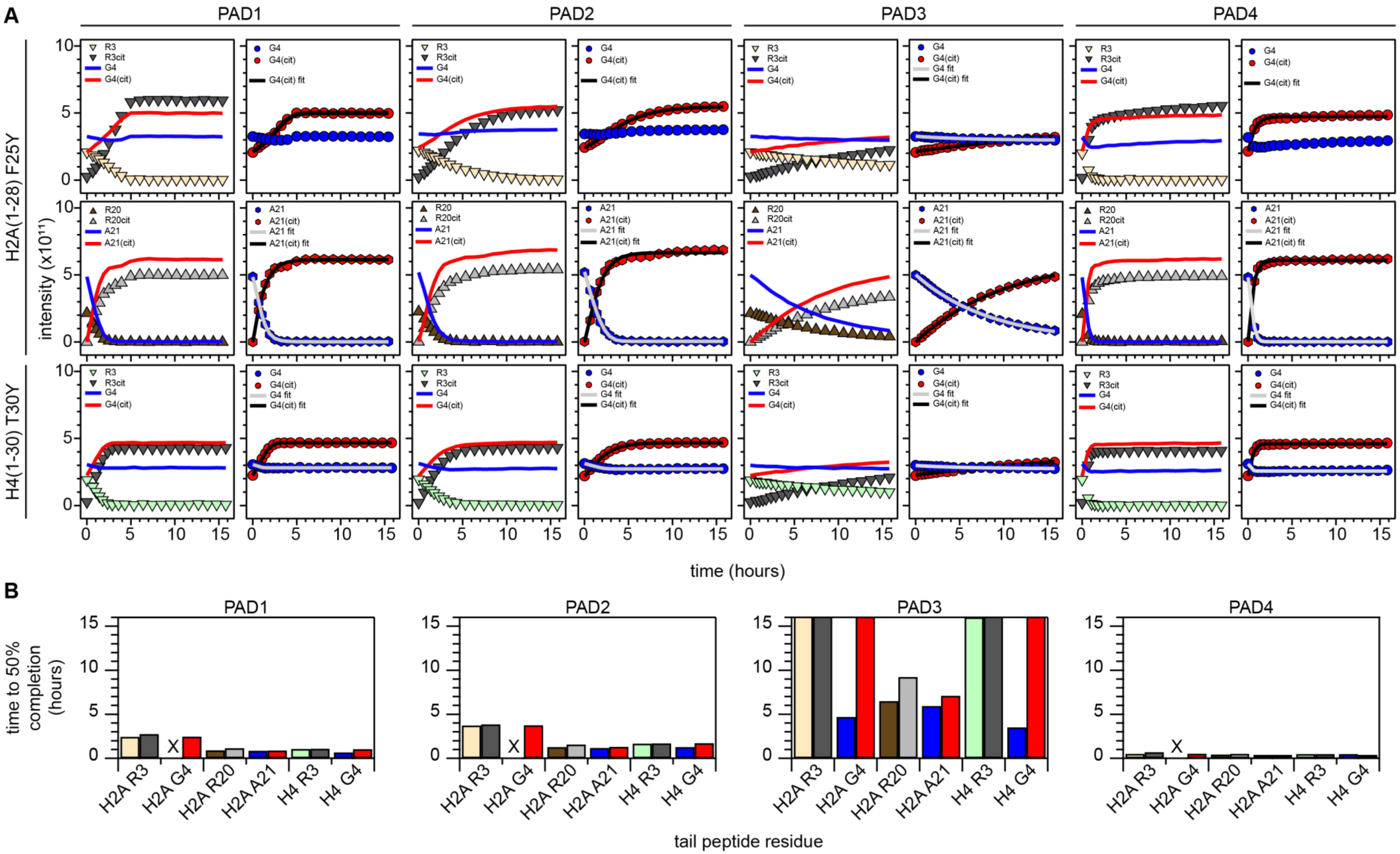


**Figure S16. Validation of utilizing i+1 residues to estimate rates of arginine citrullination.** (**A**, top to bottom): progress curves for tail peptide residues H2A R3, H2A R20, and H4 R3 plotted with their i+1 residue decay (blue) and growth (red) curves (left in each paired plot) and the i+1 residue intensity plots with fit curves (right in each paired plot) for PADs 1-4. (**B**) Time to 50% completion ($t_{50\%}$) graphs for PADs 1, 2, 3, and 4 (left to right) with select histone arginines and their i+1 residues. An “X” indicates failure of the curve fitting.

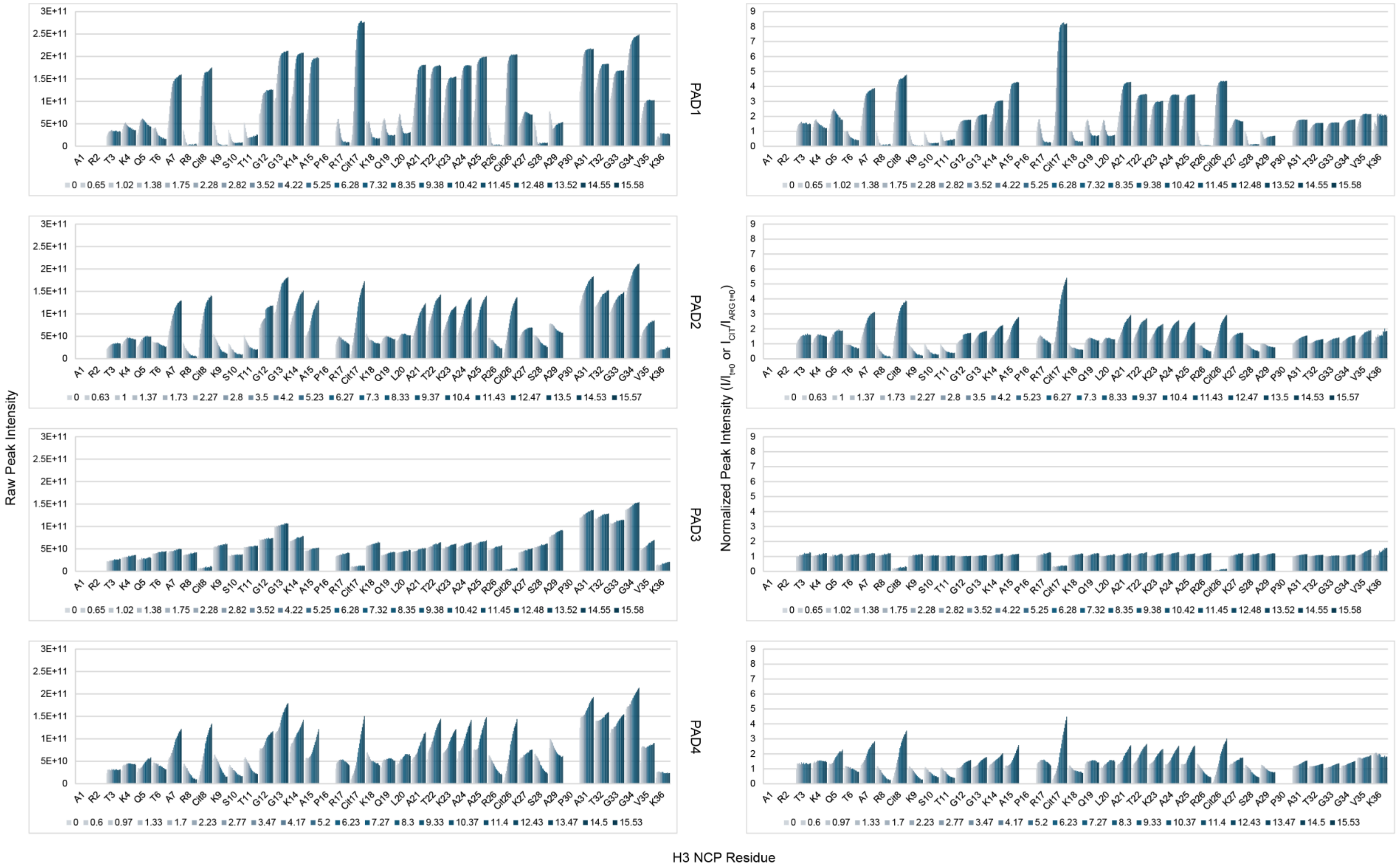


**Figure S17. Intensities of observable H3 tail residues in the NCP.** Raw peak intensities (left) and scaled intensities ($I/I_0$ or $I_{CIT}/I_{ARG\ t=0}$) (right) of H3 tail residues in $^{15}$N-H3-NCP treated with PADs 1-4 (top to bottom). The intensities of most non-arginine residues increase. Some residues (i.e., T6, K9, S10, T11, K18, Q19, L20, S28, S29) show a decrease in intensity with some of the PADs because they sense citrullination of nearby residues. These residues have a distinct peak for the citrullinated state, and the intensity of the uncitrullinated state is shown here.

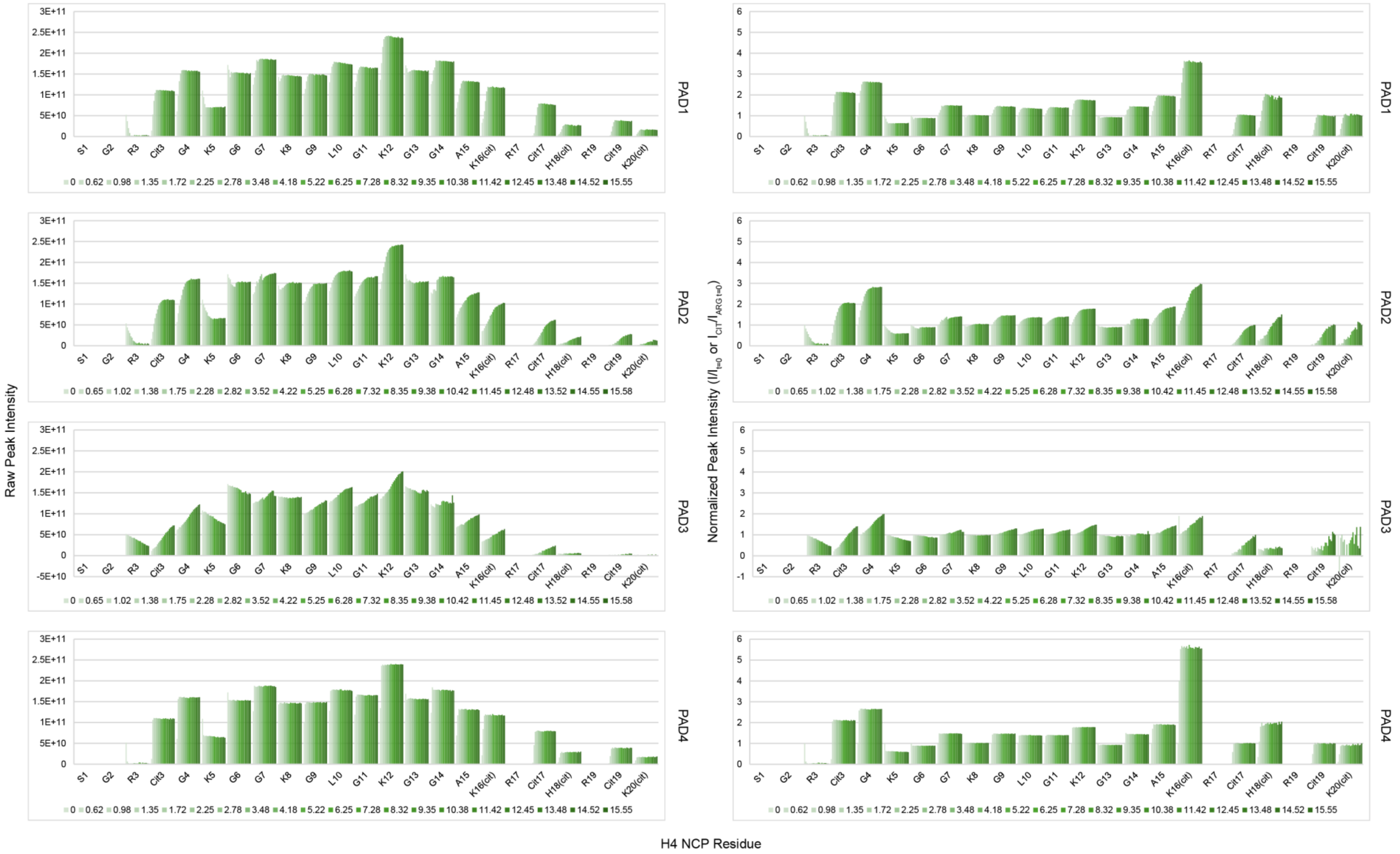


**Figure S18. Intensities of observable H4 tail residues in the NCP.** Raw peak intensities (left) and scaled intensities ($I/I_0$ or $I_{CIT}/I_{ARG\ t=0}$) (right) of H4 tail residues in $^{15}$N-H4-NCP treated with PADs 1-4 (top to bottom). The intensities of most non-arginine residues increase. A few residues (i.e., K5, G6) show a decrease in intensity, likely because they sense citrullination of nearby residues.

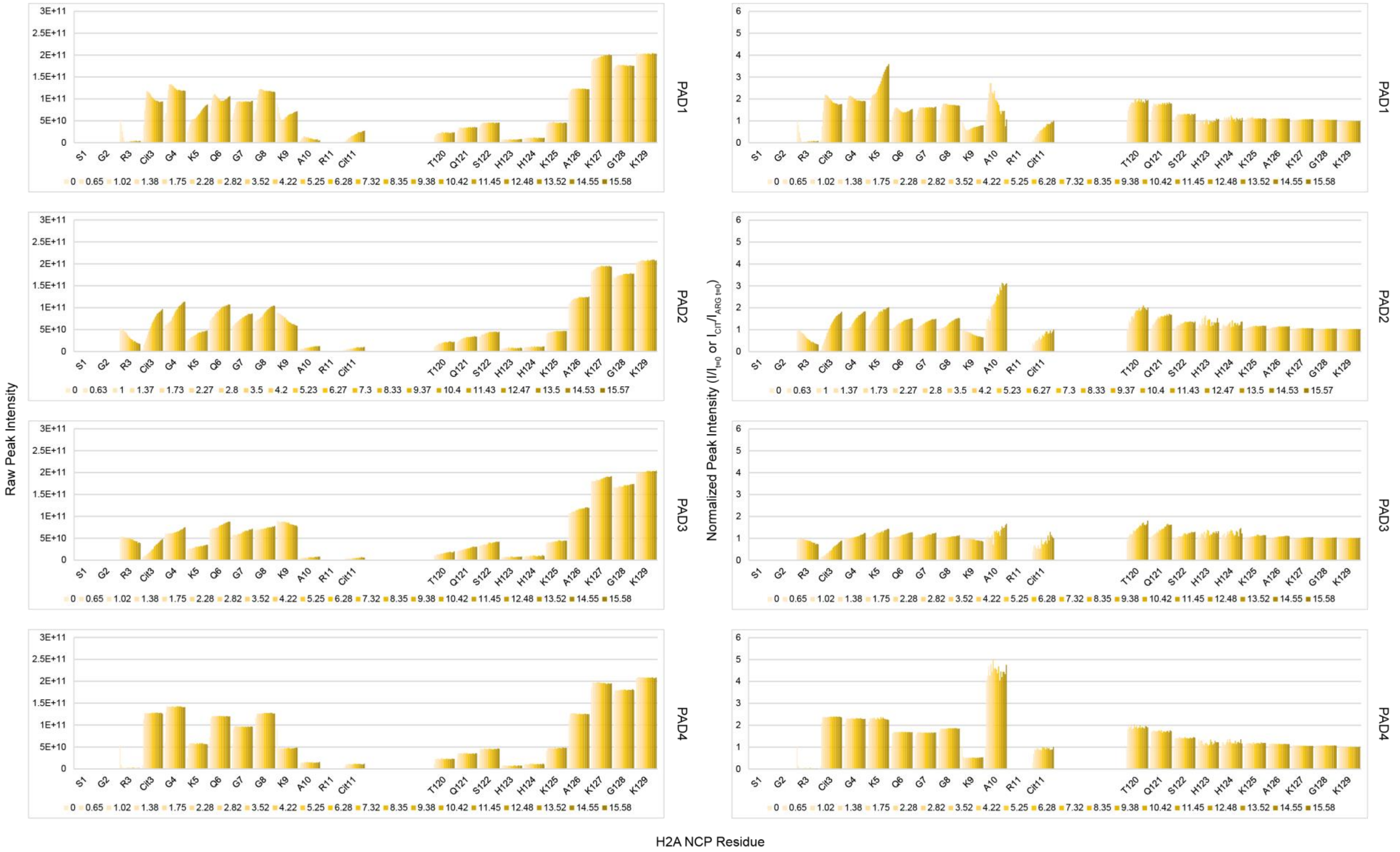


**Figure S19. Intensities of observable H2A tail residues in the NCP.** Raw peak intensities (left) and scaled intensities ($I/I_0$ or $I_{CIT}/I_{ARG\ t=0}$) (right) of H2A tail residues in $^{15}$N-H2A-NCP treated with PADs 1-4 (top to bottom). The intensities of most non-arginine residues increase. K9 shows a decrease in intensity, likely because it senses citrullination of nearby residues.

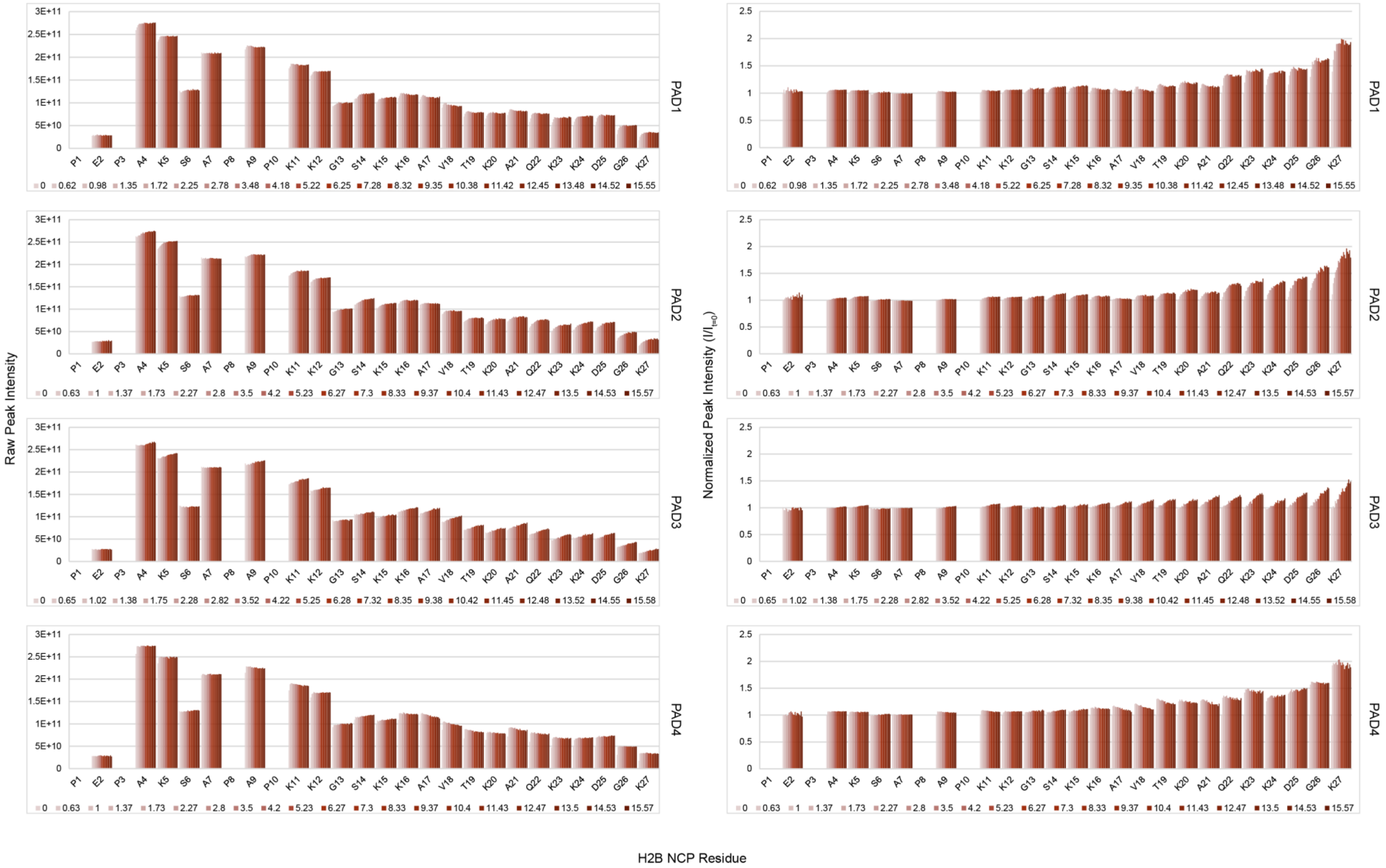


**Figure S20. Intensities of observable H2B tail residues in the NCP.** Raw peak intensities (left) and scaled intensities ($I/I_0$) (right) of H2B tail residues in $^{15}$N-H2B-NCP treated with PADs 1-4 (top to bottom). The intensities of most residues increase, with the relative increase getting larger toward the C-terminal end of the tail

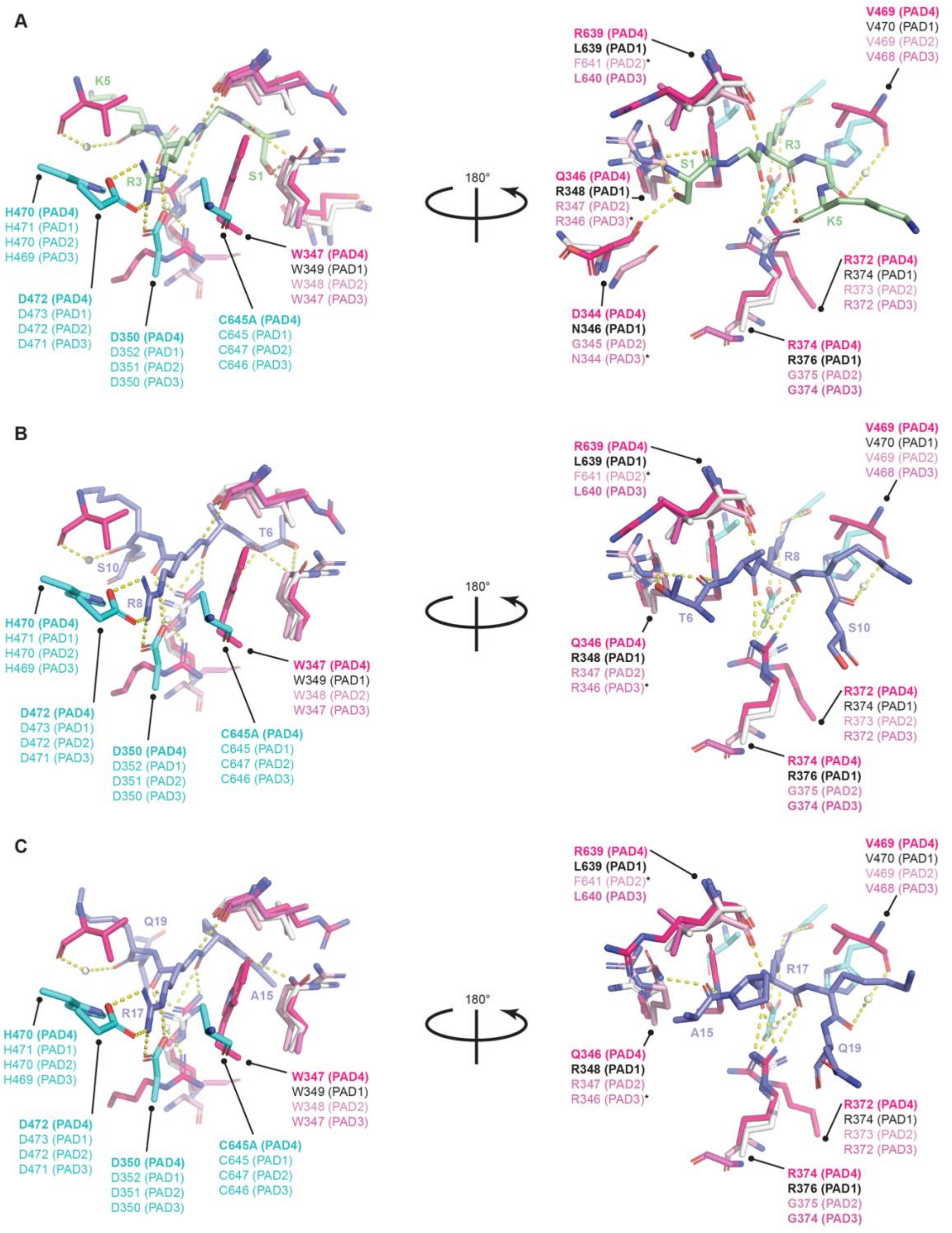


**Figure S21. PAD structural alignment**. X-ray crystal structures of PAD1 (PDB: 5HP5), PAD2 (PDB: 4N2C), and PAD3 (PDB: 7DAN) were aligned in PyMOL to PAD4 structures in complex with either (**A**) H4 tail peptide containing R3 (PDB: 2DEY), (**B**) H3 tail peptide containing R8 (PDB: 2DEW), or (**C**) H3 tail peptide containing R17 (PDB: 2DEX). These structural alignments highlight the locations of PAD4 residues that interact with histone substrate and their corresponding positions in PADs 1-3. Yellow dashed lines represent PAD4-histone tail interactions as found by Arita et. al. (6). Residues labeled in bold are visible; some homolog residues are omitted for visual clarity. Residues are colored as follows: catalytic residues in cyan, non-catalytic residues in shades of pink, and substrate residues in green and blue for histones H4 and H3, respectively. (*) indicates partial or fully unresolved residues in their respective structures. Small, white spheres are water molecules involved in water-mediated interactions.

```
PADI4_HUMAN      MAQGTLIRVTPEQPTHAVCVLGTLTQLDICSSAPEDCTSFSINASPGVVVDIAHGPP-AK      59
PADI1_HUMAN      MAPKRVVQLSLKMPTHAVCVVGVEAHVDIHSDVPKGANSFRVSGSSGVEVFMVYNRT-RV      59
PADI2_HUMAN      MLRERTVRLQYGSRVEAVYVLGTYLWTDVYSAAPAGAQTFSLKHSEHVWVEVVRDGEAEE      60
PADI3_HUMAN      MSLQRIVRVSLEHPTSAVCVAGVETLVDIYGSVPEGTEMFEVYGTPGVDIYISPNME-RG      59
                 *     :::     . ** * *.    *: . .* .   * :  :  * : :  .

PADI4_HUMAN      KKSTGSSTWPLDPGVEVTLTMKVASGSTGDQKVQISYYGPKTP--PVKALLYLTGVEISL      117
PADI1_HUMAN      KEPIGKARWPLDTDADMVVSVGTASKELKDFKVRVSYFGEQEDQALGRSVLYLTGVDISL      119
PADI2_HUMAN      VATNGKQRWLLSPSTTLRVTMSQASTEASSDKVTVNYYDEEGSIPIDQAGLFLTAIEISL      120
PADI3_HUMAN      RERADTRRWRFDATLEIIVVMNSPSNDLNDSHVQISYHSSHEPLPLAYAVLYLTCVDISL      119
                     ..  * :.    : : :   * .  . :* :.*.. .       : *:** ::***

PADI4_HUMAN      CADITRTGKVKPTRAVKDQRTWTWGPCGQGAILLVNCDRDNLESSAMDCEDDEVLDSEDL      177
PADI1_HUMAN      EVDTGRTGKVK--RSQGDKKTWRWGPEGYGAILLVNCDRDNHRSAEPDLTHSWLMSLADL      177
PADI2_HUMAN      DVDADRDGVVE--KNNPKKASWTWGPEGQGAILLVNCDRETPWLPKEDCRDEKVYSKEDL      178
PADI3_HUMAN      DCDLNCEGRQD--RNFVDKRQWVWGPSGYGGILLVNCDRDDPSCDVQDNCDQHVHCLQDL      177
                   *    *  .  :   .:  * *** * *.********:       *  .. :    **

PADI4_HUMAN      QDMSLMTLSTKTPKDFFTNHTLVLHVARSEMDKVRVFQATRGKL-SSKCSVVLGPKWPSH      236
PADI1_HUMAN      QDMSPMLLSCNGPDKLFDSHKLVLNVPFSDSKRVRVFCARGGNS-LSDYKQVLGPQCLSY      236
PADI2_HUMAN      KDMSQMILRTKGPDRLPAGYEIVLYISMSDSDKVGVFYVENPFF-GQRYIHILGRRKLYH      237
PADI3_HUMAN      EDMSVMVLRTQGPAALFDDHKLVLHTSSYDAKRAQVFHICGPEDVCEAYRHVLGQDKVSY      237
                 :*** * *  : *  :  .: :**     : .:. **         .    :**     :

PADI4_HUMAN      YLMVPGGKHNMDFYVEALAFPDTDFPGLITLTISLLDTSNLELPEAVVFQDSVVFRVAPW      296
PADI1_HUMAN      EVERQPGEQEIKFYVEGLTFPDADFLGLVSLSVSLVDPG--TLPEVTLFTDTVGFRMAPW      294
PADI2_HUMAN      VVKYTGGSAELLFFVEGLCFPDEGFSGLVSIHVSLLEYMAQDIPLTPIFTDTVIFRIAPW      297
PADI3_HUMAN      EVPRLHGDE-ERFFVEGLSFPDAGFTGLISFHVTLLDDSNEDFSASPIFTDTVVFRVAPW      296
                  :    *.    *:**.* *** .* **::: ::*::     :    :* *:* **:***

PADI4_HUMAN      IMTPNTQPPQEVYACSIFE----NEDFLKSVTTLAMKAKCKLTICPEEENMDDQWMQDEM      352
PADI1_HUMAN      IMTPNTQPPEELYVCRVMDTHGSNEKFLEDMSYLTLKANCKLTICPQVENRNDRWIQDEM      354
PADI2_HUMAN      IMTPNILPPVSVFVCCMKD----NYLFLKEVKNLVEKTNCELKVCFQYLNRGDRWIQDEI      353
PADI3_HUMAN      IMTPSTLPPLEVYVCRVRN----NTCFVDAVAELARKAGCKLTICPQAENRNDRWIQDEM      352
                 ****.  ** .::.* : :    *  *:. :  *. *: *:*.:* :  * .*:*:***:

PADI4_HUMAN      EIGYIQAPHKTLPVVFDSPRNRGLKEFPIKRVMGPDFGYVTRGPQTGGISGLDSFGNLEV      412
PADI1_HUMAN      EFGYIEAPHKSFPVVFDSPRNRGLKDFPYKRILGPDFGYVTREIPLPGPSSLDSFGNLDV      414
PADI2_HUMAN      EFGYIEAPHKGFPVVLDSPRDGNLKDFPVKELLGPDFGYVTREPLFESVTSLDSFGNLEV      413
PADI3_HUMAN      ELGYVQAPHKTLPVVFDSPRNGELQDFPYKRILGPDFGYVTREPRDRSVSGLDSFGNLEV      412
                 *:**::**** :***:****:  *::** *.::*********     . :.*******:*

PADI4_HUMAN      SPPVTVRGKEYPLGRILFGDSCYPSNDSRQMHQALQDFLSAQQVQAPVKLYSDWLSVGHV      472
PADI1_HUMAN      SPPVTVGGTEYPLGRILIGSS-FPKSGGRQMARAVRNFLKAQQVQAPVELYSDWLSVGHV      473
PADI2_HUMAN      SPPVTVNGKTYPLGRILIGSS-FPLSGGRRMTKVVRDFLKAQQVQAPVELYSDWLTVGHV      472
PADI3_HUMAN      SPPVVANGKEYPLGRILIGGN-LPGSSGRRVTQVVRDFLHAQKVQPPVELFVDWLAVGHV      471
                 ****.. *. *******:*..  * ...*:: :.:::** **:** **:*: ***:****

PADI4_HUMAN      DEFLSFVPAPDRKGFRLLLASPRSCYKLFQEQQNEGHGEALLFEGIKKKK---QQKIKNI      529
PADI1_HUMAN      DEFLTFVPTSDQKGFRLLLASPSACLKLFQEKKEEGYGEAAQFDGLKHQA---KRSINEM      530
PADI2_HUMAN      DEFMSFVPIPGTKKFLLLMASTSACYKLFREKQKDGHGEAIMFKGLGGMS-SKRITINKI      531
PADI3_HUMAN      DEFLSFVPAPDGKGFRMLLASPGACFKLFQEKQKCGHGRALLFQGVVDDEQVKTISINQV      531
                 ***::***  . * * :*:**  :* ***:*::: *:*.*  *.*:         .*:::

PADI4_HUMAN      LSNKTLREHNSFVERCIDWNRELLKRELGLAESDIIDIPQLFKLKEFSKAEAFFPNMVNM      589
PADI1_HUMAN      LADRHLQRDNLHAQKCIDWNRNVLKRELGLAESDIVDIPQLFFLKNF-YAEAFFPDMVNM      589
PADI2_HUMAN      LSNESLVQENLYFQRCLDWNRDILKKELGLTEQDIIDLPALFKMDEDHRARAFFPNMVNM      591
PADI3_HUMAN      LSNKDLINYNKFVQSCIDWNREVLKRELGLAECDIIDIPQLFKTERK-KATAFFPDLVNM      590
                 *::. * . * . : *:****::**:****:* **:*:* **  ..   * ****::***

PADI4_HUMAN      LVLGKHLGIPKPFGPVINGRCCLEEKVCSLLEPLGLQCTFINDFFTYHIRHGEVHCGTNV      649
PADI1_HUMAN      VVLGKYLGIPKPYGPIINGRCCLEEKVQSLLEPLGLHCIFIDDYLSYHELQGEIHCGTNV      649
PADI2_HUMAN      IVLDKDLGIPKPFGPQVEEECCLEMHVRGLLEPLGLECTFIDDISAYHKFLGEVHCGTNV      651
PADI3_HUMAN      LVLGKHLGIPKPFGPIINGCCCLEEKVRSLLEPLGLHCTFIDDFTPYHMLHGEVHCGTNV      650
                 :**.* ******:** ::  **** :* .*******.* **:*   **   **:******

PADI4_HUMAN      RRKPFSFKWWNMVP      663
PADI1_HUMAN      RRKPFPFKWWNMVP      663
PADI2_HUMAN      RRKPFTFKWWHMVP      665
PADI3_HUMAN      CRKPFSFKWWNMVP      664
                  **** ****:***
```

**Figure S22. PAD sequence alignment**. PADs 1-3 (UniProt sequences Q9ULC6, Q9Y2J8, Q9ULW8, respectively) were aligned to PAD4 (UniProt Q9UM07) in UniProt Clustal Omega. Highlighted residues are PAD4 residues that were found to interact with histone tail substrates and their aligned equivalent positions in PADs 1-3. Residues in red are conserved and involved with substrate catalysis. The following notations indicate conservation status: (*) fully conserved, (:) conserved between groups with strongly similar properties, (.) conserved between groups with weakly similar properties, ( ) not conserved.

| Substrate | | Function used for $t_{50\%}$ calculations | | | |
|---|---|---|---|---|---|
| | | PAD1 | PAD2 | PAD3 | PAD4 |
| H2A tail peptide | R3 | Gompertz | Gompertz | exponential | Gompertz |
| | Cit3 | Gompertz | Gompertz | exponential | Gompertz |
| | R11 | Gompertz | Gompertz | exponential | exponential |
| | Cit11 | Gompertz | Gompertz | exponential | exponential |
| | R17 | Gompertz | Gompertz | exponential | Gompertz |
| | Cit17 | Gompertz | Gompertz | Gompertz | Gompertz |
| | R20 | Gompertz | Gompertz | exponential | Gompertz |
| | Cit20 | Gompertz | Gompertz | exponential | Gompertz |
| H2B tail peptide | R29 | Gompertz | Gompertz | exponential | Gompertz |
| | Cit29 | Gompertz | Gompertz | Gompertz | Gompertz |
| | R31 | Gompertz | Gompertz | exponential | Gompertz |
| | Cit31 | Gompertz | Gompertz | exponential | Gompertz |
| | R33 | Gompertz | exponential | exponential | Gompertz |
| | Cit33 | Gompertz | Gompertz | Gompertz | Gompertz |
| H3 tail peptide | T3(R2) | Gompertz | Gompertz | linear | linear |
| | T3(Cit2) | Gompertz | Gompertz | linear | linear |
| | R8 | Gompertz | Gompertz | linear | exponential |
| | Cit8 | Gompertz | exponential | linear | Gompertz |
| | R17 | Gompertz | Gompertz | linear | exponential |
| | Cit17 | Gompertz | Gompertz | linear | exponential |
| | R26 | Gompertz | Gompertz | linear | exponential |
| | Cit26 | Gompertz | Gompertz | linear | exponential |
| | R40 | Gompertz | Gompertz | Gompertz | Gompertz |
| | Cit40 | Gompertz | Gompertz | Gompertz | Gompertz |
| | R42 | Gompertz | Gompertz | Gompertz | exponential |
| | Cit42 | Gompertz | Gompertz | Gompertz | Gompertz |
| H4 tail peptide | R3 | Gompertz | Gompertz | exponential | Gompertz |
| | Cit3 | Gompertz | Gompertz | exponential | Gompertz |
| | R17 | - | - | - | - |
| | Cit17 | Gompertz | exponential | exponential | Gompertz |
| | R19 | Gompertz | Gompertz | linear | Gompertz |
| | Cit19 | Gompertz | Gompertz | linear | Gompertz |
| | R23 | Gompertz | Gompertz | exponential | Gompertz |
| | Cit23 | Gompertz | Gompertz | exponential | Gompertz |
| H2A NCP | R3 | Gompertz | Gompertz | Gompertz | Gompertz |
| | Cit3 | Gompertz | exponential | exponential | Gompertz |
| | R11 | - | - | - | - |
| | Cit11 | Gompertz | linear | linear | linear |
| H3 NCP | T3(R2) | - | - | - | - |
| | T3(Cit2) | - | - | - | - |
| | R8 | - | - | - | - |
| | Cit8 | Gompertz | exponential | linear | exponential |
| | R17 | - | - | - | - |
| | Cit17 | Gompertz | exponential | linear | Gompertz |
| | R26 | - | - | - | - |
| | Cit26 | Gompertz | exponential | linear | exponential |
| H4 NCP | R3 | Gompertz | Gompertz | exponential | Gompertz |
| | Cit3 | Gompertz | Gompertz | exponential | Gompertz |
| | R17 | - | - | - | - |
| | Cit17 | Gompertz | Gompertz | exponential | Gompertz |
| | R19 | - | - | - | - |
| | Cit19 | Gompertz | Gompertz | linear | Gompertz |

**Table S1. Fit functions utilized for $t_{50\%}$ calculations for each PAD-histone arginine and citrulline residue.**

| Substrate | | $t_{50\%}$ values (hours) | | | |
|---|---|---|---|---|---|
| | | PAD1 | PAD2 | PAD3 | PAD4 |
| H2A tail peptide | R3 | 2.4 | 3.7 | 17 | 0.52 |
| | Cit3 | 2.8 | 3.9 | 25 | 0.70 |
| | R11 | 2.2 | 2.6 | 25 | 7.1 |
| | Cit11 | 3.1 | 3.5 | 100 | 7.3 |
| | R17 | 3.2 | 4.2 | 30 | 1.8 |
| | Cit17 | 3.4 | 4.4 | 32 | 1.6 |
| | R20 | 0.92 | 1.3 | 6.5 | 0.44 |
| | Cit20 | 1.2 | 1.6 | 9.2 | 0.51 |
| H2B tail peptide | R29 | 1.4 | 7.2 | 65 | 2.1 |
| | Cit29 | 2.2 | 11 | 96 | 2.4 |
| | R31 | 1.0 | 5.4 | 31 | 1.5 |
| | Cit31 | 2.3 | 15 | 780 | 4.9 |
| | R33 | 1.2 | 6.1 | 36 | 1.8 |
| | Cit33 | 2.2 | 11 | 110 | 2.4 |
| H3 tail peptide | T3(R2) | 7.7 | 7.5 | 660 | 51 |
| | T3(Cit2) | 8.2 | 12 | 890 | 52 |
| | R8 | 2.0 | 4.1 | 160 | 3.6 |
| | Cit8 | 1.8 | 4.6 | 170 | 3.0 |
| | R17 | 4.6 | 13.6 | 620 | 16 |
| | Cit17 | 4.7 | 15 | - | 16 |
| | R26 | 1.9 | 8.9 | 280 | 13 |
| | Cit26 | 2.2 | 11.1 | 400 | 16 |
| | R40 | 1.9 | 1.4 | 1.6 | 2.7 |
| | Cit40 | 2.9 | 2.4 | 1.5 | 3.8 |
| | R42 | 1.5 | 1.1 | 1.4 | 2.3 |
| | Cit42 | 1.6 | 1.2 | 1.5 | 2.5 |
| H4 tail peptide | R3 | 1.1 | 1.7 | 16 | 0.49 |
| | Cit3 | 1.1 | 1.7 | 18 | 0.49 |
| | R17 | - | - | - | - |
| | Cit17 | 2.4 | 10 | 140 | 7.3 |
| | R19 | 1.3 | 6.4 | 83 | 2.2 |
| | Cit19 | 2.5 | 15 | 1100 | 7.6 |
| | R23 | 1.6 | 2.3 | 5.0 | 0.29 |
| | Cit23 | 1.9 | 4.7 | 25 | 1.0 |
| H2A NCP | R3 | 0.95 | 7.4 | 27 | 0.29 |
| | Cit3 | 0.82 | 4.8 | 25 | 0.26 |
| | R11 | - | - | - | - |
| | Cit11 | 4.0 | 26 | 49 | 40 |
| H3 NCP | T3(R2) | - | - | - | - |
| | T3(Cit2) | - | - | - | - |
| | R8 | - | - | - | - |
| | Cit8 | 1.5 | 3.2 | 320 | 7.1 |
| | R17 | - | - | - | - |
| | Cit17 | 2.5 | 9.5 | 800 | 15 |
| | R26 | - | - | - | - |
| | Cit26 | 1.5 | 7.0 | 330 | 8.9 |
| H4 NCP | R3 | 0.81 | 1.5 | 12 | 0.16 |
| | Cit3 | 0.55 | 1.4 | 11 | 0.19 |
| | R17 | - | - | - | - |
| | Cit17 | 0.53 | 5.5 | 64 | 0.60 |
| | R19 | - | - | - | - |
| | Cit19 | 0.90 | 7.3 | 82 | 0.64 |

**Table S2. $t_{50\%}$ values for each PAD-histone arginine and citrulline residue.** These values correspond to **Figure 4**, and residue omissions are described in the corresponding figure legend. Calculated values are reported with two significant figures.
